# The Dominance of Geometric Graph Models in Animal Social Networks

**DOI:** 10.64898/2025.12.16.694755

**Authors:** Raima Carol Appaw, Matthew J Silk, Julie Rushmore, Kimberly VanderWaal, Michael A. Charleston, Nicholas M Fountain-Jones

## Abstract

1. Detecting patterns in animal social behaviour and movement is complicated by the diversity of ecological, evolutionary, environmental, and biological drivers of these behaviours such as migration, foraging, assortative mixing, socio-ecological factors, and human influence.
2. Addressing these complexities requires a multidisciplinary approach. Recent advances in network analysis and machine learning offer powerful tools for examining and interpreting complex network structures, aiding in the identification and quantification of movement patterns and the prediction of behavioural changes.
3. Here we use a comparative approach leveraging network analysis and machine learning techniques to assess commonalities in standard theoretical social structures governing networks across the animal kingdom. We investigate how these theoretical structures explain social organization at different scales, from entire populations to smaller groups. By leveraging interpretable machine learning techniques, we examine the predictive power of species and network construction techniques in predicting structural features of animal social networks.
4. We found that geometric graphs are the frequently predicted network model across the animal kingdom. These graphs represent both spatial and social processes and are formed by positioning individuals uniformly on a 2D plane, with links established based on proximity within a specified distance. In particular, geometric graphs demonstrate structural similarities with interaction types and data collection methods. For example, we found that this graph model had strong structural similarities with networks derived from physical contact and spatial proximity data. Networks with small-world properties, in contrast, were rare across all interaction types and collection methods. Additionally, the occurrence of these networks is influenced by the identity of species and sampling duration. Although incorporating species identity into the classification model did not improve the accuracy of the prediction, it enabled us to account for the varying dependencies of biological characteristics on specific behaviours.
5. This study highlights the value of predictive modelling for uncovering ecological drivers of animal network structures. By focusing on standard theoretical models that are well established in network science, we connect animal social networks to broader theoretical findings, while recognizing that more tailored models combining multiple generative models or network properties may offer deeper insights into network structures. We also emphasize the importance of considering how the specific methods used to build networks for each taxon could influence the biological inferences that can be drawn from those networks.

## 1 Introduction

Network theory and associated analytical techniques have been widely applied across diverse fields and real-world systems, including social science, behavioural ecology, and pathogen ecology, to investigate a broad range of processes. These include understanding animal social structures and habitat use (such as division of labour, resource allocation, and cooperation), as well as modelling the transmission dynamics of infectious diseases (Bastille-Rousseau et al., 2018; Rushmore et al., 2013; Jacoby et al., 2012; Robitaille et al., 2018; Krause, James, et al., 2015; Spiegel et al., 2016; Delmas et al., 2019; Craft, Volz, et al., 2011; Nowak et al., 1994). By leveraging network-based approaches, high-dimensional complex data in real-world systems can be transformed into a low-dimensional, simplified, and easily interpretable form where *nodes*, for example, represent individuals in population and *edges* denote interconnections or relationships between individuals (Zhang et al., 2019; Blonder et al., 2012; Newman, 2010). This simplification makes it easier to investigate complex social structural in animal communities (Beck et al., 2023). Network approaches, for example, have advanced research on cultural processes (Cantor et al., 2021), dominance hierarchies (Shizuka et al., 2012), and disease transmission (Craft, Caillaud, et al., 2011; Craft, Volz, et al., 2011; Fountain-Jones, Silk, et al., 2023).

Network generative models provide a direct approach to investigate network phenomena, that has allowed researchers to test hypotheses about network formation and evolution for a variety of species (Krause, Lusseau, et al., 2009; Romano et al., 2016; Silk & Fisher, 2017; Davis et al., 2015; Bull et al., 2012; Jacoby et al., 2012; Dijk et al., 2014; James et al., 2017). Generative models are frameworks that apply a set of fundamental principles or guidelines to create or build simulations of real-world networks (Brask et al., 2024; Keeling, Danon, et al., 2011; Keeling & Eames, 2005; Wills et al., 2020; Shirley et al., 2005; Erdös, 1960; Barabási & Albert, 1999; Barabási, 2016). These models can vary in complexity, ranging from more complex models like exponential random graph models (ERGMs) (Hunter et al., 2008) to simpler ‘standard’ models such as Erdös-Rényi(Erdös, 1960), small-world (Watts et al., 1998), and scale-free (Barabási & Albert, 1999), all of which capture different fundamental network structures. In this study, we employ these standard theoretical models that are well established in network science, to connect empirical animal networks with broader theoretical insights. For example, Erdös-Rényi random networks, created by adding edges between nodes at random (for a specific number of edges or fixed edge probability), were used to investigate the dynamics of pathogen spread, as observed in the wild Japanese macaques (Romano et al., 2016). Similarly, Watts-Strogatz small-world networks (Watts et al., 1998), formed by randomly rewiring connections in a lattice, demonstrate how high local clustering and random connections facilitate pathogen transmission, as seen in the contact networks of Serengeti lion prides (Craft, Volz, et al., 2011). Scale-free networks, formed by adding new nodes that preferentially connect to well-connected existing nodes, shows the impact of power-law distributions in examining social structures in some wildlife and livestock populations (Craft, 2015). How commonly these generative models apply across different taxonomic groups and if there are any emergent patterns across species is unknown. With the emergence of comparative social network ecology (Albery et al., 2024) and increasingly large, open-access network repositories (Collier et al., 2019), researchers now have a unique opportunity to explore broad-scale patterns of social organization across the animal kingdom.

Identifying common generative models across systems is challenging due to the multilevel nature of social organization. The model that best captures variation within animal groups may differ from those that apply at broader scales, such as across entire populations. Differences in multilevel social organization are likely widespread in nature, but have only been demonstrated in select small and large-brained species such as sharks (Mourier et al., 2019), sociable weavers (Ferreira et al., 2020), zebra finches (Brandl et al., 2021), vulturine guinea fowls (Papageorgiou et al., 2019), primates (Schreier et al., 2009), and elephants (Wittemyer et al., 2005). A notable example is found in giraffes, where social network analysis revealed multilevel organization with nested community structures (VanderWaal, Wang, et al., 2014). Building on this understanding of multilevel social organization, we examine differences in multilevel network models in four species representing different levels of social complexity and ecological niches: carpenter ants (*C. fellah*), known for their division of labor and complex social organization (Mersch et al., 2013); European badgers (*Meles meles*), which exhibit within group flexible social structures that adapt to environmental conditions (Weber, Carter, et al., 2013; Silk, Weber, et al., 2018); chimpanzees (*Pan troglodytes*), displaying human-like social hierarchies (Rushmore et al., 2013); and giraffes (*Giraffa camelopardalis*), demonstrating nested community organization (VanderWaal, Atwill, et al., 2014).

Differences in network construction methods also pose a challenge for comparative analysis of network structure of animal networks with the method employed likely impacting what structures are detected (Albery et al., 2024; De Moor et al., 2024). For example, focal sampling may miss interactions or underestimate true network structures than networks estimated from proximity data from a large number of individuals (Albery et al., 2024). Ideally, network construction methods should reflect the biological characteristics of species rather than being artifacts of specific methodology, though separating these factors often proves challenging (Albery et al., 2024).

While the standard generative models we used provide valuable insight into fundamental network structures, questions remain about their effectiveness in capturing social structural variations across the animal kingdom and the underlying biological and methodological factors driving these differences. Clarifying how data collection methods influence the resulting generative model can help improve the use of theoretical networks and make users more aware of potential biases in comparative studies.

In this comparative, we used data from the Animal Social Network Repository (ASNR) (Collier et al., 2019), to investigate the research questions outlined below. The ASNR offers a comprehensive collection of diverse datasets gathered through various methodologies and across multiple social, spatial and temporal scales for broad range of species, including insects, fish, birds, reptiles, and mammals, with variation in network size and types of behavior monitored (Rossi et al., 2015; Rossi et al., 2016). While there is some taxonomic bias, the ASNR facilitates the examination of the theoretical social structures present across the animal kingdom.

In this study, we leverage ASNR data to investigate three main questions:

i. *What common theoretical generative network models do social networks of different animal species most commonly resemble, and is this variation related to the type of social interactions or data collection methods used?*
ii. *How does resemblance to specific theoretical generative network models change across multilevel social scales, from groups to populations?*
iii. *How does species identity and network construction techniques influence the resemblance of observed animal social networks to particular theoretical generative models?*

Predicting the dominant generative model and assessing how network construction techniques (hereafter “metadata”) influence those predictions is challenging because of data complexity, nonlinear relationships, and high-order interactions that limit conventional statistical approaches. This study addresses those challenges with a powerful interpretable framework. We implement a two-stage interpretable machine learning pipeline to predict social structure and to quantify the impacts of social scale, species identity, and metadata on those predictions. Quantifying variation in network structure across species and construction methods is important for revealing more general biological and ecological principles.

## 2 Materials and Methods

We employed a range of standard theoretical network models, including Erdös-Rényi, small-world, scale-free networks, stochastic block models (SBMs), and geometric graph models. These models enabled us to investigate common social structures across the animal kingdom, accounting for different social interactions, data collection methods and taxonomic classes. These standard theoretical network models, which represent only a subset of the vast array of potential models available, are not only straightforward to simulate and analyze, but also enable us to investigate how theoretical social structures scale across different levels of networks, from small groups to entire populations. Additionally, we applied machine learning techniques to assess the relative predictive importance of species identity compared to data collection methods, social interaction types, and the duration used to observe or build networks.

### 2.1 Generative models

The generative models used in this study capture various fundamental network structures. These models are as follows:

- Erdös-Rényi (ER) networks, introduced by Paul Erdős and Alfréd Rényi in (Erdös, 1960), represent one of the most fundamental models of random networks. In this model, every possible edge between node pairs exists with a fixed probability *p*, independent of other edges (Erdös, 1960; Wills et al., 2020). Alternatively, a deterministic version fixes the number of edges (Erdös, 1960; Wills et al., 2020). The expected number of edges follows a Poisson distribution, making ER networks a useful null or baseline model for comparing real-world networks, where deviations from this random model can reveal underlying factors such as environmental constraints, trait preferences, or social hierarchies influencing connectivity patterns.
- Stochastic Block Models (SBMs) represent a random graph in which vertex sets are partitioned into *k* non-overlapping communities (where *k* = 2 in this study). In this model, edges exist independently of one another. In particular, each edge *e* = (*i, j*) has a probability *p* of existing if *i* and *j* belong to the same community, and a probability *q* if they are in different communities (Wills et al., 2020; Fathi et al., 2019; Newman, 2012; Newman, 2003). This model captures global structure without important local correlations, as the overall partitioning represents the network’s global organization or division (Wills et al., 2020). In this analysis, we utilized *k*=2 communities as a baseline to ensure simplicity in analyzing network structure and to determine whether SBM alone is sufficient to represent community structure found in real-world animal networks (Wills et al., 2020; Fathi et al., 2019; Newman, 2012; Newman, 2003). This choice aligns well with empirical observations from the ASNR data, where networks show an average module, *k* = 3 communities and an average modularity ∼0.26. See Table S9 & S10 as well as additional methods section of the electronic supplementary material for results comparing SBM network structures with different connectivity profiles and number of communities.
- The Barabási-Albert (BA) scale-free model creates networks by sequentially adding nodes and favoring connections to already well-connected nodes, a process known as “preferential attachment” (Barabási & Albert, 1999; Wills et al., 2020; Shirley et al., 2005). This model captures the degree heterogeneity often observed in real-world networks, where a small number of nodes have high connectivity, and numerous nodes with low connectivity (Barabási & Albert, 1999; Wills et al., 2020; Shirley et al., 2005). We use the BA model to explore the degree heterogeneity or variability in how connected individuals are within animal social networks.
- Real-world networks often exhibit the “small-world phenomenon,” where the average shortest path length between any two randomly selected individuals grows logarithmically as the network size increases (Watts et al., 1998; Wills et al., 2020). This means that, even in large networks, any two individuals can be connected through just a few steps (Wills et al., 2020). In the Watts-Strogatz small-world (SW) model, this phenomenon is captured by introducing random connections among nodes that are initially arranged in a regular lattice, where each node is connected to its nearest neighbors (Watts et al., 1998; Wills et al., 2020; Shirley et al., 2005). We utilized the SW model to understand how localized interactions in animal social networks can still lead to efficient information transfer across the network.

We constructed geometric graphs, a simplified form of “spatial network” model that has been widely studied across fields such as mathematics, biology, and physics (Higham et al., 2008; Boguñá et al., 2020; Barthélemy, 2011). In geometric graphs, approximately *N* nodes are distributed across a 2D (two-dimensional) space, with connections formed between pairs of nodes in the same and adjacent cells whose Euclidean distance is below a threshold radius, *r*. To promote general connectivity throughout the network, the algorithm also add edges between the nearest unconnected components. These networks capture both spatial and social processes, as the “space” can be interpreted both literally representing animal movement or abstractly as hidden or latent dimensions, with individual characteristics represented by their Euclidean distances in social or trait-based space (Janssen, Hurshman, et al., 2012; Sosa et al., 2020; Boguñá et al., 2020). Latent similarity spaces abstract dyadic social and spatial interactions into a latent dimension, allowing for the effective modeling of homophily (where similar nodes form stronger connection), particularly in the context of latent distance models in social networks (Sosa et al., 2020; Boguñá et al., 2020). Like latent models, geometric graphs often capture real-world network properties, including clustering and the small-world characteristics (Duchemin et al., 2023; Janssen, Hurshman, et al., 2012; Janssen, Prałat, et al., 2010; Higham et al., 2008; Herrmann et al., 2003; Boguñá et al., 2020; Barthélemy, 2011). We employ geometric graphs to explore how social preferences and spatial proximity influence ecological and biological processes, including mating, foraging, and cooperation in animals.

We selected these five simple models for their ability to capture the basic structural characteristics commonly observed in real-world networks. These models are both user-friendly and computationally efficient.

### 2.2 Machine learning pipeline

We implemented a two-phase classification pipeline to address the research questions, utilizing theoretical network generative models as target class labels for prediction.

#### 2.2.1 First stage classification

In the first phase, we developed a machine learning classifier utilizing the *Extreme Gradient Boosting Machine* (XGBM) algorithm, which iteratively refines its predictions by correcting errors from previous trees during training (Kuhn et al., 2022). This classifier is capable of accurately distinguishing between the standard theoretical network models, achieving an AUC close to one in prior study (Appaw et al., 2025). This classifier was trained on metrics including spectral measures (normalized Fiedler, spectral radius, eigencentrality), centrality measures (degree, betweenness, closeness centrality), as well as modularity, degree assortativity, mean path length, transitivity, mean eccentricity, minimum cut, and mean degree using the *igraph* package in R version 4.2.3 (see electronic supplementary material, Table S2 for a complete summary of the network features). These features were consistent with those used as predictors for training the classification model in a previous study by Appaw *et al*.(Appaw et al., 2025). Further details regarding the classifier, generative networks, number of simulated networks, and parameter configurations can be found in a previous work by Appaw *et al*.(Appaw et al., 2025).

#### 2.2.2 Second stage classification

In the second phase, the XGBM algorithm was employed to train two models: one including species alongside other network metadata (hereafter ‘Model 1’), and another that excluded species and used only the network metadata (hereafter ‘Model 2’). This approach allowed us to assess how the inclusion or exclusion of species influenced the overall classification outcomes.

To prepare the metadata (see electronic supplementary material, table S3 for a complete summary of the metadata) for training, we utilized categorical encoding to transform the data into a more interpretable format that the machine learning algorithms can interpret.

Both Model 1 and Model 2 were trained and tested using a 70:30 split that balanced the need for a sufficient training set to build robust models with a substantial testing set to reliably evaluate model performance. This approach was implemented within the *Tidymodel* framework in the R environment (Kuhn et al., 2022).

The second classification pipeline specifically addressed the third research question (Question 3), which was to investigate how species and the network metadata influence the resemblance of observed animal social networks to the theoretical generative models.

While there was some class imbalance in the dataset we used to train the metadata models, augmenting the minority classes or down sampling did not alter the prediction results. Moreover, XGBM have been sown to effectively handle class imbalance (Goswami et al., 2024).

### 2.3 Analyzing social structures across different methodological factors

We sourced empirical networks from the animal social network repository (ASNR) on May 3, 2023 (Collier et al., 2019; Sah, Méndez, et al., 2019). Additionally, we incorporated previously published animal network data, from studies on European badgers (*Meles meles*) (Weber, Carter, et al., 2013; Silk, Weber, et al., 2018), chimpanzees (*Pan troglodytes*) (Rushmore et al., 2013), and giraffes (*Giraffa camelopardalis*) (VanderWaal, Atwill, et al., 2014). As the first classifier was trained on binary (unweighted) networks, we did not include any edge weight information (i.e., contact frequency, duration, or intensity) from empirical datasets. We extracted metadata from these networks including species, the duration used to observe/build networks, social interactions (e.g., physical contact, grooming), data collection methods (e.g., video), and captivity status (see electronic supplementary material, Tables S2 & S3 for summaries). Networks with fewer than 10 individuals were omitted to ensure there was sufficient information for classification. For the remaining 636 networks, drawn from 33 species, we focused on the single largest connected component (group of individuals that are directly or indirectly linked) and estimated network features for each. Details on the species can be found in the supplementary material (Table S8).

To classify social structures across various animal taxa, as well as different data collection methods, types of social interactions, captivity status, and taxonomic classes (Question 1), we employed a classifier from previous studies (Appaw et al., 2025) trained using the *boosted tree* (XGBM) algorithm in the *Tidymodel* framework (Kuhn et al., 2022) within the R environment. The XGBM algorithm used data on the estimated network properties as feature vectors and the generative models as labels to train the algorithm. This method has advantages over other methods, such as random forests, as it fits trees sequentially, allowing for the correction of errors from previous iterations (Kuhn et al., 2022; Fountain-Jones, Machado, et al., 2019).

### 2.4 Analyzing social structures across group and population scales

To investigate how empirical social structures align with theoretical network models across different organizational scales, we analyzed published network data from four selected species: badgers, which exhibit flexible social groupings (Weber, Carter, et al., 2013; Silk, Weber, et al., 2018); chimpanzees (studies at both 5-meter and 50-meter interaction radii and displaying human-like social hierarchies) (Rushmore et al., 2013); giraffes, which show clear multilevel social organization (VanderWaal, Atwill, et al., 2014); and ants (Mersch et al., 2013), known for their complex division of labor and hierarchical organization within colonies. These species represent a diverse range of ecological habitats and well-documented network data as well as complex social organizational, making them ideal for comparing empirical social structures against the ensemble of fundamental theoretical models across different scales (Question 2).

The groups for each network were assembled through the following steps:

- *Group membership:* We clustered the networks using the walktrap algorithm, which is implemented using the *‘cluster_ walktrap’* function in R software version 4.2.3. This algorithm identifies densely connected groups of nodes, known as communities or clusters, within the network through random walks (Pons et al., 2005). The core principle is that short random walks tend to remain within the same community, facilitating the identification of groups (Pons et al., 2005). The *‘cluster_ walktrap’* method was specifically applied to determine the group memberships for the giraffe, chimpanzee, and ant networks. For the badger networks, we leveraged pre-existing data on ‘badger groups’ (Silk, Weber, et al., 2018) to assign group memberships to nodes within the badger networks across four distinct seasons.
- *Modularity Estimation:* We then used information on group memberships to calculate the modularity of the network in order to identify distinct communities. Modularity is typically employed alongside community detection approaches to quantify the extent to which a network can be partitioned into separate groups (Newman, 2006; Newman, 2013). A modularity score ∼0.3 and above indicates a more effective division of the network into communities (Newman, 2006; Newman, 2013; Grueter et al., 2020).
- *Group Extraction:* We processed each network in conjunction with their corresponding group membership information to extract only the largest connected components (i.e., communities or groups).
- *Generative Model Prediction:* We estimated graph features for each of the communities or groups. These features were specifically chosen for their alignment with those used to train the classification model in prior research study (Appaw et al., 2025). Subsequently, these estimated features were employed to predict the generative models most structurally similar to the social network structures within groups or communities and see if they were different to those identified at the population level.

### 2.5 Predicting social structures using machine learning approaches

To predict social structures that resemble network generative models, we employed the second phase classifier. This enabled us to assess the comparative predictive importance of species identity against the network metadata to determine how much they influence the resemblance of observed networks to theoretical frameworks (i.e., were networks constructed using mark-recapture methods more likely to be geometric). We utilized various model evaluation measures outlined below to ensure a comprehensive assessment of model performance. Additionally, we leveraged recent advancements in machine learning interpretability techniques to gain insights into the decision-making processes of the models. These methods facilitated the identification of which theoretical network models were accurately predicted when considering both species identity and network metadata (Question 3), allowing us to draw meaningful conclusions about the factors driving theoretical social structures in animal populations.

#### 2.5.1 Model interpretation

##### Shapley Additive Explanations (SHAP)

To gain insights into feature importance and interactions in the model, we used the Shapley Additive Explanations (SHAP) technique. The Shapley Additive Explanations (SHAP) technique, grounded in game theory principles, offers a perspective on feature importance for model output, aligning closely with human perceptual reasoning (Marcílio et al., 2020). SHAP assigns unique importance scores to individual features, accounting for their direct contributions to model outputs and their interactions with other features. The computation of SHAP values involves solving a set of linear equations using a specialized weighted linear regression approximation method. This approach quantifies the relative importance of each feature in the model’s final predictions. SHAP analysis provides important insights into pairwise interactions and visualizes both local and global feature importance through various tools (Marcílio et al., 2020; Lundberg et al., 2023; Štrumbelj et al., 2014).

We used the SHAP waterfall plot to display the contribution of each feature to a model’s output for a specific data instance. The SHAP feature effect or variable importance plot was used to identify the most influential model features for the global predictions of all data instances, with the most important features appearing at the top. Additionally, we used the SHAP dependency plot to show the relationship (linear or monotonic) between a feature and its impact on the model’s final prediction using scatter plots, providing an intuitive visualization of how changes in a feature affect overall predictions. More details about these plots can be found in a previous study (Appaw et al., 2025).

Detailed methodological procedures for model evaluation and analyses of interaction effects are provided in the additional methods section of the electronic supplementary material.

## 3 Results

### 3.1 Analyzing social structures across kingdoms and methodologies

Geometric graph models were the most commonly classified social network structure across animal taxa, particularly in insects, birds, and mammals (Fig. 2(a)). Geometric graphs were also most structurally comparable to particular network construction methods such as spatial proximity in reptiles and mammals, group membership in mammals and birds, and social projection bipartite associations in birds (Fig. 2 (b)). Social networks of direct interactions such as physical contact in insects and grooming in mammals were also structurally most similar to geometric graphs (Fig. 2(b)). Furthermore, behaviors like trophallaxis, dominance, and grooming in mammals, along with social projection bipartite associations in reptiles, showed structural greater similarity to to scale-free networks (Fig. 2(b)).

**Fig. 1:**
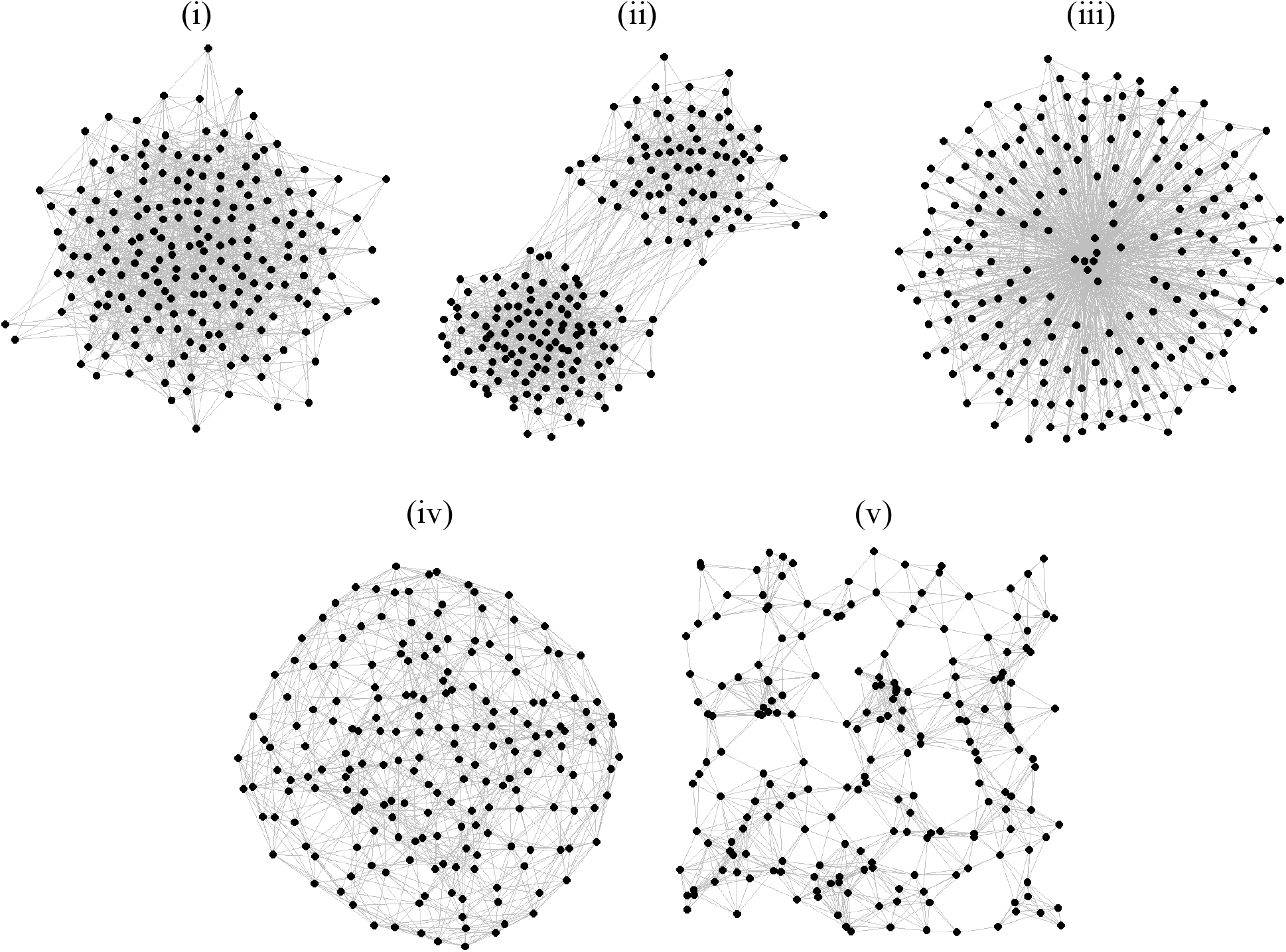
Examples of the five network generative models used in this study, each constructed with 100 nodes (*n* = 100), and roughly the same number of edges (*e* =1210): (i) Erdös-Rényi, which connects pairs of nodes with an equal probability (*p*) of 0.24; (ii) stochastic block model, composed of two blocks with sizes proportional to 40% and 60% of the total nodes (block.sizes = (0.4*n*, 0.6*n*)); (iii) scale-free, where each new node connects to 13 existing nodes (*m* = 13) based on a power-law distribution with an exponent of 2 (*power* = 2); (iv) small-world, characterized by each node being connected to its 3 nearest neighbors (*nei* = 3) and a 0.01 probability (*p*) of random rewiring; (v) geometric graphs, where nodes are connected if they fall within a threshold distance of 0.32 (*r* = 0.32).

**Fig. 2:**
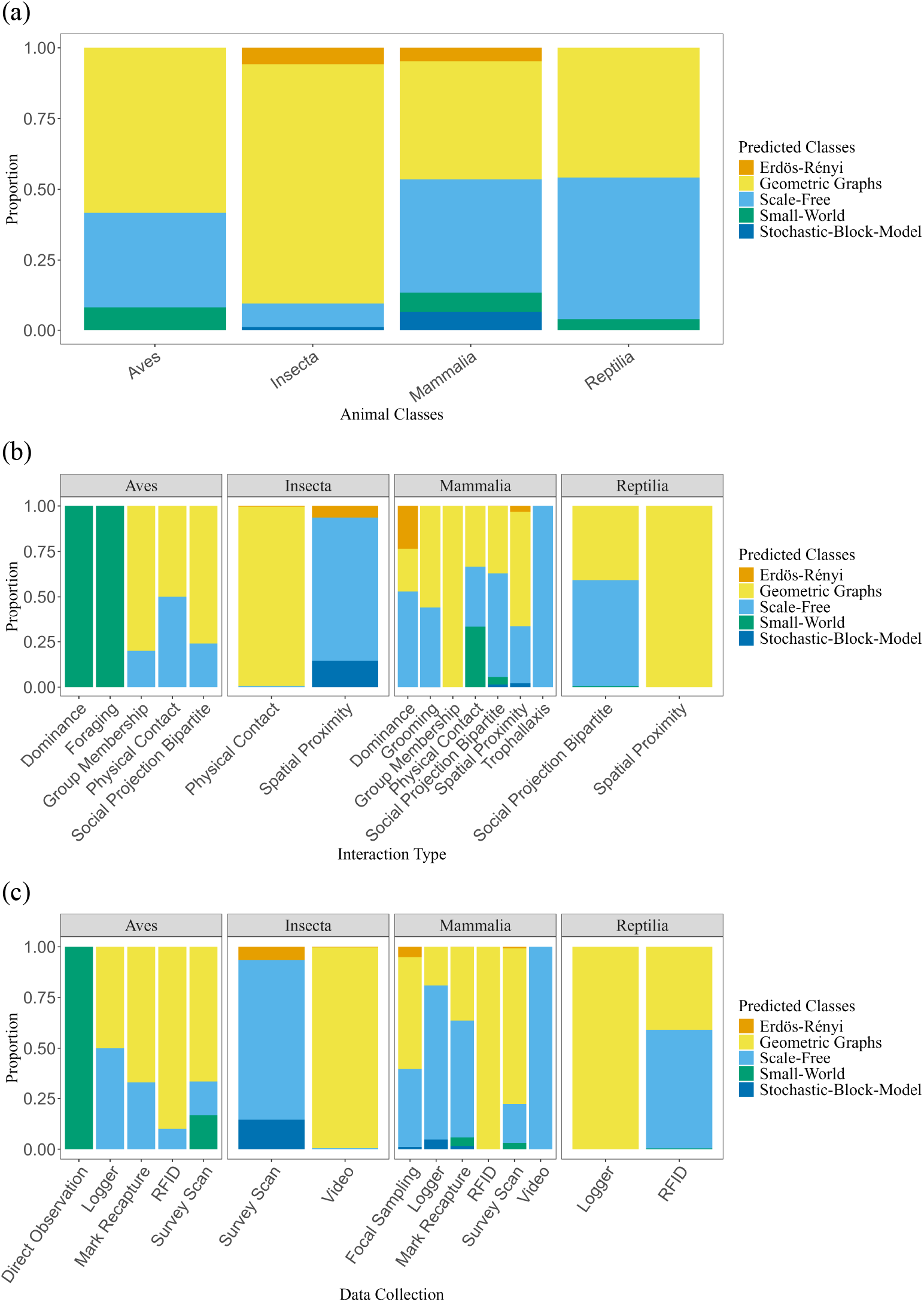
Classification of animal social networks based on (a) different animal classes, (b) interaction patterns, and (c) data collection methods. Geometric graphs were frequently observed across various taxa, especially in insects, birds, and mammals. These graphs showed structural similarities to networks derived from physical contact, spatial proximity, group membership, and social projection bipartite data. Additionally, networks obtained through data collection methods such as RFID, video recordings, survey scans, and mark-recapture techniques were often structurally similar to geometric graphs.

Social networks exhibiting structures most similar to the geometric graph were predicted across a variety of data collection methods, including video for insects, loggers for reptiles, RFID and survey scans for mammals and birds, and mark-recapture for birds (Fig. 2(c)). Direct observations in birds were structurally most similar to small-world, while survey scans in insects were structurally most similar to scale-free networks (Fig. 2(c)). In mammals, loggers and video methods showed structural similarity to scale-free networks, whereas in reptiles, RFID methods were most structurally comparable to scale-free structures (Fig. 2(c)). Additionally, networks observed in captive insects, wild birds and reptiles, as well as wild and semi-ranging mammals were also structurally similar to geometric graph models (Fig. S1).

The empirical network data obtained from the ASNR was unevenly distributed across taxonomic classes, with insects and mammals being the most represented groups. Notably, the dataset included an important number of records from *C. fellah* (a species of carpenter ants, around 250 individuals) and *M. agrestis* (voles,around 151 individuals). Nevertheless, excluding these two taxa did not alter the qualitative findings, as the predicted theoretical generative models remained consistent across multiple replicates and thus were retained in the models.

### 3.2 Analyzing social structures across group and population scales

Geometric graphs were also dominant across groups within the same population. For example, the population-level group membership social network of giraffes shows a structural resemblance to a geometric graph. The network’s modularity score was 0.26, revealing two separate large communities, both of which, similar to the overall network, exhibited structural similarities to geometric graphs (Table 1).

**Table 1:** Predicted generative models for the primary networks and groups associated with chimpanzees (May network), badgers (spring network), and giraffes. Additional results for the remaining badger networks (summer, autumn, and winter), other months of chimpanzee networks, and ant networks are provided in the electronic supplementary material.

| Main Network |  |  |  | Group Information |  |  |
| --- | --- | --- | --- | --- | --- | --- |
| Network | #Nodes | Modularity | Prediction | Groups | #Nodes | Prediction |
| Chimp_May (5m) | 32 | 0.066 | Geometric Graph | 3 | 7 | Geometric Graph |
|  |  |  |  | 1 | 16 | Geometric Graph |
|  |  |  |  | 2 | 10 | Geometric Graph |
| Badger_Spring1 | 25 | 0.544 | Geometric Graph | 5 | 3 | Geometric Graph |
|  |  |  |  | 8 | 6 | Geometric Graph |
|  |  |  |  | 1 | 9 | Geometric Graph |
|  |  |  |  | 6 | 5 | Erdős- Rényi |
|  |  |  |  | 3 | 2 | Geometric Graph |
| Badger_Spring2 | 16 | 0.289 | Geometric Graph | 5 | 3 | Geometric Graph |
|  |  |  |  | 1 | 8 | Geometric Graph |
|  |  |  |  | 6 | 2 | Geometric Graph |
|  |  |  |  | 4 | 3 | Scale-free |
| Badger_Spring3 | 14 | 0.445 | Geometric Graph | 5 | 2 | Erdős-Rényi |
|  |  |  |  | 8 | 5 | Scale-Free |
|  |  |  |  | 6 | 5 | Stochastic-Block-Model |
|  |  |  |  | 3 | 2 | Erdős-Rényi |
| Giraffe | 194 | 0.260 | Geometric Graph | 1 | 71 | Geometric Graph |
|  |  |  |  | 2 | 123 | Geometric Graph |

The European badger population-level social networks across different seasons are structurally similar to the geometric graph model (Tables 1 & S4). The modularity of the population-level badger network ranges from 0.289 to 0.701 for the eight social groups throughout the seasons (Tables 1 & S4). Although some badger groups (e.g., group 1 and 7) were consistently most similar to geometric graph structures, other groups occasionally deviated from this structure, being more similar to alternative social structures like Erdös-Rényi and scale-free in particular seasons (Table 1 & S4). Nevertheless, geometric graphs remained the most similar of the generative model set for most group-level networks as well as the population-level networks (Tables 1 & S4).

In ants, the social networks spanning five out of six colonies (colony 2-6), were structurally most similar to the Erdös-Rényi generative model (Table S7). Modularity scores ranged from 0 to 0.16. Among these networks, colony 2, 3, and 5 showed no community structure, while colony 4 and 6 exhibited two discernible groups–one large group (group 1) structurally resembling an ER network, and one very small group (group 2), classified as small-world network (Table S7). Conversely, the social network of colony 1 was structurally similar to geometric graphs and comprised three groups, mostly structurally similar to the small-world generative model (Table S7). We also found that temporal aggregation can complicate detection and characterization of social structure. For example, when networks were examined for selected days (e.g., days 5, 20, and 35) across each of the six colonies, the detected group formations were most structurally similar to geometric (spatial) graphs (Table S7).

For chimpanzees, social structures in networks based on 5 and 50-meter radii, along with their detected groups are most structurally similar to geometric graphs from the generative model set (Tables 1 & S6). The modularity of the networks across the year ranges from 0 to 0.255 with six groups detected, none of which maintained a consistent social structure over the year (Tables 1 & S6). Notably, chimpanzee social networks observed using a 5-meter radius to define proximity exhibited higher modularity and a greater number of groups compared to networks built using a 50-meter radius (Tables 1 & S6).

### 3.3 Predicting social structures using machine learning approaches

#### 3.3.1 Model Evaluation

Analysis of the theoretical generative network types showed that both metadata-based models or frameworks had the best performance in predicting geometric graphs, achieving the highest F1-score among the theoretical network structures tested (Table S1). Additionally, Model 1 (including species) achieved an overall accuracy of 0.702, while Model 2 (excluding species) had a slightly higher accuracy of 0.708 in predicting geometric graphs (Table S1). The AUC values for geometric graphs were similarly close, with Model 1 at 0.762 and Model 2 at 0.764, indicating that the inclusion of species does not contribute additional information. Scale-free networks also displayed notable precision, recall, and F1-score (Table S1). Therefore, we illustrate how the species identity and network construction methods influence the prediction of these theoretical structures.

#### 3.3.2 Model Interpretation

In the species-excluded model, the strongest predictor was data duration, accounting for ∼14% of the overall main effect in predicting the geometric graph model (Figs. 3 & S6(a)). Specifically, shorter observation periods tended to produce networks with structural similarities to geometric graphs (Fig. 4(b)). Furthermore, data collection methods explained ∼10% of the overall effect in predicting geometric graphs (Figs. 3 & S6(a)). For instance, networks generated from video recordings and RFID tracking typically showed structural similarities to geometric graphs (Fig. 4(a)). Social interactions types also contributed to ∼10% of the overall main effect in predicting geometric graphs (Figs. 3 & S6(a)), with networks formed from physical contact, and group membership showing positive SHAP values, indicating their positive influence on geometric graph structures being predicted (Fig. 4(d)).

**Fig. 3:**
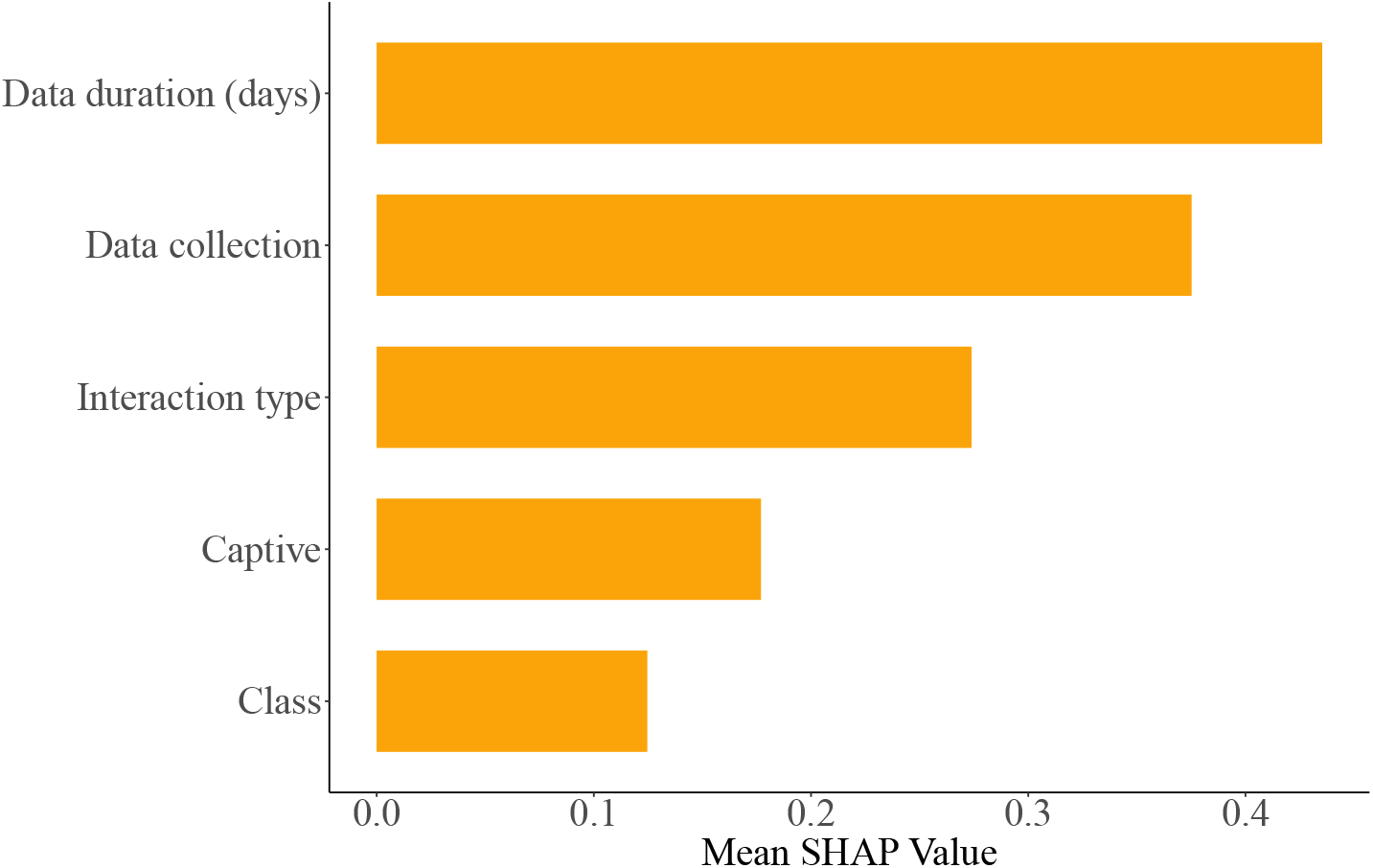
SHAP variable importance for the metadata in predicting geometric graph model for the species-excluded model is depicted by the most influential variables positioned at the top with the longest orange bars. These bars represent the variables that have the greatest impact on the model’s final output. For instance, the time span (e.g., days, weeks, years) used to construct or observe networks emerged as the most important predictor, followed closely by the data collection method and interaction type.

**Fig. 4:**
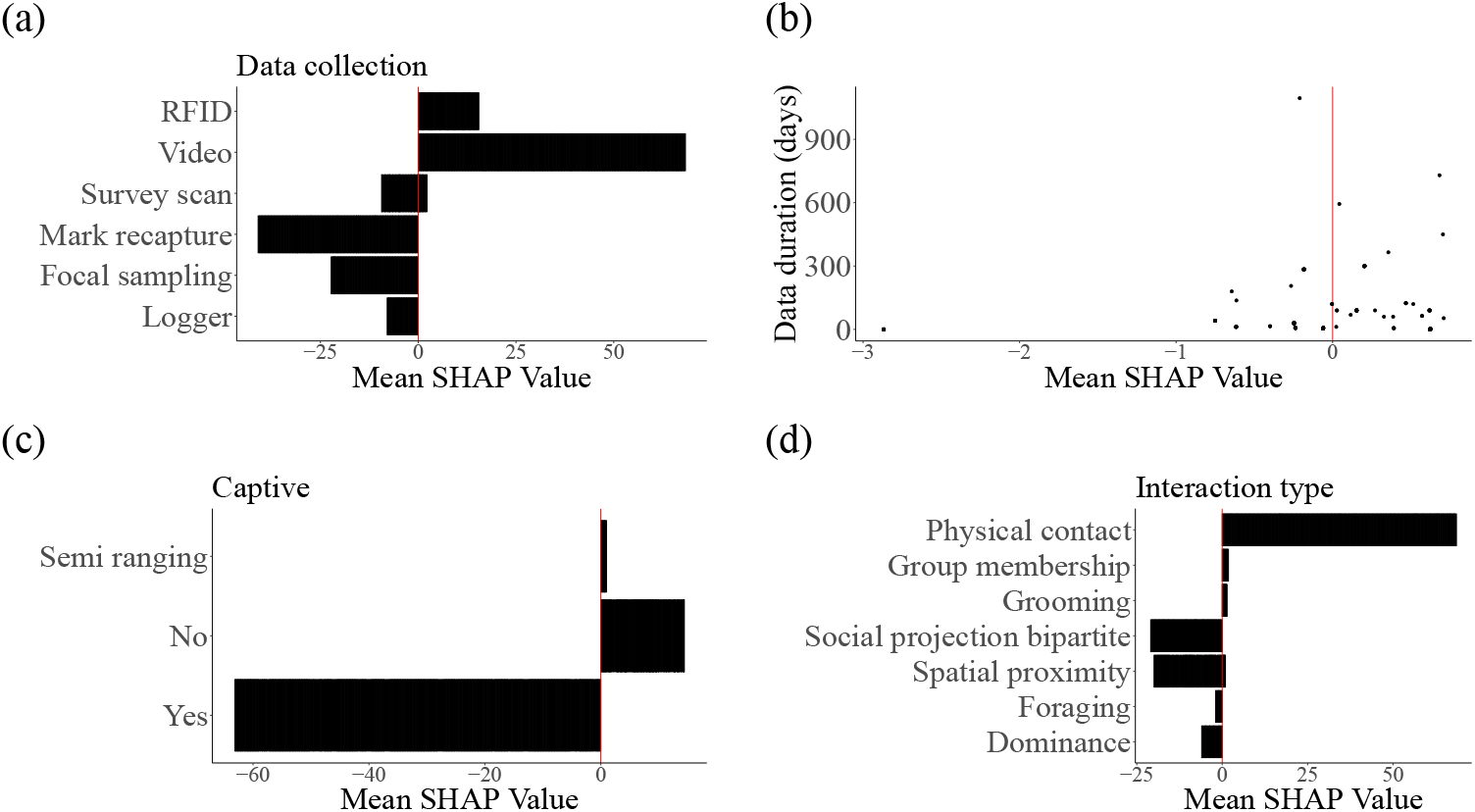
SHAP dependency plots for the species-excluded model illustrating the relationship between various network metadata and the prediction of the geometric graphs for: (a) data collection method, (b) data duration (days), (c) captive status, and (d) interaction type. The x-axis represents the SHAP value, indicating the impact of each metadata variable on the model’s prediction, while the y-axis displays the levels of these metadata. These plots offer insights into how the levels of different metadata influence the model’s final predictions. Each subplot shows how the model’s prediction changes with respect to the levels of the different metadata. For instance, the use of RFID and video methods positively impacts the prediction of geometric graphs. Additionally, interaction types such as physical contact, group membership association, and grooming networks also positively influence the prediction of geometric graphs. Similarly, shorter duration use to collect data are positively structurally most similar to predicting geometric graphs.

In contrast, networks generated from data collection methods such as mark-recapture, loggers, and focal sampling, as well as social interaction types like dominance, grooming, and social projection in bipartite networks, were positively associated with the prediction of scale-free structures (Figs. S5(a), (b), & (e)). Meanwhile, networks based on physical contact and spatial proximity in this metadata model showed no clear tendency to favour prediction of any of the five potential generative models in the species-excluded model (Figs. S5(a), (b), & (e)).

Results for the species-included model, evaluation and analyses of interaction effects are provided in the additional results section of the electronic supplementary material.

## 4 Discussion

### 4.1 Analyzing social structures across kingdoms and methodologies

This study found that the geometric graph model was consistently predicted across all behaviors, suggesting limited heterogeneity in animal social structures, which other studies supported (James et al., 2017). This highlights spatial and social preferences as key drivers of animal social structures, especially in insects and birds found in this study. In contrast, mammals and reptiles exhibited both geometric and scale-free structures, indicating heterogeneous behavior in these species, consistent with findings from research studies (Craft, 2015; Dale et al., 2017; Bull et al., 2012). For instance, desert tortoise social networks, tracked through radio telemetry, associated with spatial distance (Sah, Nussear, et al., 2016; Niblick et al., 1994; Bull et al., 2012) –closer to the geometric model–and scale-free behaviors observed in some livestock and wildlife populations (Craft, 2015).

The social structures of very few animal species resemble the Erdös-Rényi (ER) random graph model (Wills et al., 2020; Pržulj, 2007; Brask et al., 2024), and the results in this study support the fact that it is unsuitable for capturing real-world properties. While the ER model is sometimes used as a null model in comparative studies (Wandelt et al., 2019; Sah, Nussear, et al., 2016; Wills et al., 2020), it may only be applicable in specific scenarios, such as studying behavioural patterns in highly fluid social groups like schools of fish that often move in unpredictable directions (Reuter et al., 2016; Couzin, Krause, et al., 2005; Couzin, Krause, et al., 2003).

The small-world network, characterized by high local clustering with short paths, have proven valuable in studying pathogen transmission in animals, such as in the contact networks of Serengeti lion (Craft, Volz, et al., 2011). Similarly, studies on the social structures of leaf-roosting bats *T. tricolor* have also demonstrated small-world properties (Chaverri, 2010). However, the results in this study suggest that small-world networks are relatively rare in nature, suggesting that geometric graph model or extended versions may be better suited to incorporate long-distance connections.

This study also found limited evidence that stochastic-block-models (SBMs) alone capture social structures more accurately than other models we considered. SBMs used to analyze community structures incorporate explicit communities with higher edge probabilities within groups (Newman, 2012; Wills et al., 2020), potentially being adapted to ‘spatial’ networks like geometric graphs if connections depend on node distance (Barthélemy, 2011). However, this similarity makes it challenging to distinguish SBMs from geometric graphs. Nevertheless, when preferences are based on a mix of spatial and social factors, geometric graphs may be favoured and still generate networks with clear community structures.

In this study, we found that data collection methods such as RFID tracking, video, and observing interactions via spatial proximity, direct physical contact and group membership produce networks classified as geometric graphs. This suggests that both spatial and social processes which emphasize encounters within shared spaces and behaviors or preferences based on traits, and shared environment are primary drivers of animal social networks. For instance, it has been demonstrated that ‘spatial models’ (based on home range and movement patterns) outperform scale-free or Erdős–Rényi (ER) models in predicting animal social contact networks (James et al., 2017). However, data collection methods like focal sampling, and mark recapture alongside behaviors like grooming, and dominance, align more with networks like scale-free. For instance, network data on dog social interactions collected using focal sampling illustrate that some higher-ranking, more dominant dogs engage in more aggressive interactions (Silk, Cant, et al., 2019). Similarly, animals involved in grooming preferentially interact with high-ranking or socially dominant individuals (Schino, 2001).

### 4.2 Analyzing social structures across group and population scales

We demonstrate that geometric graph models explain theoretical social structures across various multilevel network scales, from groups to entire populations. This suggests that both spatial and social factors primarily drive multilevel social organizations (Grueter et al., 2020). The association patterns observed at the multilevel societies provide insights into the micro-dynamics of social structure, revealing specific biological information that informs behavioural and ecological processes within smaller social units. For instance, hamadryas baboons’ (Papio hamadryas) foraging patterns are linked to sparse resource distribution (Schreier et al., 2009), while vulturine guinea fowl show preferential associations during social activities (Papageorgiou et al., 2019). Although geometric graphs are common across scales, the depth and complexity of social interactions may vary temporally, with more nuanced patterns emerging at the group level (Papageorgiou et al., 2019; Grueter et al., 2020). These patterns may evolve over time and be less apparent at population scales. This highlights the importance of considering hierarchical multilevel networks in examining social organizations, as each scale offers unique perspectives on the dynamics of social networks.

For example, we found that social structures in giraffes across group and population level are primarily shaped by spatial and social behaviors, represented by geometric graphs. This is consistent with studies that highlighted that spatial networks enhance sociality in this species, especially as individuals with overlapping home ranges tend to form strong associations or social bonds, facilitating pathogen spread (VanderWaal, Atwill, et al., 2014; VanderWaal, Wang, et al., 2014).

Badgers exhibit the strongest community structure among the animals we investigated, especially across seasons, with network structures at both population and group level typically resembling the geometric graph model. The geometric graph models model may be more appropriate to describe badger social structure, due to the spatial proximity of setts, with its importance for group-level structure perhaps explained by either spatial patterns within social communities caused by some social grouping incorporating multiple setts (Silk, Weber, et al., 2018; Weber, Bearhop, et al., 2013) or potentially also social preferences within groups. Additionally, seasonal shifts, particularly in spring, and temporal changes reveal flexibility within these social structures, consistent with previous findings (Silk, Weber, et al., 2018; Weber, Bearhop, et al., 2013).

In the 41 daily interaction networks from six ant colonies, three showed no detectable modularity, while the other three exhibited weak structure with up to three groups. These networks resembled small-world or Erdös-Rényi forms, contrasting with the stronger, more stable groupings reported by Mersch et al., 2013. These differences likely reflects methodological and analytical differences. We applied modularity optimization, which detects densely connected clusters but can miss weak or overlapping associations (Newman, 2006), whereas Mersch et al., 2013 used Infomap algorithm, a random-walk based method more sensitive to subtle interaction flows. Additionally, their analysis aggregated interactions into four multi-day periods (11, 10, 10, and 10 days), whereas our approach examined colony-level networks across all 41 days, which may have resulted in fewer detectable modules. We also found that temporal aggregation of networks can complicate accurate characterization of social structures across scales. For instance, further analysis of social structures on some selected days for the six colonies detected groups structurally similar to geometric graph (that is, spatial network), aligning with the findings of Mersch *et al*.(Mersch et al., 2013). Future work could explore alternative community detection methods or temporal aggregation schemes to assess whether stable subgroups emerge under different resolutions.

In this study, wild chimpanzees displayed community structures, consistent with previous works (Rushmore et al., 2013; Schreier et al., 2009; Grueter et al., 2020; Papageorgiou et al., 2019). Wild chimpanzees have been shown to live in communities that split into smaller, sub-communities with varying sizes (Rushmore et al., 2013). Social structures across the multilevel communities in this species resemble the geometric graph model, likely due to association observed across various spatial levels (i.e., 5-meters and 50-meters). Furthermore, social factors, such as family connections and group size, influence chimpanzee associations (Rushmore et al., 2013), reinforcing why social structures resemble the geometric graph model.

### 4.3 Predicting social structures using machine learning approaches

Network models incorporating spatial and social architecture, as represented by the unweighted geometric graph approach, better reflect real-world animal social structures compared to random or degree-based models, aligning with previous findings (James et al., 2017; Kappeler et al., 2015). Both metadata analyses reveal that spatial and social behaviors influence the predictive power of theoretical network models. For example, possum home-range dynamics were best approximated by space-utilization model, which outperformed conventional null models such as Erdös-Rényiand scale-free networks in explaining empirical contact patterns (James et al., 2017).

In this study, networks observed via behaviors such as physical contact, proximity and group membership was predicted to have geometric graphs structure, whilst dominance, and grooming behaviors were predicted to have scale-free structures. Furthermore, data collection methods also impact the prediction of the network structures (De Moor et al., 2024; Albery et al., 2024; Spiegel et al., 2016). For example, if network construction is based on spatial proximity, employing continuous focal sampling—which involves observing specific individuals over time—rather than using proximity loggers may misinform the true underlying network structure (Albery et al., 2024; De Moor et al., 2024). In this study, networks generated using methods such as focal sampling, mark-recapture, and loggers are more likely to reveal scale-free structures, while RFID and video monitoring are associated with predicting geometric graph-like structures.

The findings of this research study align with studies that underscore the impact of network observation periods on network structures (Farine et al., 2015; Franks et al., 2021; Delmas et al., 2019; De Moor et al., 2024). Although networks observed over extended periods may reflect real-world patterns (De Moor et al., 2024; Farine et al., 2015; Franks et al., 2021), they could mask the true network dynamics by capturing rare and random connections (Albery et al., 2024). Overall, to improve prediction accuracy, incorporating a more comprehensive range of metadata, including species identity and individual-level data, along with increased repetition across metadata types, could be beneficial.

This study underscores the importance of considering how networks are built and the influence that methodological factors have on predicting the network structures. Establishing a consistent criteria or precedence for selecting methodological factors in network structure analysis, and refraining from comparing networks built or observed by differing approaches can help maintain accuracy and provide a better insight (Albery et al., 2024). See Albery *et al*.(Albery et al., 2024), and De moor *et al*.(De Moor et al., 2024) for a comprehensive discussion on the complexities of accounting for the confounding effects of methodological factors in network structure analysis.

#### Limitations

Despite providing valuable insights, this study has some important limitations. While we chose to use unweighted networks to avoid the inconsistencies and complications that arise from varying edge weight interpretations across different network models, this choice limited the ability to examine the nuances associated with weighted edges (Brask et al., 2024; De Moor et al., 2024; Sosa et al., 2020). Weighted networks, which can represent frequencies, durations, counts of interaction, are important for understanding the varying strengths of interactions (De Moor et al., 2024). The use of unweighted networks, while widely adopted due to its analytical simplicity and value as a starting point for analysis (Brask et al., 2024), carries important limitations. For instance, social dynamics change over time, so in studies with extended data collection, neglecting interaction strength can result in oversimplified interpretations and potentially inaccurate conclusions (Albery et al., 2024; De Moor et al., 2024).

Another limitation of this study is that we did not explore other, more flexible generative models, as the evaluation was confined to only five standard theoretical models. While these models serve as a sensible starting point, many additional rules from behavioural ecology could be applied. For instance, the social inheritance model builds on an established social science model for network growth and describes how individuals form social connections, with bolder individuals more likely to connect outside their parents’ social circle while others tend to inherit their parents’ social connections (Ilany et al., 2016). We found that behaviours like proximity, direct physical contact, and group membership are inherently tied to both spatial and social factors, as represented by the geometric graph model. This suggests that methods that can disentangle both models are needed. In contrast, behaviours like dominance and grooming involve decision-making like degree heterogeneity and popularity, characteristics of scale-free structures. Furthermore, geometric graphs often overlap with other models; for instance, Stochastic-Block-Models can be adapted to spatial networks when connection probabilities are distance-dependent (Barthélemy, 2011). These insights suggest that employing multiple interacting generative models that produce networks representing varying levels of spatial and social preferences, and capturing different structures, may provide a more nuanced understanding of network structures. For example, the *GenNetDem* R package (Silk & Gimenez, 2023), integrates features of a simple two-dimensional latent space model and a stochastic block model to generate underlying social networks.

## 5 Conclusion

This research study underscores the important role of spatial and social processes in explaining theoretical generative models in the animal taxa we observed. Generally, results in this study suggest that geometric graph models more accurately represent the basic social structures in animal networks compared with other conventional theoretical network models like scale-free networks. Furthermore, this research study illustrates how taxonomic biases, species identity, and methodological factors can impact network construction and interpretation. This underscores the necessity of incorporating multiple interacting generative models, and a comprehensive range of metadata, including species identity and individual-level data, as well as increasing the repetition across metadata types and network methods to enhance predictive accuracy. Future research will focus on exploring how various species and their traits correspond to different generative models for animal social networks. This study provides a foundation for employing advanced machine learning techniques to test different possible explanations for theoretical network structure, an approach that has the potential to enhance the macroecological understanding of animal social behaviour.

## 6 Authors’ contributions

R.C.A contributed to the idea, developed the model training code, conducted the analysis, and wrote the paper. M.J.S. contributed to the conceptualization of the idea, provided data, and engaged in methodology, review, and editing. J.R. provided data, methodology, review and editing. K.V. provided data, methodology, review and editing. M.C. contributed to the conceptualization of the idea, methodology, review and editing. N.F.J. conceived the idea, methodology, review and editing. All authors provided final approval for publication.

## 7 Acknowledgements

This project was supported by an Australian Research Council Discovery Project Grant (DP190102020).

## 8 Data statement

All the data and code to perform the analysis can be found at https://github.com/araimacarol/Dominance-of-Geometric-Graph-in-Animal-Social-Networks

