## Supplementary Material for "The Dominance of Geometric Graph Models in Animal Social Networks"

### 1 Supplementary Material

#### 1.1 Additional Methods

##### 1.1.1 Generative models

**Stochastic Block Models (SBMs)**

To evaluate our classifier's performance across different community structures, we conducted simulations of SBMs varying both the number of communities ($k$ = 2,3,4,5,6) and network sizes (100-1000 nodes). These simulations tested two scenarios of between-group connectivity: sparse ($0.01 \leq q \leq 0.04$ ) and dense ($0.1 \leq q \leq 0.4$ ), while maintaining consistent within-group connectivity ($0.5 \leq p \leq 0.8$) (Table S9 & S10). The simulated SBMs with dense between-group connectivity had an average modularity (~ 0.22) closely matching that observed in empirical Animal Social Network Repository (ASNR) data, while sparse connectivity resulted in much higher modularity (0.608). This correspondence between the SBMs simulated with the dense between-group probability and empirical networks suggests that our classifier with the SBM network simulated with $k = 2$ communities, employing similar connectivity parameters, captures fundamental aspects of community structure found in the ASNR. Future studies should however, focus on training a classifier across ranges of density and modularity values and assess predictive performance on empirical networks.

##### 1.1.2 Model evaluation

In evaluating the machine learning model, we employ combinations of these metrics to evaluate the performance of the network metadata and species identity in predicting the theoretical networks across the two metadata machine learning classification models. These metrics are described below.

**Recall**

Recall, also known as sensitivity or true positive rate, is the ratio of correctly predicted positive observations to the total actual positives (Yacouby & Axman 2020; Sokolova & Lapalme 2009; Powers 2020). It is calculated as:

$$\text{Recall} = \frac{\text{True Positives}}{\text{True Positives} + \text{False Negatives}}$$

Recall is a is threshold-dependent metric, as it varies with different classification thresholds (Yacouby & Axman 2020).

**Precision**

Precision, or positive predictive value, is the ratio of correctly predicted positive observations to the total predicted positives (Yacouby & Axman 2020; Sokolova & Lapalme 2009; Powers 2020). It is defined as:

$$\text{Precision} = \frac{\text{True Positives}}{\text{True Positives} + \text{False Positives}}$$

Like recall, precision is threshold-dependent metric (Yacouby & Axman 2020) .

##### Accuracy

Accuracy is the ratio of correctly predicted observations (both true positives and true negatives) to the total observations (Yacouby & Axman 2020; Sokolova & Lapalme 2009; Powers 2020). It is given by:

$$\text{Accuracy} = \frac{\text{True Positives} + \text{True Negatives}}{\text{Total Observations}}$$

Accuracy is also threshold-dependent metric (Yacouby & Axman 2020).

##### F1-score

The F1-score is the harmonic mean of precision and recall. It provides a balance between precision and recall, especially when the class distribution is imbalanced (Yacouby & Axman 2020; Sokolova & Lapalme 2009; Powers 2020). It is calculated as:

$$\text{F1-score} = 2 \times \frac{\text{Precision} \times \text{Recall}}{\text{Precision} + \text{Recall}}$$

##### ROC Curve and AUC

The Receiver Operating Characteristic (ROC) curve is a graphical plot that illustrates the diagnostic ability of a binary classifier system as its discrimination threshold is varied (Yacouby & Axman 2020; Sokolova & Lapalme 2009; Powers 2020). The Area Under the ROC Curve (AUC-ROC) provides an aggregate measure of performance across all possible classification thresholds. ROC is a rank-based metric (Yacouby & Axman 2020).

##### 1.1.3 Model interpretation (Interaction effect)

To identify variables engaged in interactions, and to measure the strength and importance of feature interactions and their influence on model predictions, we integrated Friedman and Popescu's H-statistics into our analysis (Friedman & Popescu 2008). These statistics serve as additional tools for evaluating the performance of machine learning algorithms in both classification and regression tasks. Specifically, $H^2$ quantifies the proportion of prediction variability attributed to interactions among features, highlighting how much of the variability is explained beyond the main effects alone. This metric is important because it reflects the importance of feature interactions in capturing the complexities of the data. Additionally, the overall interaction effect measures how individual features contribute to variability when considering their interactions with other features. Pairwise and three-way interactions further assess how combinations of two or three features together influence the model. Our computations of these statistics follow methodologies detailed in a GitHub resource at https://github.com/mayer79/hstats#background. We also employed the SHAP 2D dependency plot to examine the nature or shape of feature interactions and visualize how the combined effects of two features influence the model's final predictions (Marcílio & Eler 2020; Lundberg et al., 2023; Štrumbelj & Kononenko 2014).

By incorporating these advanced statistical techniques, we gain a clearer picture of the interactions between features and their contribution to model performance. This approach not only complements SHAP values but also reveals how interactions between features provide a richer and more nuanced understanding of the model predictions.

#### 1.2 Additional Results

##### 1.2.1 Model Evaluation

Table S1: Confusion matrices depicting the classification performance of our pipeline for (a) Model 1 (species included) and (b) Model 2 (species excluded) using metadata test data. Each matrix contrasts predicted and actual class labels, offering detailed counts of true positives, true negatives, false positives, and false negatives across all classes. Furthermore, precision, recall, and F1-score metrics are provided for each class, providing insights into the models' accuracy in identifying the true positives, true negatives, false positives, and false negatives. The table also includes $H^2$ (Friedman & Popescu 2008), which measures the proportion of predicted variability that cannot be explained by main feature effects alone.

(a) Model 1

| | Erdös-Rényi | SBM | Scale-Free | Geometric Graph | Small-World | Recall | $H^2$ |
|---|---|---|---|---|---|---|---|
| Erdös-Rényi | 2 | 0 | 6 | 1 | 0 | 0.22 | 0.58 |
| SBM | 1 | 0 | 5 | 1 | 0 | 0 | 0.45 |
| Scale-Free | 5 | 0 | 34 | 10 | 0 | 0.694 | 0.97 |
| Geometric Graph | 0 | 0 | 19 | 97 | 0 | 0.836 | 0.67 |
| Small-World | 0 | 0 | 8 | 1 | 1 | 0.1 | 0.38 |
| Precision | 0.25 | 0 | 0.472 | 0.882 | 1 | | |
| F1-score | 0.234 | 0 | 0.562 | 0.860 | 0.182 | | |

(b) Model 2

| | Erdös-Rényi | SBM | Scale-Free | Geometric Graph | Small- World | Recall | $H^2$ |
|---|---|---|---|---|---|---|---|
| Erdös-Rényi | 0 | 0 | 4 | 0 | 0 | 0 | 0.74 |
| SBM | 0 | 0 | 7 | 2 | 0 | 0 | 0.16 |
| Scale-Free | 0 | 0 | 35 | 20 | 1 | 0.625 | 0.27 |
| Geometric Graph | 0 | 0 | 14 | 101 | 0 | 0.871 | 0.24 |
| Small-World | 0 | 0 | 7 | 1 | 0 | 0 | 0.60 |
| Precision | 0 | 0 | 0.522 | 0.815 | 0 | | |
| F1-score | 0 | 0 | 0.569 | 0.842 | 0 | | |

##### 1.2.2 Model Interpretation

**Species-included model**

In the species-included model, species identity was the strongest predictor, followed closely by the duration of network observations, data collection methods, social interaction types, and captive status (Fig. S3(a)). Specifically, networks measured over shorter observation periods, for small to medium-sized species — such as captive ants interacting via physical contacts and monitored via video and wild badgers tracked with RFID interacting via spatial proximity — were more likely to show structural similarities to geometric graphs relative to our other model types (Figs. S3(b), (c), (d), (e) & (f)).

Conversely, networks derived from data collection methods like mark-recapture, loggers, and focal sampling, along with social interactions such as dominance, grooming, and social projection in bipartite networks, exhibited greater structural similarity to scale-free networks compared with our other model types, but lacked resemblance to geometric graphs in the species-included model (Figs. S4 (a), (b), (c) & (f)). Results on the species-excluded model can be obtain in the additional results section of the electronic supplementary material.

**Interaction effect**

For the metadata model without species identity, we found that ~24% of the prediction variability ($H^2$) for the geometric graph model was unexplained by the main effects alone (Table S1 (b)). The most important two-way interaction predicting structures similar to geometric graphs was between data duration and captivity status, which accounted for ~8.5% more of joint effect probability compared with the individual main effects (Fig. S6 (b)). Additionally, the interactions between data duration and data collection methods, data duration and taxonomic classes, and data duration and interaction type explained ~7%, ~6.8%, and ~6.7% more of the joint effect probability, respectively, than the main effects alone (Fig. S6 (b)).

SHAP analysis plotting the interaction between data duration and data collection methods reveals that geometric graph structures are more likely to be predicted when shorter observation periods are paired with RFID tracking and video monitoring methods (Fig. S7 (b)). Similarly, SHAP analysis plotting the interaction between data duration and captivity status shows a higher likelihood of predicting geometric graphs when non-captive and semi ranging animals are observed over shorter periods (Fig. S7 (c)). Furthermore, SHAP plots illustrating the interaction between data duration and interaction type suggest that geometric graph structures are more likely to be predicted when data on direct physical contact, close spatial proximity, group membership, and grooming networks are collected over shorter time frames (Fig. S7 (a)). SHAP plots showing the interaction between data collection methods and interaction types reveal that direct physical contact observed via video, RFID monitoring of animals in close spatial proximity, and survey scans of group membership are key in predicting geometric graph-like network structures (Fig. S7 (d)).

In the species-excluded model, the interaction patterns identified in predicting geometric graphs closely mirror those found in the species-inclusive model (Figs. S7 & S8). Additionally, the interaction patterns associated with network construction methods for predicting scale-free networks showed comparable trends in both the species-included and species-excluded models (Figs. S9 & S10).

Table S2: Definitions of graph features used as feature vectors in training the machine learning classification model used to classify networks.

| Features | Definition |
| --- | --- |
| Fiedler Value | The second smallest non-trivial eigenvalue of the laplacian matrix (Seary & Richards 2003; Wills & Meyer 2020). |
| Normalized Fiedler Value | Measures the connectivity of a graph by normalizing the Fiedler value, which is the second smallest eigenvalue of the Laplacian matrix (Seary & Richards 2003; Wills & Meyer 2020). |
| Spectral Radius | The largest eigenvalue of the adjacency matrix, representing the overall connectivity strength in a network (Wills & Meyer 2020; Seary & Richards 2003). |
| Eigenvector Centrality | This measure centralizes a graph according to the eigenvector centrality of nodes (Freeman et al., 2002). Eigen centrality considers not only the number of direct connections a node has (like degree centrality) but also the "influence" of the neighbouring nodes in the graph (Bonner et al., 2016). |
| Degree Centrality | Measures how the degrees of nodes in a graph differ from the degree of the most central node (Freeman et al., 2002; Wasserman & Faust 1994). Nodes with high degrees assume crucial central roles in graphs and are important for the overall network functionality (Kajdanowicz & Morzy 2016). |
| Betweenness Centrality | This is defined as the number of geodesic distances (shortest path) traversing a node. (Kajdanowicz & Morzy 2016; Freeman et al., 2002; Wasserman & Faust 1994). |
| Closeness Centrality | Measures the average shortest path length from a node to every other node in the network (a measure of the number of steps to access every other vertex from a given vertex) (Kajdanowicz & Morzy 2016; Freeman et al., 2002; Wasserman & Faust 1994). |

| | |
|---|---|
| Modularity | A measure of the strength of division of a network into modules (communities), indicating the degree of clustering (M. E. Newman 2013; Sah et al., 2017; M. E. J. Newman 2003). |
| Degree Assortativity | A measure of the level of homophily, thus, the tendency of nodes to connect with other nodes with similar attributes or degree (M. E. J. Newman 2003; Carnegie 2018). |
| Mean Path Length | The *mean path length* (or mean geodesic distance) of a graph measures the average number of steps traverse from a node $v_i$ to each other node in a graph (Li et al., 2012; Harary 2018; Keeling et al., 2011; M. Newman 2010). |
| Transitivity (Clustering Coefficient) | The *clustering coefficient* is defined as the proportion of a node's neighbors that are also neighbors of each other (Wills & Meyer 2020). |
| Mean Eccentricity | The *eccentricity* of a node is the maximum distance from that node to any other node in the graph (Li et al., 2012; Harary 2018; Bondy 1976). Therefore, the *mean eccentricity* of a graph is the average eccentricity of all nodes in a connected graph (Li et al., 2012; Harary 2018; Bondy 1976), |
| Minimum Cut | A *minimum cut size* of a graph is the minimum total number of edges required to disconnect the network into more connected components (at least two components) and is a coarse measure of the connectedness of the network (Stoer & Wagner 1997; Seary & Richards 2003). |
| Mean Degree | The mean degree of a graph is defined as the average number of all the node degrees of the graph (Shirley & Rushton 2005; Bondy 1976). |
| Graph energy | The *graph energy* is defined as the squared sum of the absolute values of all eigenvalues of the adjacency matrix (Li et al., 2012; Harary 2018). |

Table S3: Network metadata recorded for each network.

| Metadata | Description | Type |
|---|---|---|
| Host species | Host species and taxonomic class that the network data came from | Factor |
| Interaction type | Do edges reflect physical contact, dominance, spatial proximity, grooming, foraging, or group membership? | Factor |
| Data collection method | Was the data collected using a video, focal sampling, logger, survey scan, or other method? | Factor |
| Network duration | Over what period was the data collected or observed (days) | Numeric |
| Captive/free (wild) | Was the network recorded for captive or free-ranging populations? | Factor |

Table S4: Definition for types of Animal social interactions

| Interaction Type | Definition |
|---|---|
| Physical Contact | Direct physical touch or interaction between individuals (Rocha et al., 2021). |
| Spatial Proximity | The closeness of individuals in space (geographic location or distance), without necessarily involving direct contact, from which one can infer their relationships or potential interactions (Boyland et al., 2013). |
| Grooming | Social behavior where one individual cleans or maintains another, often serving to cement and strengthen affiliative relationships, but can also be exchanged for social goods and services (Crofoot et al., 2011). |
| Social Projection Bipartite | Social projection Bipartite (SPB) networks transform a bipartite network of individuals and entities (e.g., birds and nest chambers) into a unipartite network (all nodes belong to a single class, and connections (edges) occur between any pair of nodes in that class). In an SPB networks, individuals are connected if they share the same entity (e.g., nest chamber) (Dijk et al., 2014). |
| Group Membership | The state of belonging to a specific social group (where groups are defined through spatial or temporal proximity) (Croft et al., 2011). |
| Foraging | Behavior related to the search and acquisition of resources like food or water, which often involve collaboration or competition to survive and reproduce (Barack 2024). |
| Dominance | Social hierarchy that determines access to resources or mating opportunities, often involving aggressive interactions (Shizuka & McDonald 2012). |

Table S5: Predicted generative models for the badger networks and groups for autumn and winter.

| Main Network | | | | Group Information | | |
|---|---|---|---|---|---|---|
| Network | #Nodes | Modularity | Prediction | Groups | #Nodes | Prediction |
| Badger_Spring1 | 25 | 0.544 | Geometric Graph | 5 | 3 | Geometric Graph |
| | | | | 8 | 6 | Geometric Graph |
| | | | | 1 | 9 | Geometric Graph |
| | | | | 6 | 5 | Erdös- Rényi |
| | | | | 3 | 2 | Geometric Graph |
| Badger_Spring2 | 16 | 0.289 | Geometric Graph | 5 | 3 | Geometric Graph |
| | | | | 1 | 8 | Geometric Graph |
| | | | | 6 | 2 | Geometric Graph |
| | | | | 4 | 3 | Scale-Free |
| Badger_Spring3 | 14 | 0.445 | Geometric Graph | 5 | 2 | Erdös- Rényi |
| | | | | 8 | 5 | Scale-Free |
| | | | | 6 | 5 | Stochastic-Block-Model |
| | | | | 3 | 2 | Erdös- Rényi |
| Badger_Autumn1 | 20 | 0.504 | Geometric Graph | 1 | 8 | Geometric Graph |
| | | | | 6 | 8 | Scale-Free |
| Badger_Autumn2 | 20 | 0.504 | Geometric Graph | 1 | 8 | Geometric Graph |
| | | | | 6 | 8 | Scale-Free |
| Badger_Autumn3 | 26 | 0.478 | Geometric Graph | 5 | 3 | Geometric Graph |
| | | | | 8 | 8 | Geometric Graph |
| | | | | 6 | 9 | Geometric Graph |
| | | | | 2 | 3 | Scale-Free |
| Badger_Autumn4 | 41 | 0.701 | Geometric Graph | 5 | 3 | Geometric Graph |
| | | | | 8 | 8 | Geometric Graph |
| | | | | 1 | 9 | Geometric Graph |
| | | | | 6 | 9 | Geometric Graph |
| | | | | 7 | 4 | Geometric Graph |
| | | | | 4 | 5 | Geometric Graph |

| Main Network | | | | Group Information | | |
|---|---|---|---|---|---|---|
| Network | #Nodes | Modularity | Prediction | Groups | #Nodes | Prediction |
| Badger_Winter2 | 20 | 0.461 | Geometric Graph | 5 | 3 | Geometric Graph |
| | | | | 1 | 9 | Geometric Graph |
| | | | | 2 | 2 | Geometric Graph |
| | | | | 4 | 5 | Geometric Graph |
| Badger_Winter3 | 34 | 0.556 | Geometric Graph | 5 | 3 | Geometric Graph |
| | | | | 1 | 9 | Geometric Graph |
| | | | | 3 | 2 | Geometric Graph |
| | | | | 4 | 5 | Geometric Graph |
| Badger_Winter4 | 20 | 0.541 | Geometric Graph | 5 | 6 | Geometric Graph |
| | | | | 8 | 9 | Scale-Free |
| | | | | 1 | 9 | Geometric Graph |
| | | | | 3 | 9 | Scale-Free |
| | | | | 4 | 9 | Erdös- Rényi |
| Badger_ Summer1 | 38 | 0.654 | Geometric Graph | 5 | 3 | Geometric Graph |
| | | | | 8 | 5 | Scale-Free |
| | | | | 3 | 5 | Erdös- Rényi |
| | | | | 1 | 9 | Geometric Graph |
| | | | | 6 | 9 | Geometric Graph |
| | | | | 2 | 3 | Geometric Graph |
| | | | | 4 | 4 | Geometric Graph |
| Badger_ Summer2 | 24 | 0.429 | Geometric Graph | 5 | 3 | Geometric Graph |
| | | | | 8 | 5 | Geometric Graph |
| | | | | 6 | 9 | Geometric Graph |
| | | | | 2 | 3 | Geometric Graph |
| Badger_ Summer3 | 35 | 0.592 | Geometric Graph | 5 | 2 | Geometric Graph |
| | | | | 1 | 9 | Geometric Graph |
| | | | | 6 | 9 | Geometric Graph |
| | | | | 2 | 3 | Geometric Graph |
| | | | | 4 | 5 | Geometric Graph |
| Badger_ Summer4 | 23 | 0.548 | Geometric Graph | 1 | 9 | Geometric Graph |
| | | | | 6 | 8 | Geometric Graph |
| | | | | 4 | 5 | Geometric Graph |

Table S6: Shows the predicted generative models for the Chimpanzee networks and groups across the year.

| Main Network | | | | Modules | | |
|---|---|---|---|---|---|---|
| Network | #Nodes | Prediction | Modularity | Module | #Nodes | Prediction |
| Chimp_Jan (50m) | 33 | 0.0023 | Geometric Graph | 1 | 30 | Geometric Graph |
| | | | | 2 | 3 | Geometric Graph |
| Chimp_Jan (5m) | 33 | 0.0069 | Geometric Graph | 2 | 5 | Geometric Graph |
| | | | | 6 | 5 | Geometric Graph |
| | | | | 4 | 7 | Geometric Graph |
| | | | | 3 | 5 | Scale-Free |
| | | | | 5 | 5 | Geometric Graph |
| | | | | 1 | 3 | Geometric Graph |
| Chimp_Feb (50m) | 31 | 0 | Geometric Graph | - | - | - |
| Chimp_Feb (5m) | 31 | 0.025 | Geometric Graph | 2 | 24 | Geometric Graph |
| | | | | 1 | 2 | Erdös- Rényi |
| | | | | 3 | 5 | Scale-Free |
| Chimp_Mar (50m) | 29 | 0 | Geometric Graph | - | - | - |
| Chimp_Mar (5m) | 28 | 0.0267 | Geometric Graph | 1 | 8 | Geometric Graph |
| | | | | 2 | 13 | Scale-Free |
| | | | | 3 | 3 | Geometric Graph |
| | | | | 5 | 2 | Scale-Free |
| | | | | 4 | 2 | Scale-Free |
| Chimp_Apr (50m) | 33 | 0 | Geometric Graph | - | - | - |
| Chimp_Apr (5m) | 32 | 0.255 | Scale-Free | 2 | 11 | Geometric Graph |
| | | | | 1 | 11 | Geometric Graph |
| | | | | 3 | 7 | Geometric Graph |
| | | | | 4 | 2 | Scale-Free |
| Chimp_May (50m) | 33 | 0 | Geometric Graph | - | - | - |
| Chimp_Jun (50m) | 33 | 0.002 | Geometric Graph | 1 | 26 | Geometric Graph |
| | | | | 2 | 6 | Geometric Graph |
| Chimp_Jun (5m) | 32 | 0.0314 | Geometric Graph | 4 | 19 | Scale-Free |
| | | | | 5 | 4 | Geometric Graph |
| | | | | 3 | 4 | Scale-Free |
| | | | | 2 | 2 | Scale-Free |
| | | | | 1 | 2 | Scale-Free |

| Main Network | | | | Modules | | |
|---|---|---|---|---|---|---|
| Network | #Nodes | Prediction | Modularity | Module | #Nodes | Prediction |
| Chimp_Jul (50m) | 37 | 0.060 | Geometric Graph | 1 | 4 | Geometric Graph |
| | | | | 2 | 28 | Geometric Graph |
| | | | | 3 | 5 | Geometric Graph |
| Chimp_Jul (5m) | 33 | 0.089 | Geometric Graph | 2 | 4 | Geometric Graph |
| | | | | 1 | 23 | Geometric Graph |
| | | | | 3 | 5 | Geometric Graph |
| Chimp_Aug (50m) | 36 | 0 | Geometric Graph | - | - | - |
| Chimp_Aug (5m) | 35 | 0.244 | Scale-Free | 2 | 17 | Scale-Free |
| | | | | 1 | 11 | Scale-Free |
| | | | | 3 | 6 | Geometric Graph |
| Chimp_Dec (50m) | 28 | 0.0081 | Geometric Graph | 1 | 24 | Small World |
| | | | | 2 | 2 | Small World |
| Chimp_Dec (5m) | 26 | 0.235 | Geometric Graph | 1 | 14 | Geometric Graph |
| | | | | 2 | 11 | Geometric Graph |

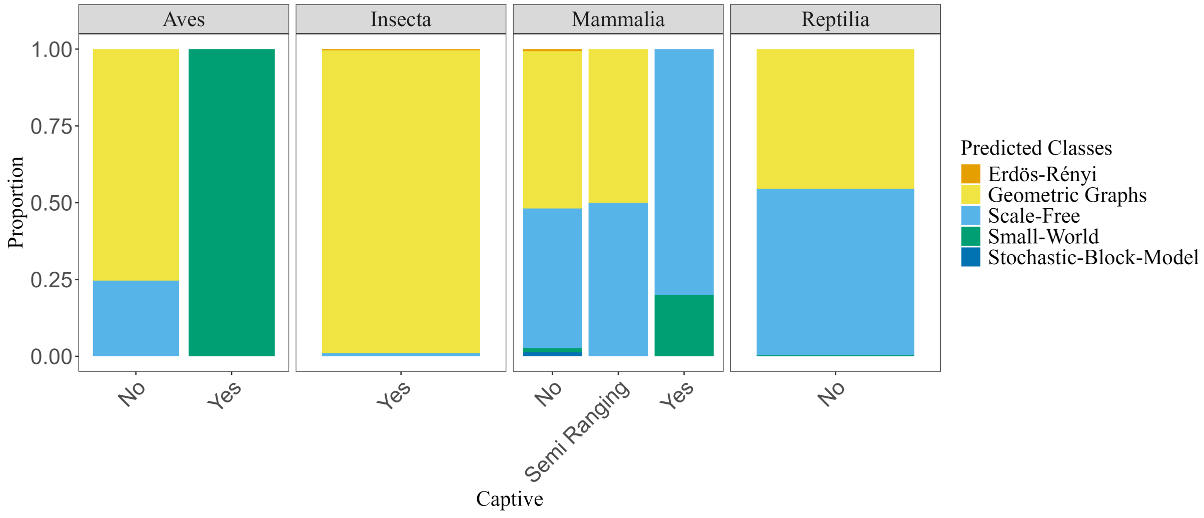


Fig. S1: Classification of animal social networks based on the captive status of animal subjects. Geometric graph models were common across various taxa for both captive and wild animals, particularly among captive insects, wild birds, semi-ranged animals, and wild mammals.

Table S7: Shows the predicted generative models for the Ant (*C. fellah*) network and groups across the six colonies.

| Main Network | | | | Modules | | |
|---|---|---|---|---|---|---|
| Network | #Nodes | Modularity | Prediction | Module | #Nodes | Prediction |
| Insecta_Ant_Colony_1 | 113 | 0.003 | Geometric Graph | 3 | 51 | Small World |
| | | | | 1 | 58 | Small World |
| | | | | 2 | 4 | Small World |
| Colony_1-Day05 | 113 | 0.139 | Geometric Graph | 1 | 69 | Geometric Graph |
| | | | | 2 | 44 | Scale Free |
| Colony_1-Day20 | 99 | 0.160 | Geometric Graph | 1 | 41 | Geometric Graph |
| | | | | 2 | 58 | Small World |
| Colony_1-Day35 | 56 | 0.049 | Geometric Graph | 3 | 30 | Geometric Graph |
| | | | | 1 | 23 | Geometric Graph |
| | | | | 2 | 2 | Geometric Graph |
| Insecta_Ant_Colony_2 | 131 | 0 | Erdös- Rényi | x | x | x |
| Colony_2-Day05 | 131 | 0.0937 | Geometric Graph | 2 | 62 | Geometric Graph |
| | | | | 1 | 69 | Geometric Graph |
| Colony_2-Day20 | 111 | 0.0909 | Geometric Graph | 4 | 14 | Geometric Graph |
| | | | | 2 | 61 | Geometric Graph |
| | | | | 3 | 11 | Geometric Graph |
| | | | | 1 | 25 | Geometric Graph |
| Colony_2-Day35 | 58 | 0.0804 | Geometric Graph | 4 | 11 | Scale Free |
| | | | | 3 | 5 | Geometric Graph |
| | | | | 2 | 35 | Geometric Graph |
| | | | | 1 | 7 | Scale Free |
| Insecta_Ant_Colony_3 | 160 | 0 | Erdös- Rényi | x | x | x |
| Colony_3-Day05 | 160 | 0.104 | Geometric Graph | 2 | 102 | Erdös- Rényi |
| | | | | 1 | 58 | Geometric Graph |
| Colony_3-Day20 | 141 | 0.109 | Geometric Graph | 1 | 97 | Geometric Graph |
| | | | | 2 | 43 | Geometric Graph |
| Colony_3-Day35 | 58 | 0.0751 | Geometric Graph | 3 | 27 | Geometric Graph |

| Main Network | | | | Modules | | |
|---|---|---|---|---|---|---|
| Network | #Nodes | Modularity | Prediction | Module | #Nodes | Prediction |
| Insecta_Ant_Colony_4 | 102 | 0.00002 | Erdös-Rényi | 1 | 97 | Erdös-Rényi |
| | | | | 2 | 5 | Small World |
| Colony_4-Day05 | 102 | 0.0519 | Geometric Graph | 3 | 37 | Geometric Graph |
| | | | | 2 | 52 | Erdös-Rényi |
| | | | | 1 | 11 | Small World |
| | | | | 4 | 2 | Small World |
| Colony_4-Day20 | 73 | 0.0301 | Erdös-Rényi | 1 | 97 | Scale Free |
| | | | | 2 | 43 | Geometric Graph |
| | | | | 3 | 43 | Geometric Graph |
| Colony_4-Day35 | 35 | 0.0014 | Scale Free | 1 | 24 | Geometric Graph |
| | | | | 2 | 9 | Scale Free |
| Insecta_Ant_Colony_5 | 152 | 0 | Erdös-Rényi | x | x | x |
| Colony_5-Day05 | 152 | 0.080 | Geometric Graph | 1 | 91 | Geometric Graph |
| | | | | 2 | 61 | Geometric Graph |
| Colony_5-Day20 | 133 | 0.132 | Geometric Graph | 1 | 55 | Geometric Graph |
| | | | | 2 | 78 | Geometric Graph |
| Colony_5-Day35 | 66 | 0.0428 | Geometric Graph | 3 | 34 | Geometric Graph |
| | | | | 2 | 25 | Scale Free |
| | | | | 1 | 4 | Geometric Graph |
| Insecta_Ant_Colony_6 | 164 | 0.0003 | Erdös-Rényi | 1 | 158 | Erdös-Rényi |
| | | | | 2 | 6 | Small World |
| Colony_6-Day05 | 164 | 0.099 | Geometric Graph | 1 | 67 | Erdös-Rényi |
| | | | | 2 | 97 | Erdös-Rényi |
| Colony_6-Day20 | 143 | 0.140 | Geometric Graph | 1 | 73 | Geometric Graph |
| | | | | 2 | 70 | Geometric Graph |
| Colony_6-Day35 | 91 | 0.0471 | Geometric Graph | 1 | 58 | Geometric Graph |
| | | | | 2 | 11 | Geometric Graph |
| | | | | 3 | 5 | Geometric Graph |
| | | | | 4 | 17 | Geometric Graph |

Table S8: Distinct species list

| Network ID | Species | Association type | Data collection method |
|---|---|---|---|
| Aves-barn-swallow | *Hirundo rustica* | Physical contact | Logger |
| Aves-geese | *Branta leucopsis* | Foraging | Survey scan |
| Aves-hens-pecking-order | *Gallus gallus* | Dominance | Direct observation |
| Aves-songbird | *Haemorhous mexicanus* | Social projection bipartite | RFID |
| Aves-sparrow | *Zonotrichia atricapilla* | Group membership | Survey scan |
| Aves-thornbill-farine | *Acanthiza* sp. | Group membership | Survey scan |
| Aves-weaver | *Philetairus socius* | Social projection bipartite | Mark recapture |
| Badger | *Meles meles* | Spatial proximity | RFID |
| Chimpanzee | *Pan troglodytes* | Spatial proximity | Focal sampling |
| Dogs | *Canis familiaris* | Spatial proximity | RFID |
| Giraffe | *Giraffa camelopardalis* | Group membership | Survey scan |
| Ants | *Camponotus fellah* | Physical contact | Video |
| Ants-trophallaxis | *Camponotus pennsylvanicus* | Physical contact | Video |
| Beetles | *Bolitotherus cornutus* | Spatial proximity | Survey scan |
| Asian elephant | *Elephas maximus* | Dominance | Focal sampling |
| Babbons | *Papio cynocephalus* | Spatial proximity | Focal sampling |
| Bats | *Myotis sodalis* | Social projection bipartite | RFID |
| Bison | *Bison bison* | Dominance | Survey scan |
| Cattle | *Bos taurus* | Dominance | Survey scan |
| Dolphin | *Tursiops truncatus* | Social projection bipartite | Survey scan |
| Elephant-seal | *Mirounga angustirostris* | Dominance | Survey scan |
| Hyena | *Crocuta crocuta* | Group membership | Survey scan |
| Kangaroo | *Macropus giganteus* | Spatial proximity | Survey scan |
| Macaque-contact | *Macaca mulatta* | Physical contact | Survey scan |
| Macaque | *Macaca fuscata* | Dominance/grooming | Survey scan/focal sampling |
| Racoon | *Procyon lotor* | Spatial proximity | Logger |
| Sheep | *Ovis canadensis* | Dominance | Focal sampling |
| Spider-monkey | *Ateles hybridus* | Physical contact | Focal sampling |
| Vampire-bats | *Desmodus rotundus* | Trophallaxis | Video |
| Voles | *Microtus agrestis* | Social projection bipartite | Mark recapture |
| Zebra | *Equus grevyi* | Group membership | Survey scan |
| Lizard | *Tiliqua rugosa* | Spatial proximity | Logger |
| Tortoise | *Gopherus agassizii* | Social projection bipartite | RFID |

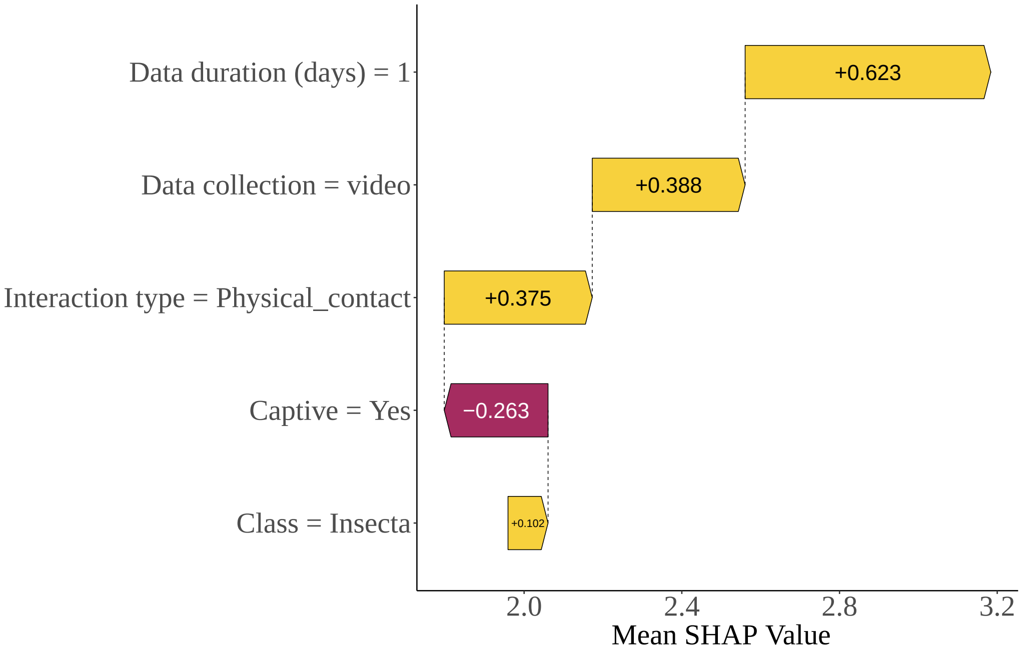


Fig. S2: SHAP Waterfall plot explaining how the network metadata of the empirical networks for a single data instance impact the final output of the geometric graph for the model without species. Metadata impacting the model prediction higher (towards the right) are shown in yellow and those pushing the model prediction lower (toward the left) are shown in red.

(a)

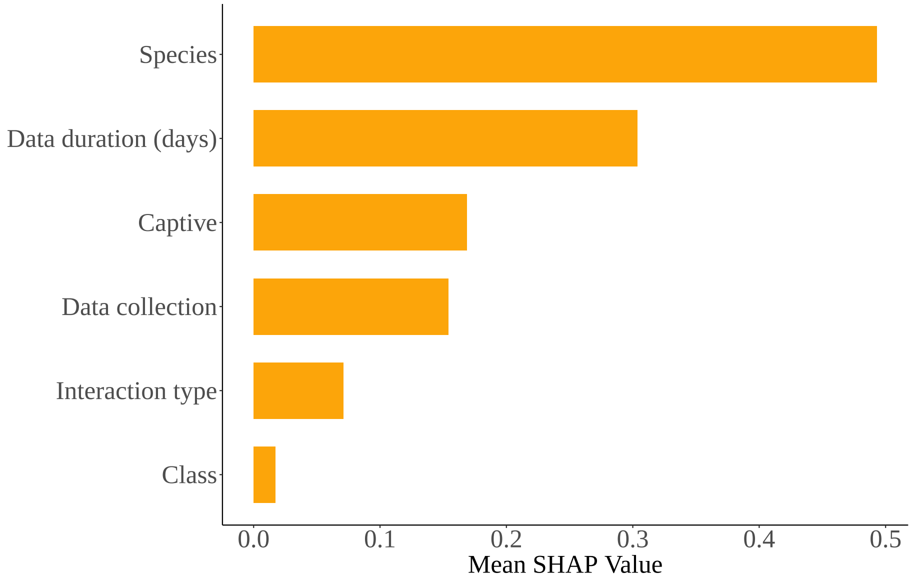

Species
Data duration (days)
Captive
Data collection
Interaction type
Class
0.0
0.1
0.2
0.3
0.4
0.5
Mean SHAP Value


(b)

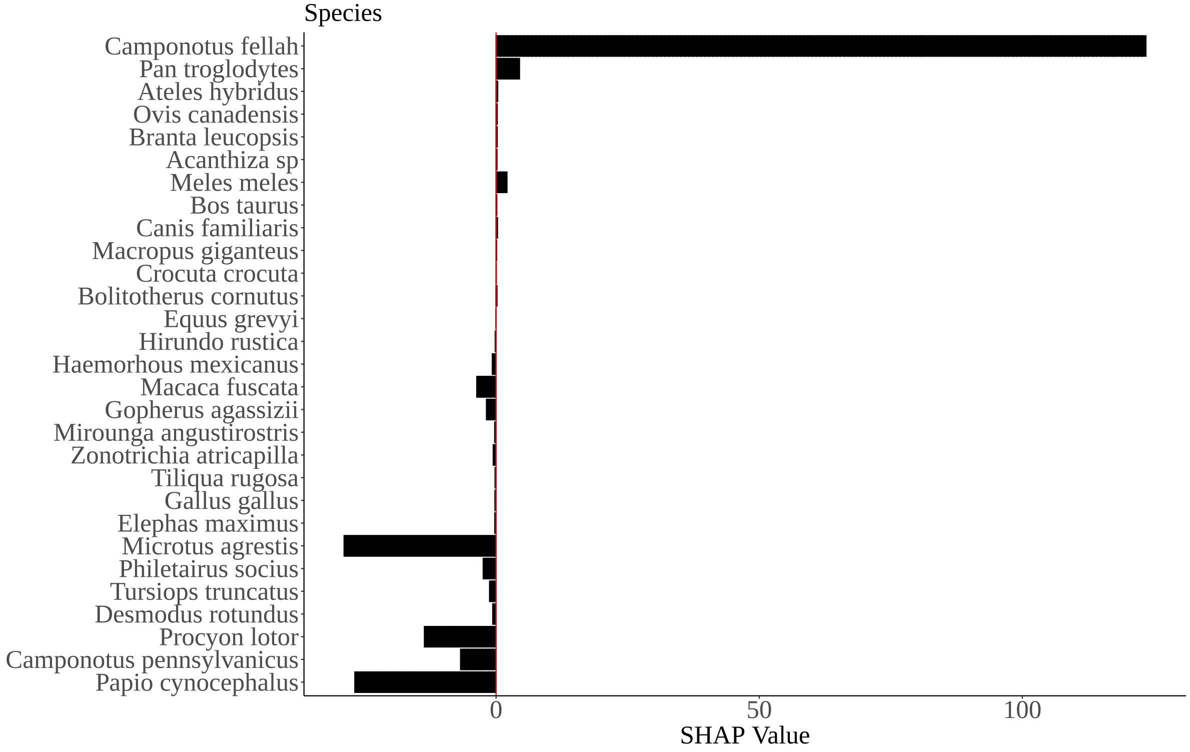

Species
Camponotus fellah
Pan troglodytes
Ateles hybridus
Ovis canadensis
Branta leucopsis
Acanthiza sp
Meles meles
Bos taurus
Canis familiaris
Macropus giganteus
Crocuta crocuta
Bolitotherus cornutus
Equus grevyi
Hirundo rustica
Haemorhous mexicanus
Macaca fuscata
Gopherus agassizii
Mirounga angustirostris
Zonotrichia atricapilla
Tiliqua rugosa
Gallus gallus
Elephas maximus
Microtus agrestis
Philetairus socius
Tursiops truncatus
Desmodus rotundus
Procyon lotor
Camponotus pennsylvanicus
Papio cynocephalus
0
50
100
SHAP Value

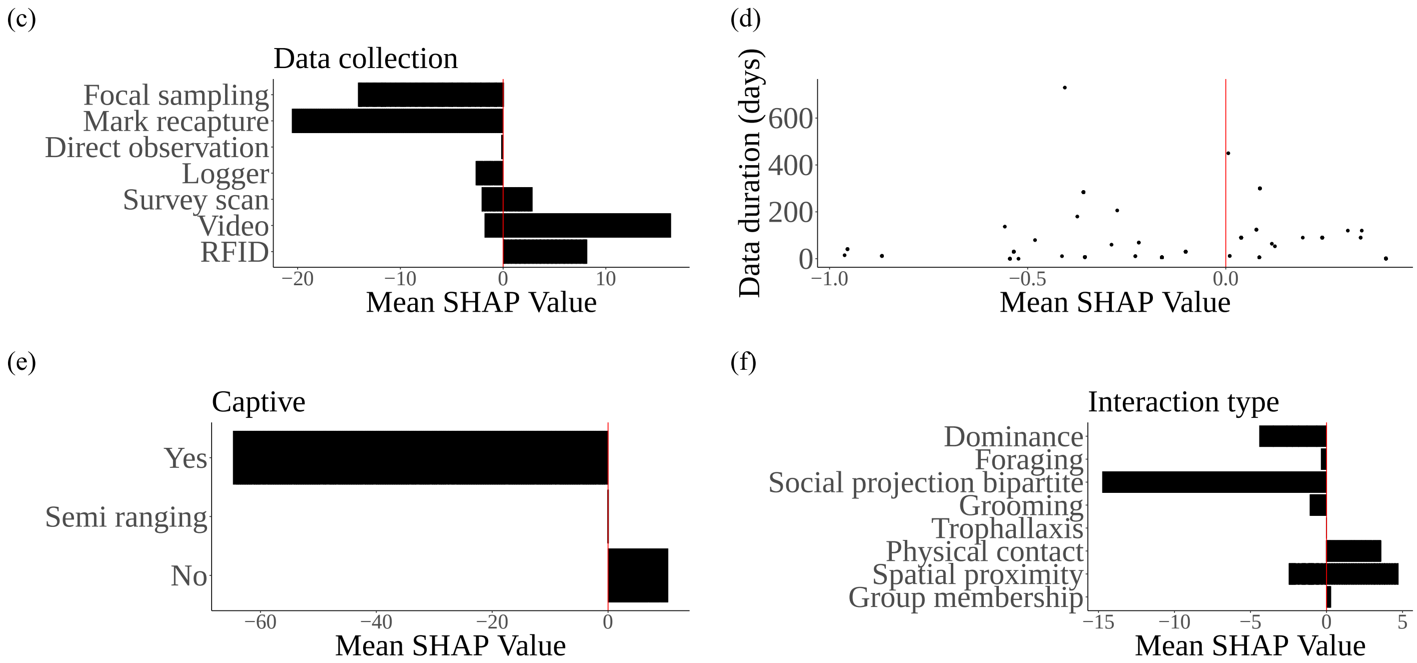


Fig. S3: SHAP variabe importance plot (a) the model with species, and dependency plots illustrating the relationship between: (b) species, (c) data collection methods, (d) data duration, (e) captive, and (f) social interactions on the geometric graph models global output.

(a)

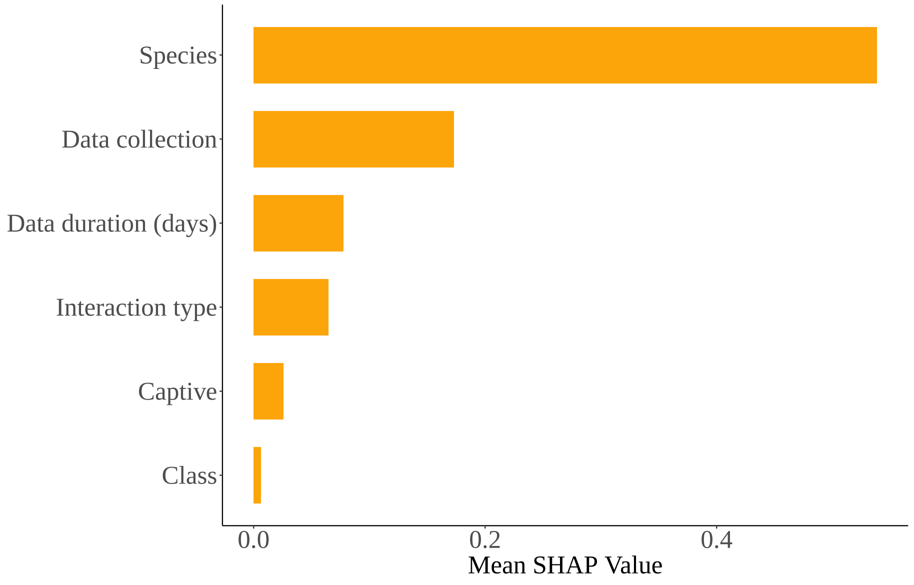
Species
Data collection
Data duration (days)
Interaction type
Captive
Class
0.0
0.2
0.4
Mean SHAP Value

(b)

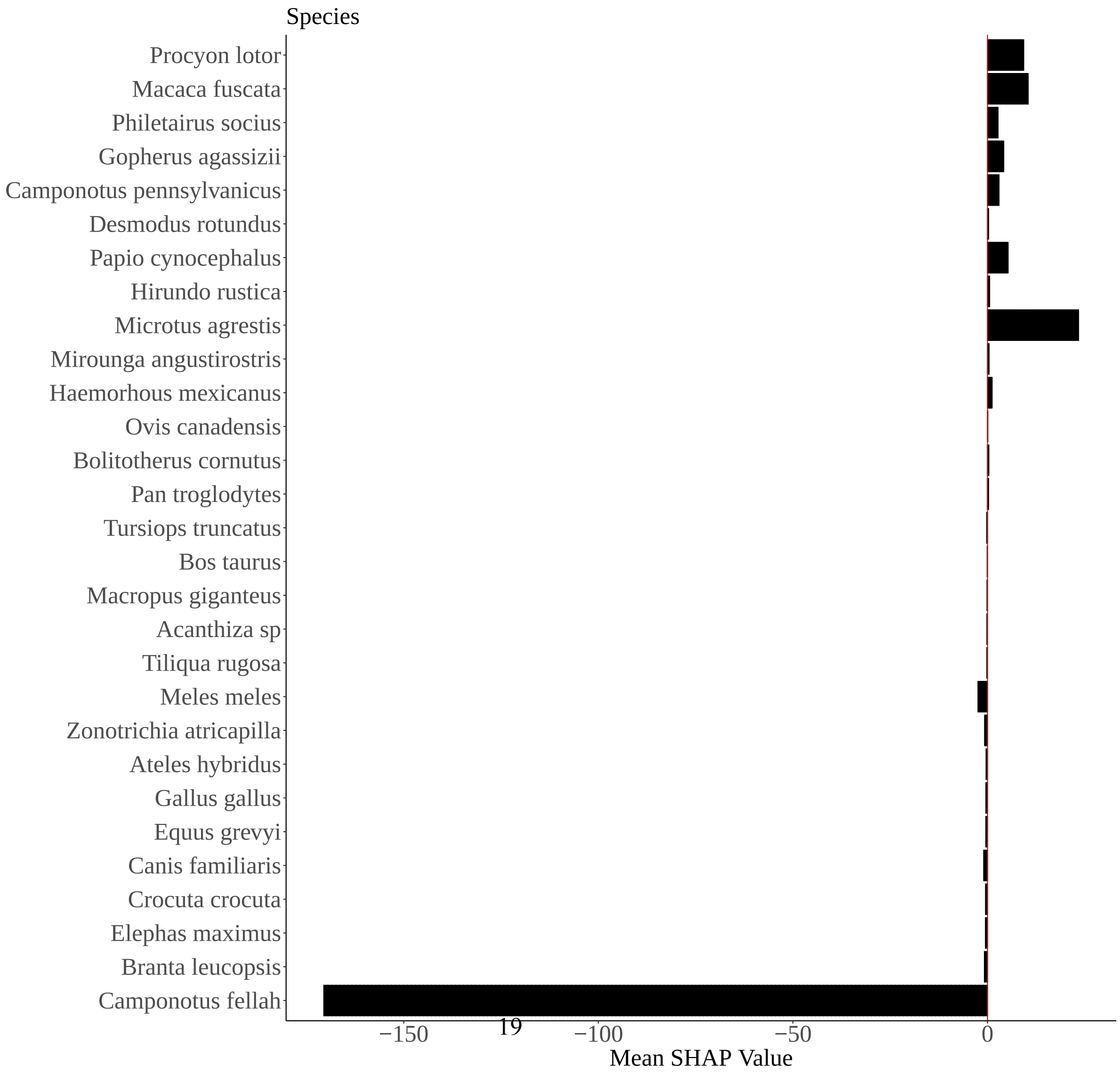
Species
Procyon lotor
Macaca fuscata
Philetairus socius
Gopherus agassizii
Camponotus pennsylvanicus
Desmodus rotundus
Papio cynocephalus
Hirundo rustica
Microtus agrestis
Mirounga angustirostris
Haemorhous mexicanus
Ovis canadensis
Bolitotherus cornutus
Pan troglodytes
Tursiops truncatus
Bos taurus
Macropus giganteus
Acanthiza sp
Tiliqua rugosa
Meles meles
Zonotrichia atricapilla
Ateles hybridus
Gallus gallus
Equus grevyi
Canis familiaris
Crocuta crocuta
Elephas maximus
Branta leucopsis
Camponotus fellah
−150
−100
−50
0
Mean SHAP Value

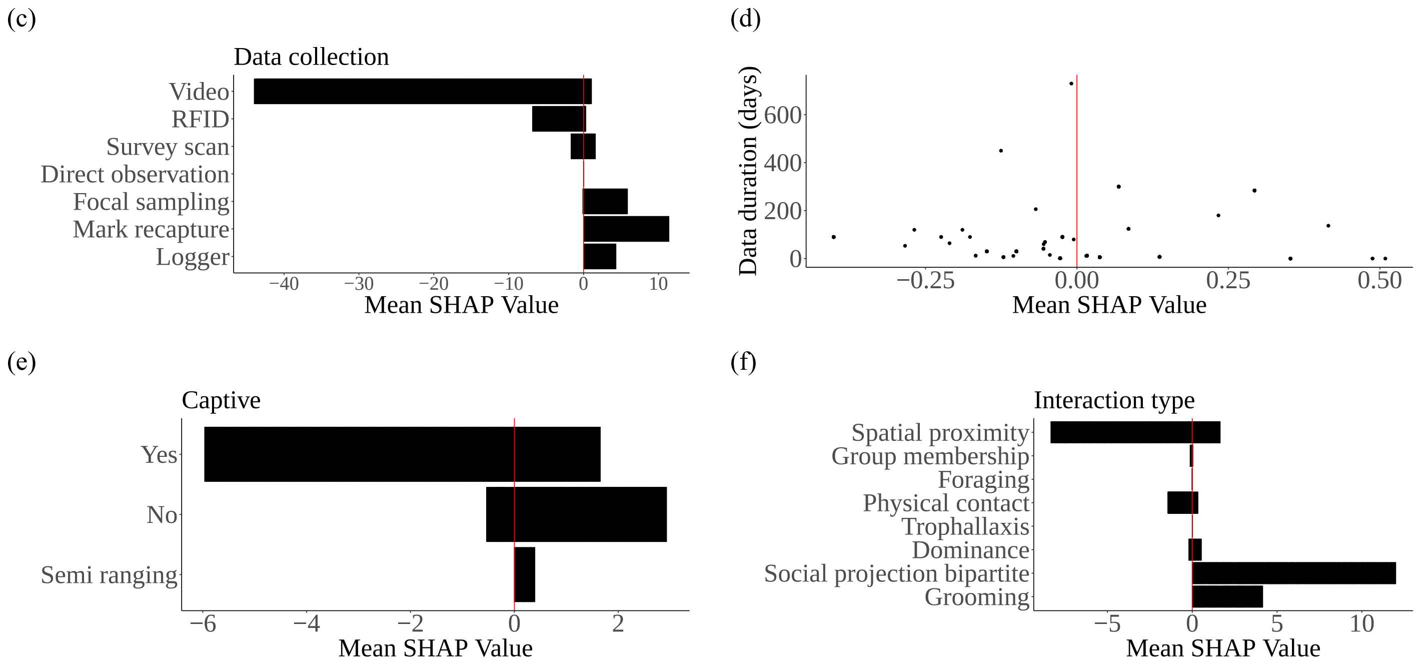


Fig. S4: SHAP variabe importance plot (a) the model with species, and dependency plots illustrating the relationship between: (b) species, (c) data collection methods, (d) data duration, (e) captive, and (f) social interactions on the scale-free models global output.

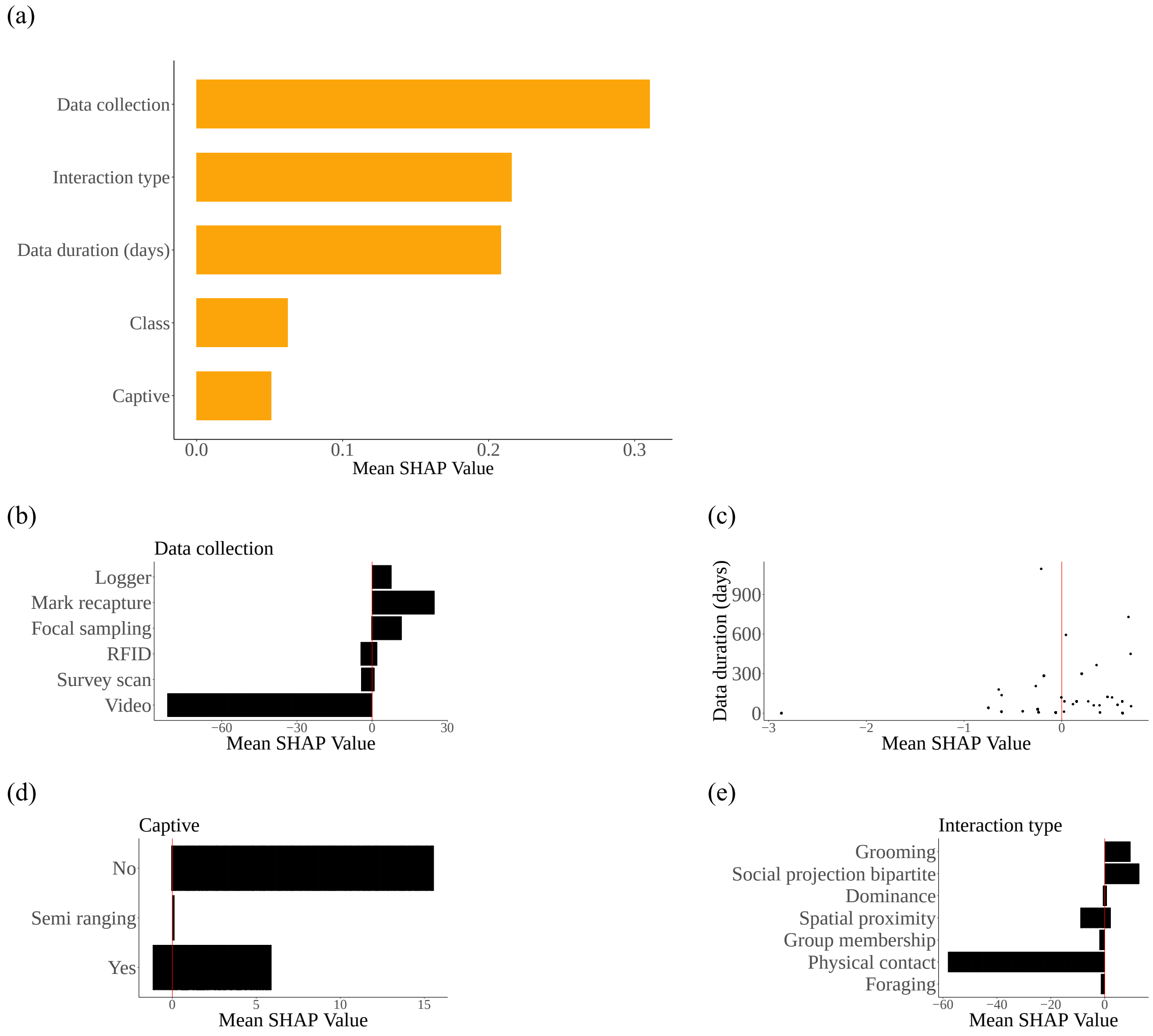


Fig. S5: SHAP variabe importance plot (a) the model without species, and dependency plots illustrating the relationship between: (b) data collection methods, (c) data duration, (d) captive, and (e) social interactions on the scale-free models global output.

(a)

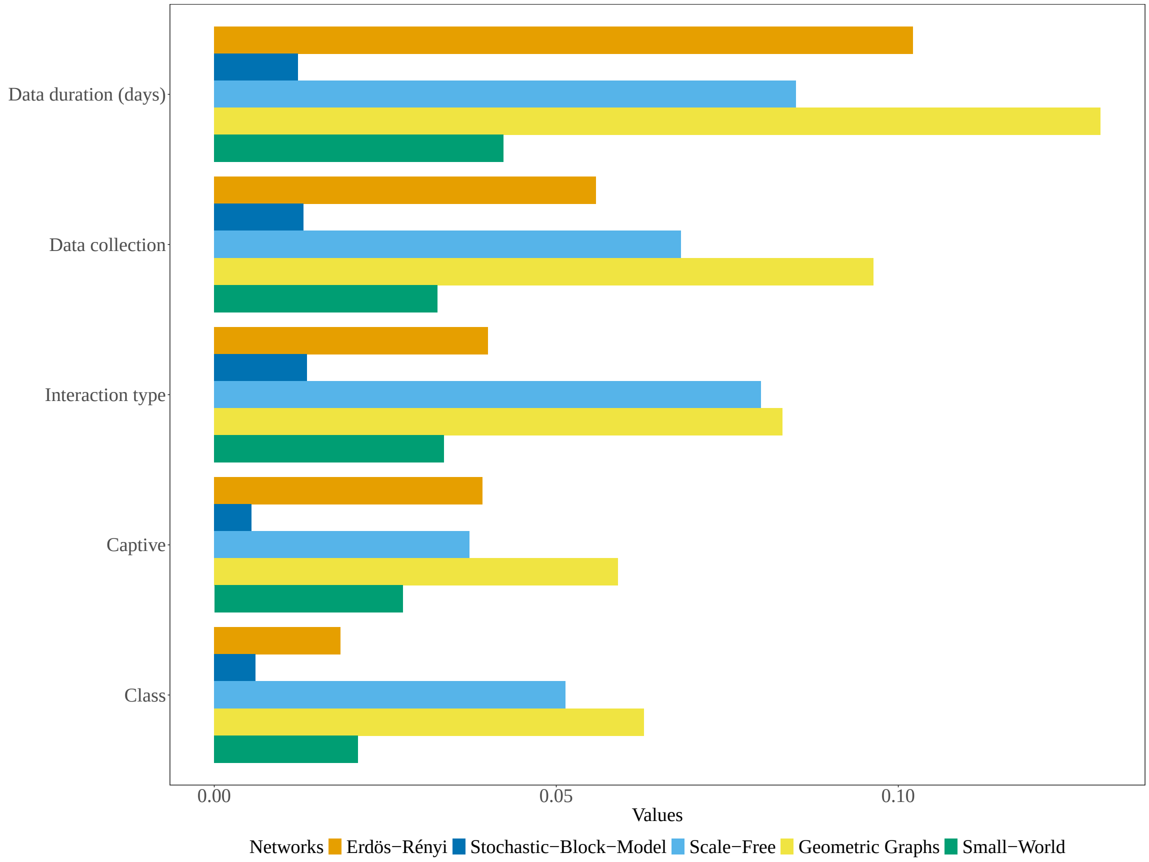
Data duration (days)
Data collection
Interaction type
Captive
Class
0.00
0.05
0.10
Values
Networks
Erdös−Rényi
Stochastic−Block−Model
Scale−Free
Geometric Graphs
Small−World

(b)

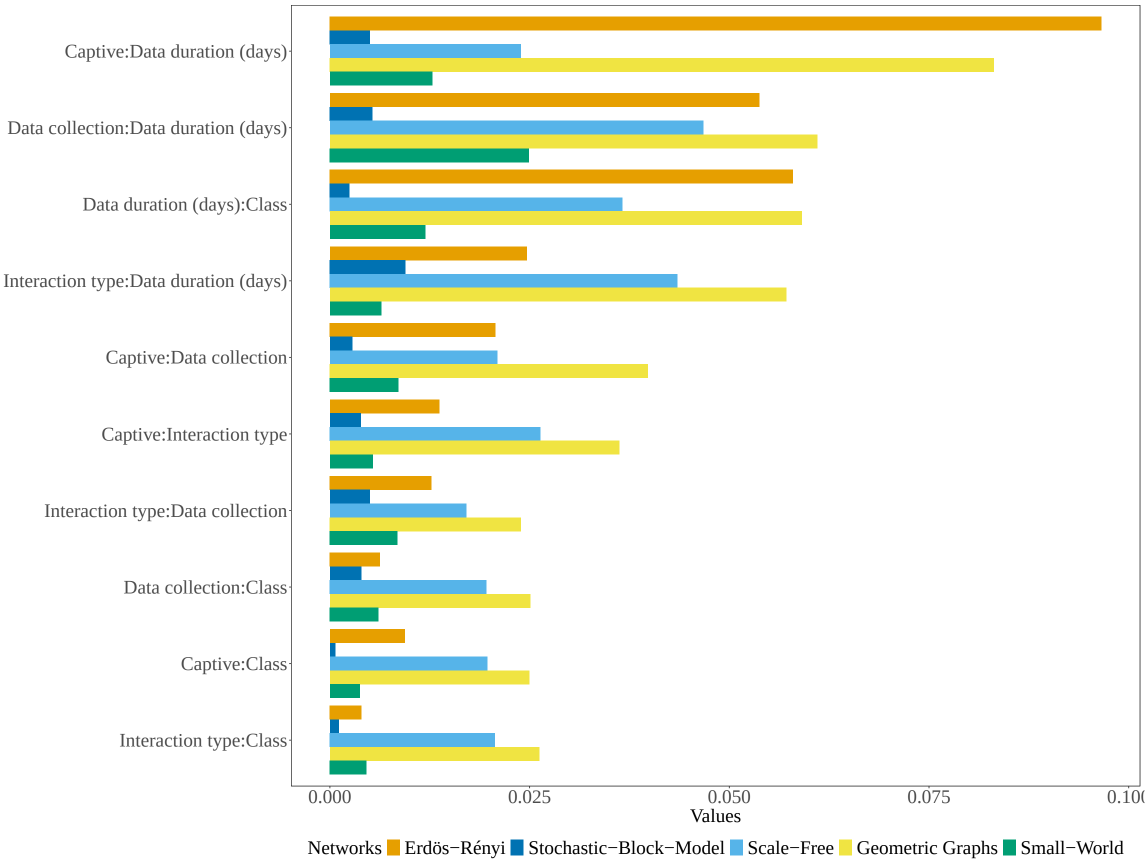
Captive:Data duration (days)
Data collection:Data duration (days)
Data duration (days):Class
Interaction type:Data duration (days)
Captive:Data collection
Captive:Interaction type
Interaction type:Data collection
Data collection:Class
Captive:Class
Interaction type:Class
0.000
0.025
0.050
0.075
0.100
Values
Networks
Erdös−Rényi
Stochastic−Block−Model
Scale−Free
Geometric Graphs
Small−World

(c)

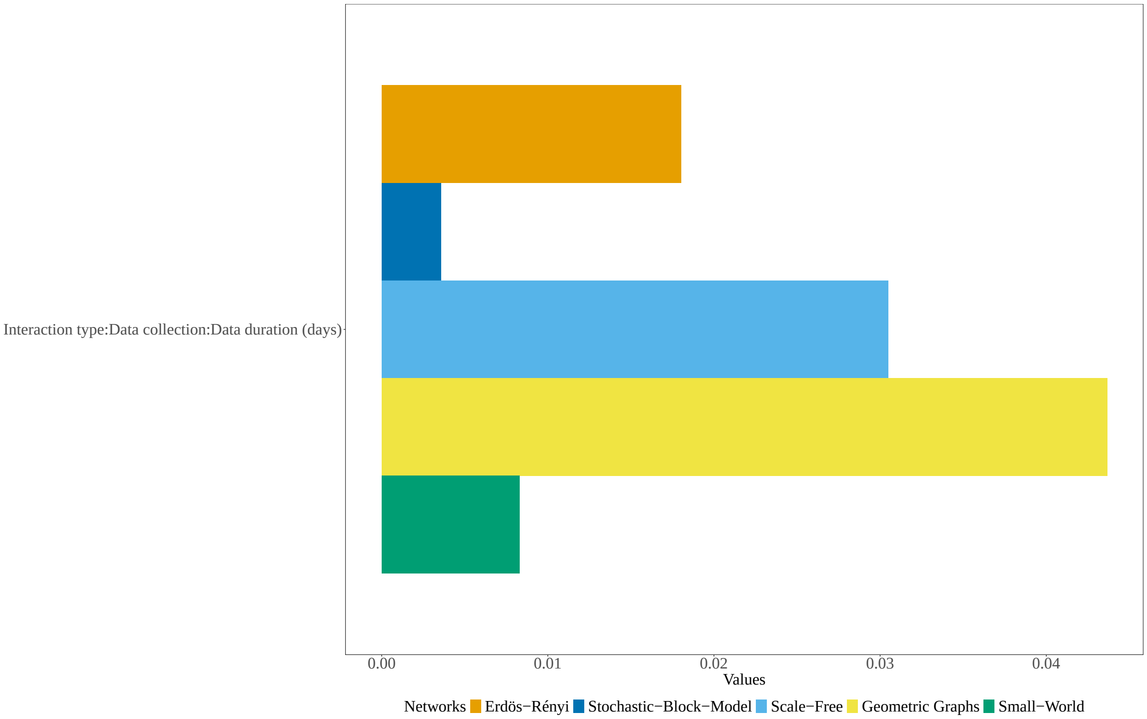


Fig. S6: This plot shows the proportion of predicted variability of the metadata variables with the geometric graph model excluding species associated with (a) overall interaction effect, (b) pairwise interaction effect, and (c) three-way interaction effects. The length of the bar associated with the metadata on the y-axis typically indicates the strength of the overall, pairwise or three-way interaction and how these interactions influence the model's prediction across the generative models. The network correctly predicted by the overall, pairwise and three-way interaction of the metadata is the geometric graph model.

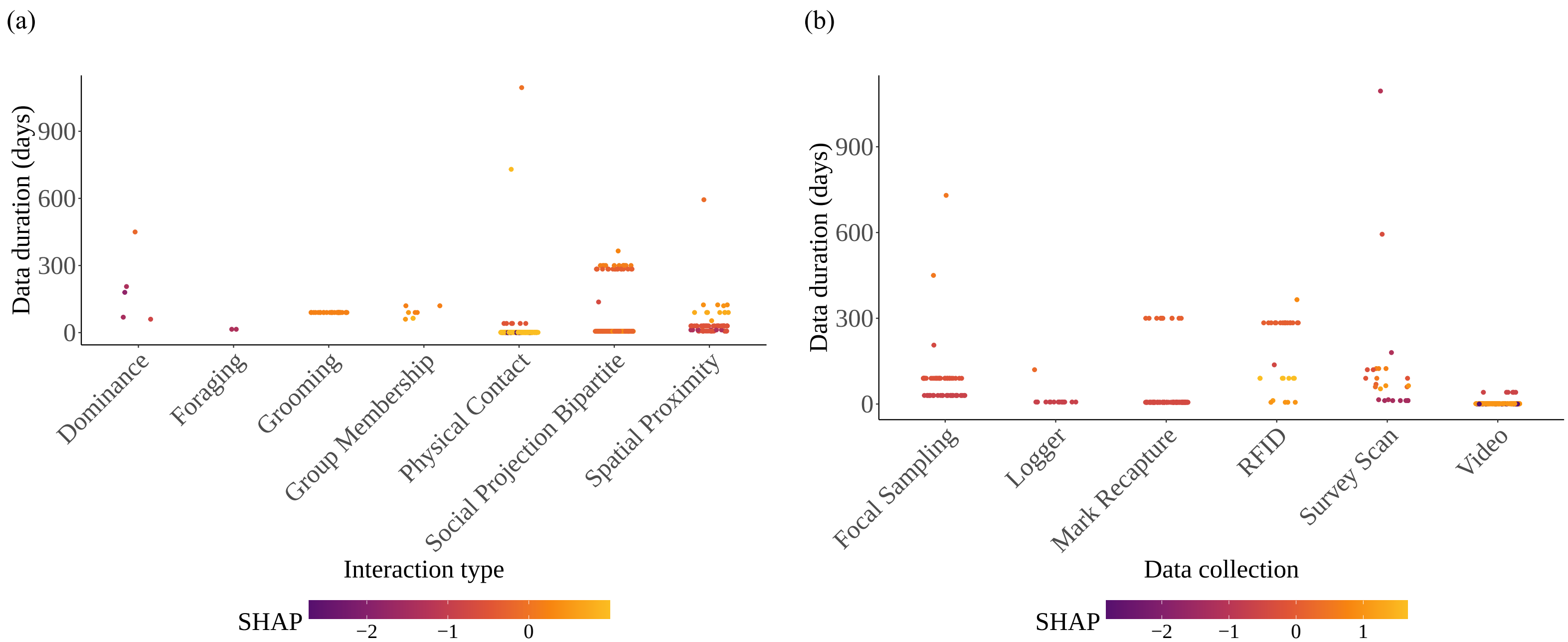
(a)
Data duration (days)
900
600
300
0
Dominance
Foraging
Grooming
Group Membership
Physical Contact
Social Projection Bipartite
Spatial Proximity
Interaction type
SHAP
−2
−1
0
(b)
Data duration (days)
900
600
300
0
Focal Sampling
Logger
Mark Recapture
RFID
Survey Scan
Video
Data collection
SHAP
−2
−1
0
1

(c) (d)

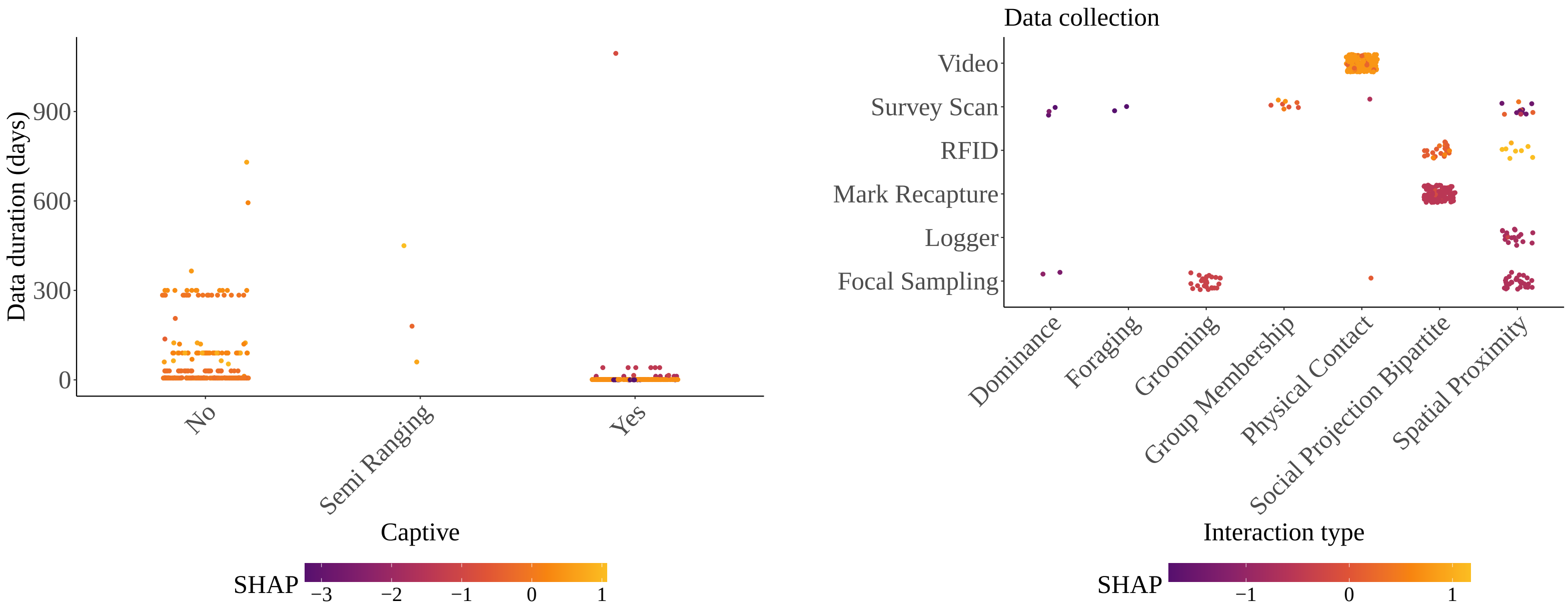
Data duration (days)
0
300
600
900
No
Semi Ranging
Yes
Captive
SHAP
−3
−2
−1
0
1
Data collection
Video
Survey Scan
RFID
Mark Recapture
Logger
Focal Sampling
Dominance
Foraging
Grooming
Group Membership
Physical Contact
Social Projection Bipartite
Spatial Proximity
Interaction type
SHAP
−1
0
1

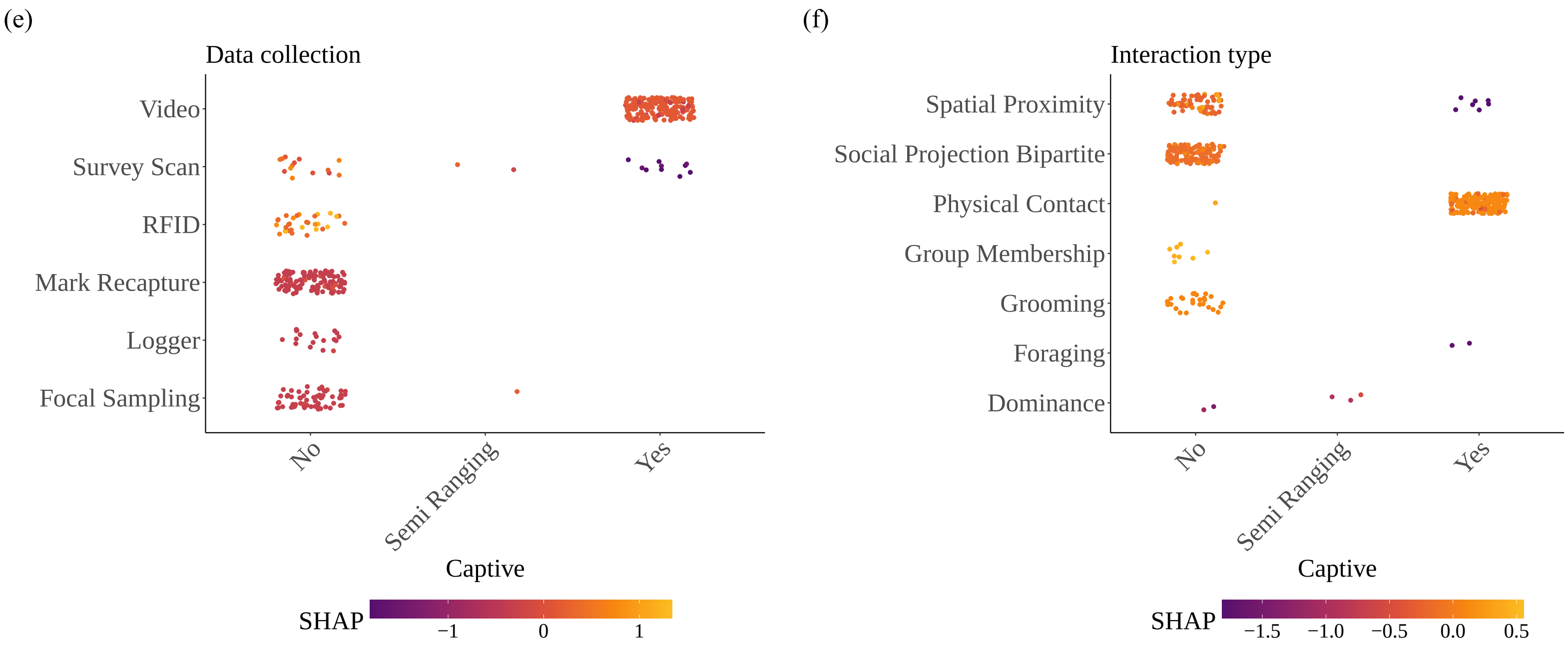


Fig. S7: This plot shows the pairwise interaction effect of (a) data duration (days) and interaction type, (b) data duration (days) and data collection, (c) data duration (days) and captive, (d) data collection and animal interaction type, (e) data collection and captive, and (f) interaction type and captive on the prediction of the geometric graph model without species. This plot provides insights into how the model's prediction changes as different metadata interacts. Overall, this plot helps to understand the model's behavior with respect to metadata interaction. The vertical color scale for the SHAP joint interaction plots indicates interactions that either negatively or positively influence the prediction of the geometric graph model.

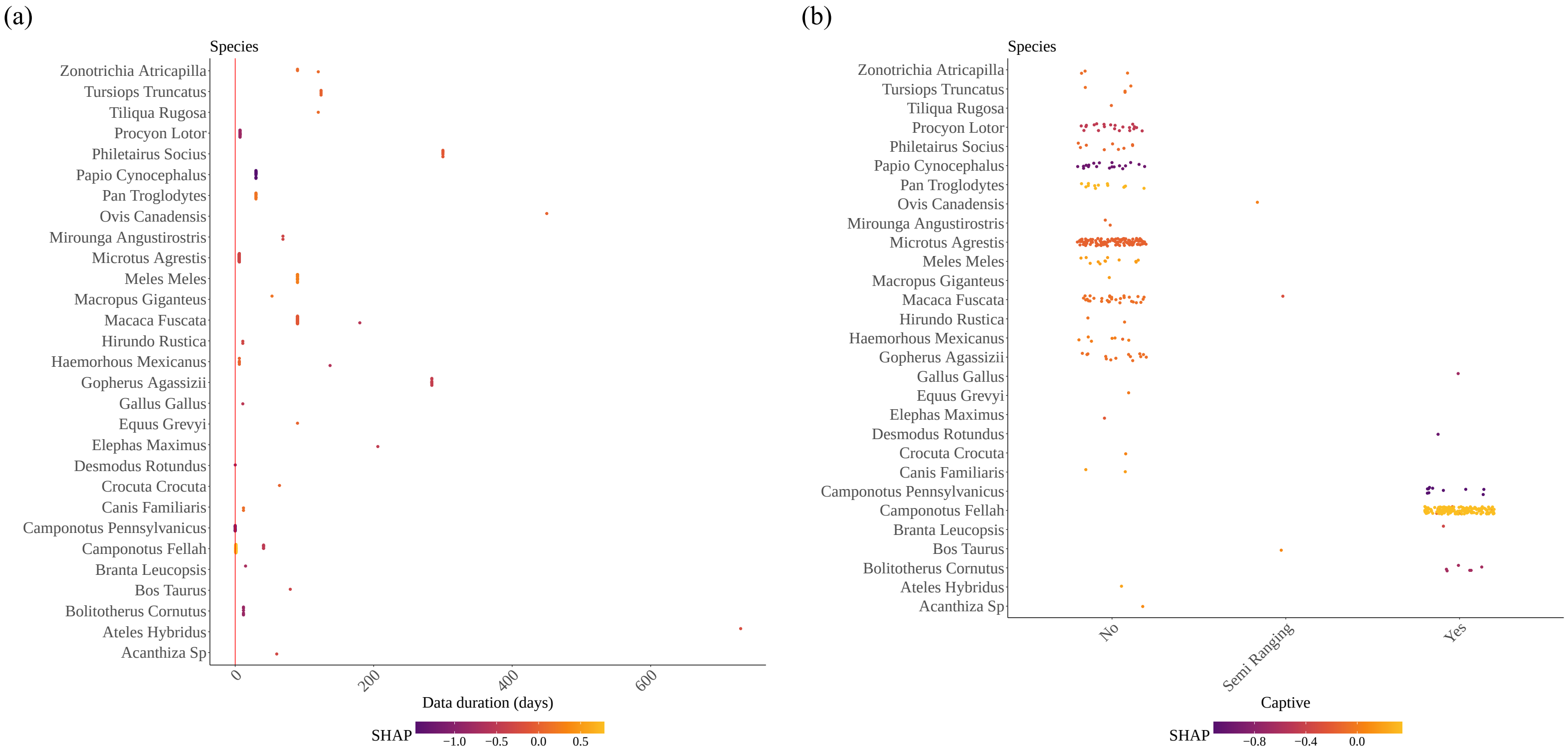
(a)
Species
Zonotrichia Atricapilla
Tursiops Truncatus
Tiliqua Rugosa
Procyon Lotor
Philetairus Socius
Papio Cynocephalus
Pan Troglodytes
Ovis Canadensis
Mirounga Angustirostris
Microtus Agrestis
Meles Meles
Macropus Giganteus
Macaca Fuscata
Hirundo Rustica
Haemorhous Mexicanus
Gopherus Agassizii
Gallus Gallus
Equus Grevyi
Elephas Maximus
Desmodus Rotundus
Crocuta Crocuta
Canis Familiaris
Camponotus Pennsylvanicus
Camponotus Fellah
Branta Leucopsis
Bos Taurus
Bolitotherus Cornutus
Ateles Hybridus
Acanthiza Sp
0
200
400
600
Data duration (days)
SHAP
−1.0
−0.5
0.0
0.5
(b)
Species
Zonotrichia Atricapilla
Tursiops Truncatus
Tiliqua Rugosa
Procyon Lotor
Philetairus Socius
Papio Cynocephalus
Pan Troglodytes
Ovis Canadensis
Mirounga Angustirostris
Microtus Agrestis
Meles Meles
Macropus Giganteus
Macaca Fuscata
Hirundo Rustica
Haemorhous Mexicanus
Gopherus Agassizii
Gallus Gallus
Equus Grevyi
Elephas Maximus
Desmodus Rotundus
Crocuta Crocuta
Canis Familiaris
Camponotus Pennsylvanicus
Camponotus Fellah
Branta Leucopsis
Bos Taurus
Bolitotherus Cornutus
Ateles Hybridus
Acanthiza Sp
No
Semi Ranging
Yes
Captive
SHAP
−0.8
−0.4
0.0

(c) (d)

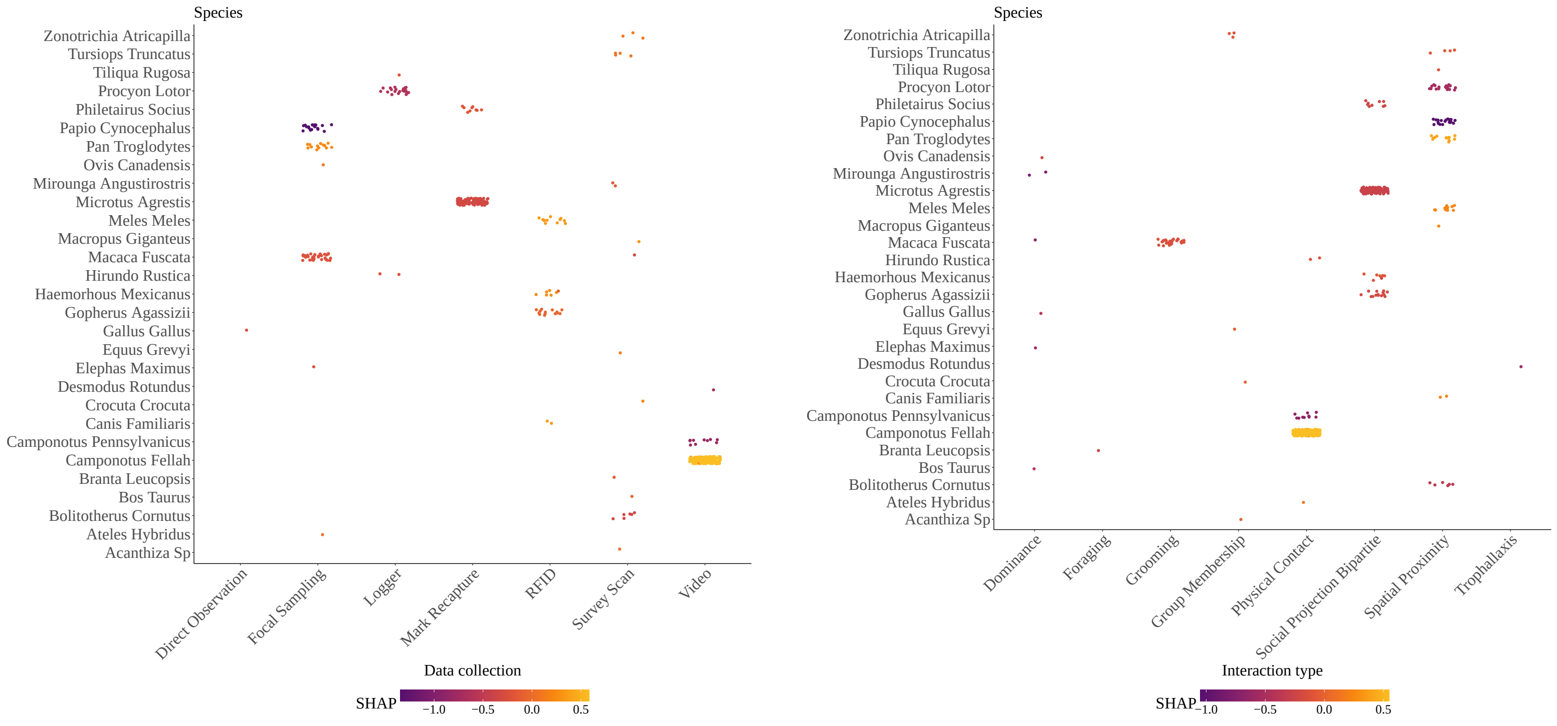
Species
Zonotrichia Atricapilla
Tursiops Truncatus
Tiliqua Rugosa
Procyon Lotor
Philetairus Socius
Papio Cynocephalus
Pan Troglodytes
Ovis Canadensis
Mirounga Angustirostris
Microtus Agrestis
Meles Meles
Macropus Giganteus
Macaca Fuscata
Hirundo Rustica
Haemorhous Mexicanus
Gopherus Agassizii
Gallus Gallus
Equus Grevyi
Elephas Maximus
Desmodus Rotundus
Crocuta Crocuta
Canis Familiaris
Camponotus Pennsylvanicus
Camponotus Fellah
Branta Leucopsis
Bos Taurus
Bolitotherus Cornutus
Ateles Hybridus
Acanthiza Sp
Direct Observation
Focal Sampling
Logger
Mark Recapture
RFID
Survey Scan
Video
Data collection
SHAP
−1.0
−0.5
0.0
0.5
Species
Dominance
Foraging
Grooming
Group Membership
Physical Contact
Social Projection Bipartite
Spatial Proximity
Trophallaxis
Interaction type
SHAP
−1.0
−0.5
0.0
0.5

(e)

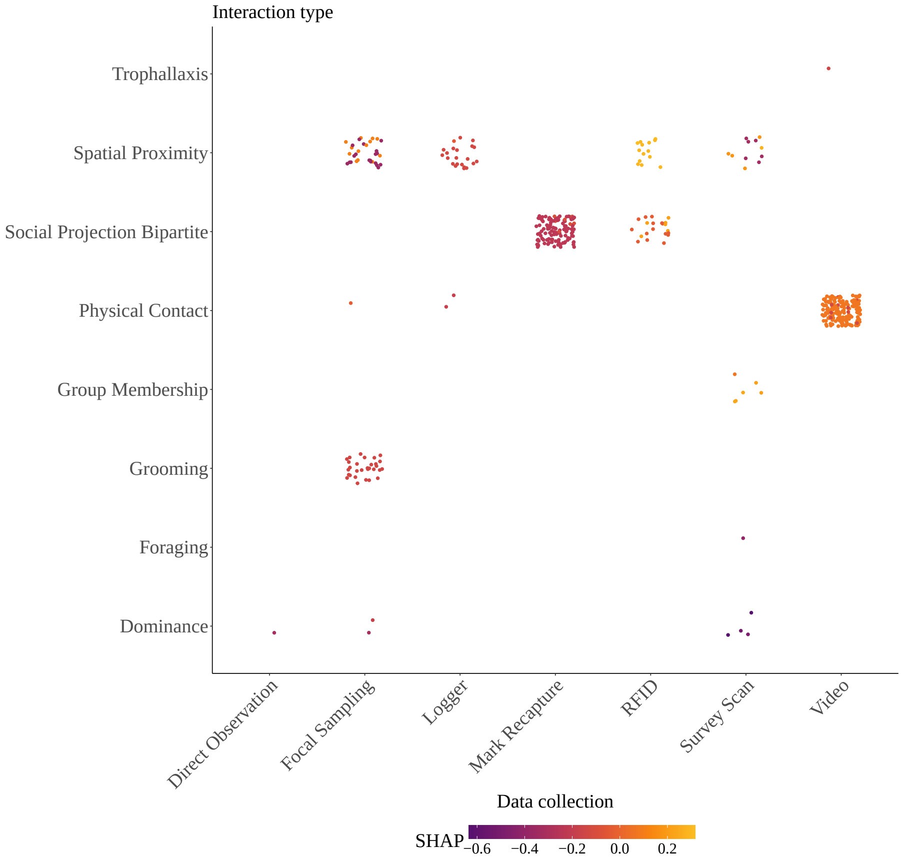


Fig. S8: This plot shows the pairwise interaction effect of (a) species and data duration (days), (b) species and captive status, (c) species and data collection, (d) species and interaction types, (e) interaction type and data collection on the prediction of the geometric graph model with species characteristics.

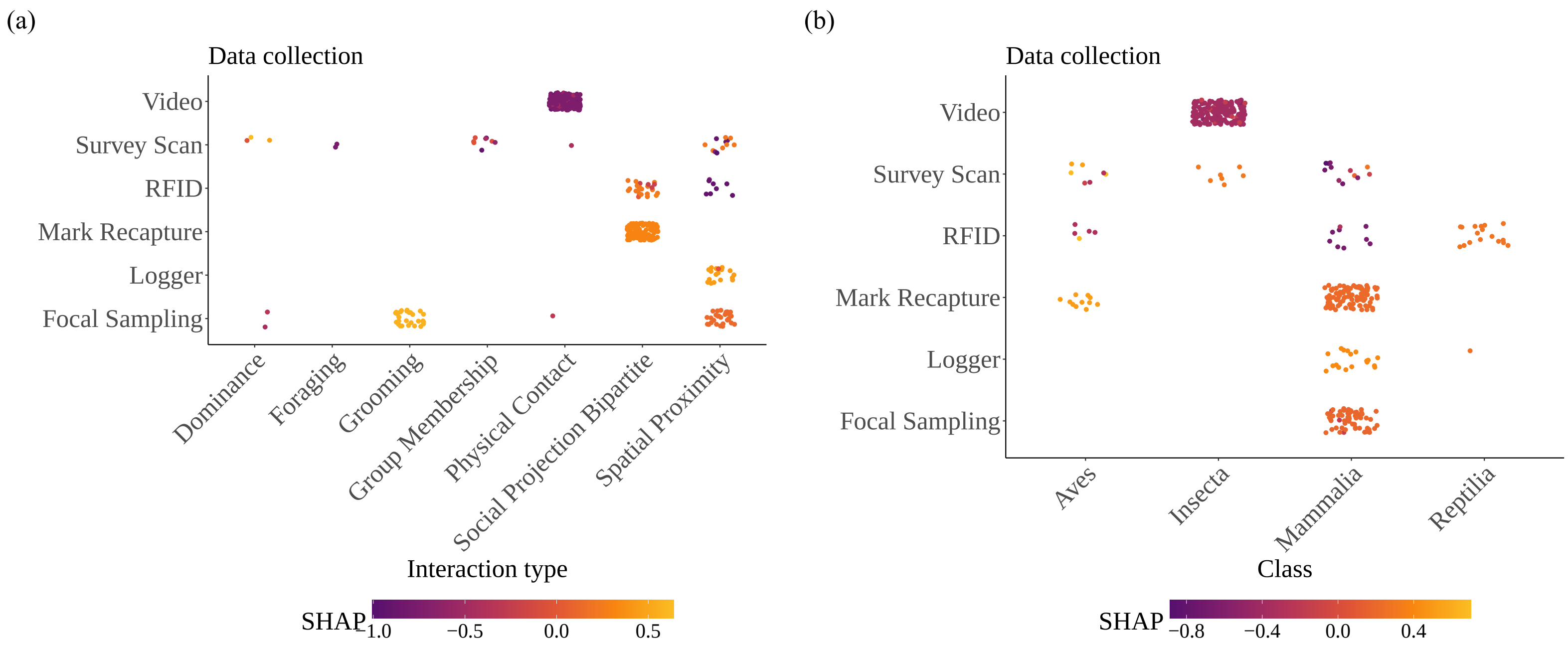

(a)
Data collection
Video
Survey Scan
RFID
Mark Recapture
Logger
Focal Sampling
Dominance
Foraging
Grooming
Group Membership
Physical Contact
Social Projection Bipartite
Spatial Proximity
Interaction type
SHAP
1.0
−0.5
0.0
0.5
(b)
Data collection
Video
Survey Scan
RFID
Mark Recapture
Logger
Focal Sampling
Aves
Insecta
Mammalia
Reptilia
Class
SHAP
−0.8
−0.4
0.0
0.4

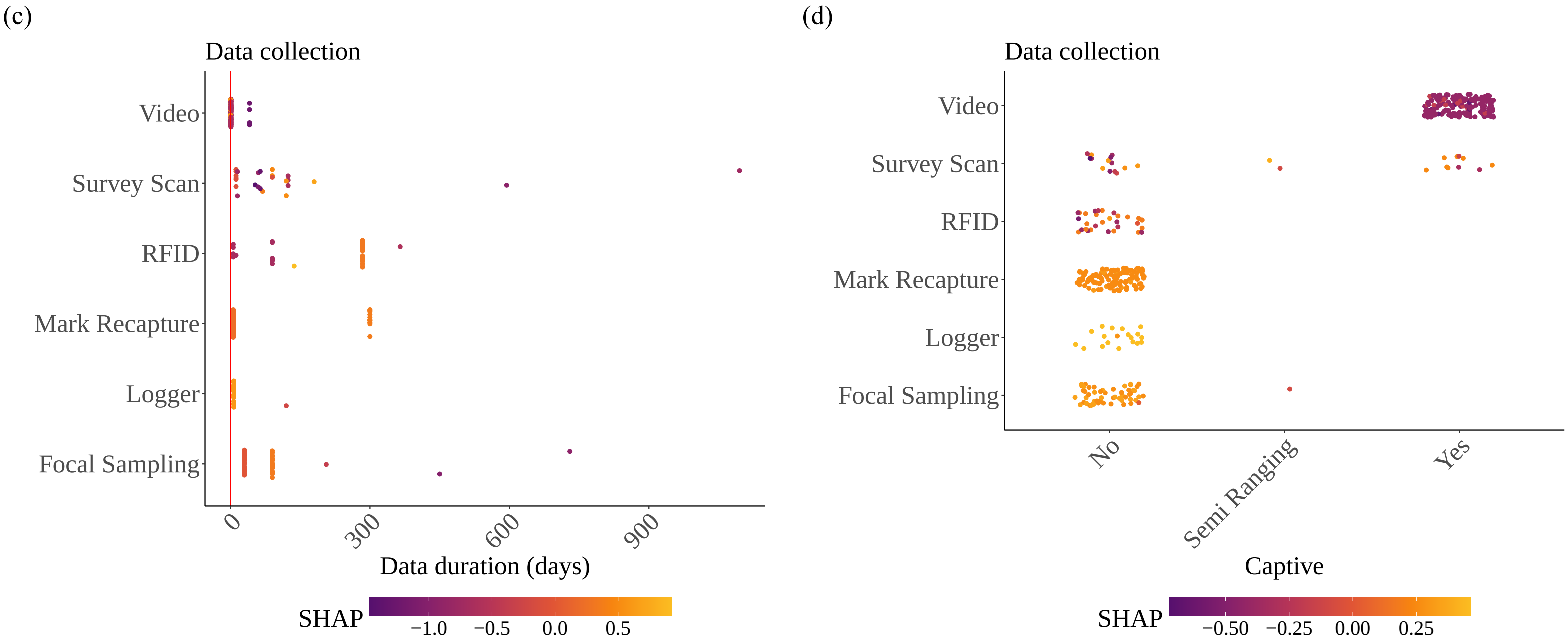


Fig. S9: This plot shows the pairwise interaction effect of (a) data collection and interaction type, (b) data collection and class, (c) data collection and data duration, (d) data collection and captive on the prediction of the scale-free class without species. This plot provides insights into how the model's prediction changes as different metadata interacts. Overall, this plot helps to understand the model's behavior with respect to metadata interaction.

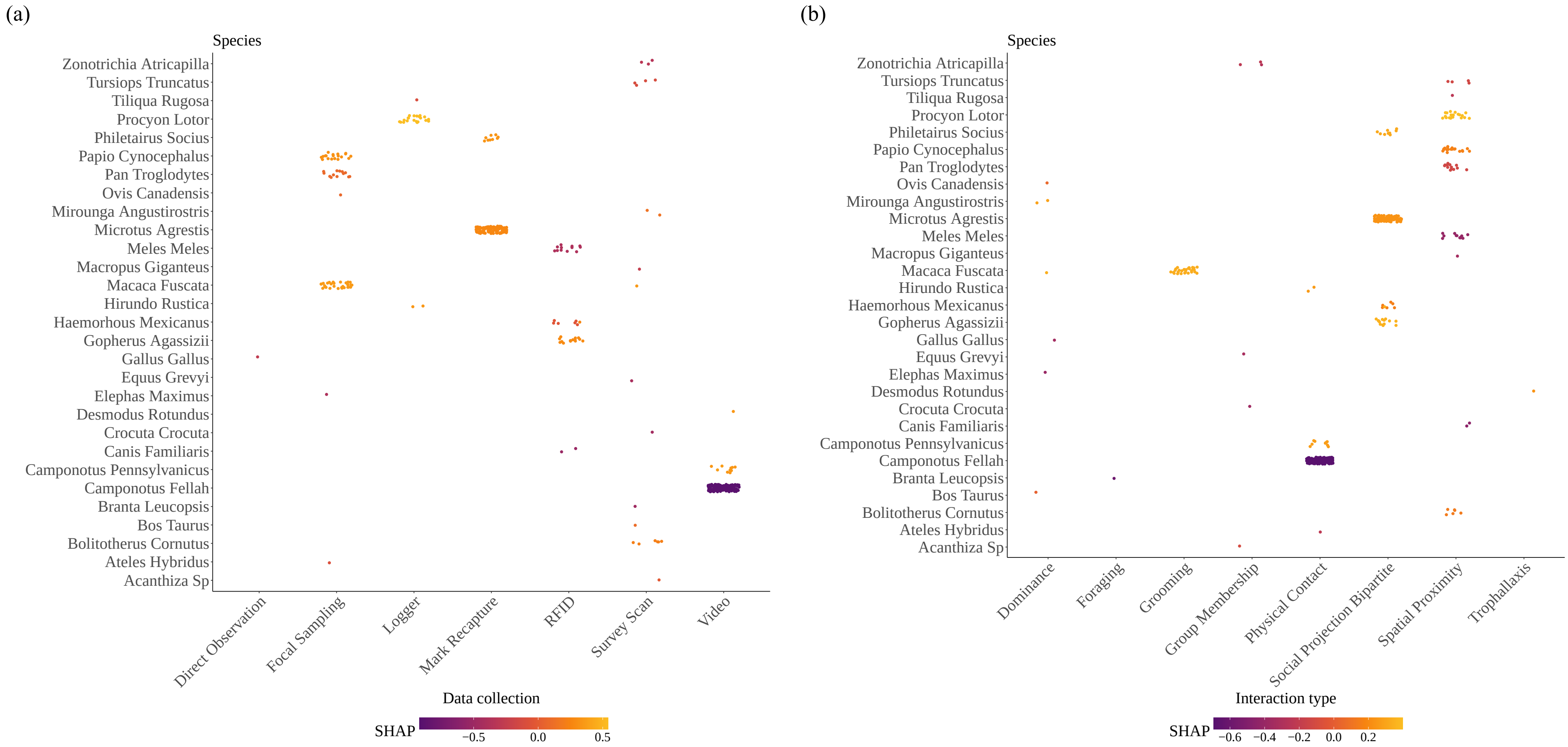
(a)
Species
Zonotrichia Atricapilla
Tursiops Truncatus
Tiliqua Rugosa
Procyon Lotor
Philetairus Socius
Papio Cynocephalus
Pan Troglodytes
Ovis Canadensis
Mirounga Angustirostris
Microtus Agrestis
Meles Meles
Macropus Giganteus
Macaca Fuscata
Hirundo Rustica
Haemorhous Mexicanus
Gopherus Agassizii
Gallus Gallus
Equus Grevyi
Elephas Maximus
Desmodus Rotundus
Crocuta Crocuta
Canis Familiaris
Camponotus Pennsylvanicus
Camponotus Fellah
Branta Leucopsis
Bos Taurus
Bolitotherus Cornutus
Ateles Hybridus
Acanthiza Sp
Direct Observation
Focal Sampling
Logger
Mark Recapture
RFID
Survey Scan
Video
Data collection
SHAP
−0.5
0.0
0.5
(b)
Species
Zonotrichia Atricapilla
Tursiops Truncatus
Tiliqua Rugosa
Procyon Lotor
Philetairus Socius
Papio Cynocephalus
Pan Troglodytes
Ovis Canadensis
Mirounga Angustirostris
Microtus Agrestis
Meles Meles
Macropus Giganteus
Macaca Fuscata
Hirundo Rustica
Haemorhous Mexicanus
Gopherus Agassizii
Gallus Gallus
Equus Grevyi
Elephas Maximus
Desmodus Rotundus
Crocuta Crocuta
Canis Familiaris
Camponotus Pennsylvanicus
Camponotus Fellah
Branta Leucopsis
Bos Taurus
Bolitotherus Cornutus
Ateles Hybridus
Acanthiza Sp
Dominance
Foraging
Grooming
Group Membership
Physical Contact
Social Projection Bipartite
Spatial Proximity
Trophallaxis
Interaction type
SHAP
−0.6
−0.4
−0.2
0.0
0.2

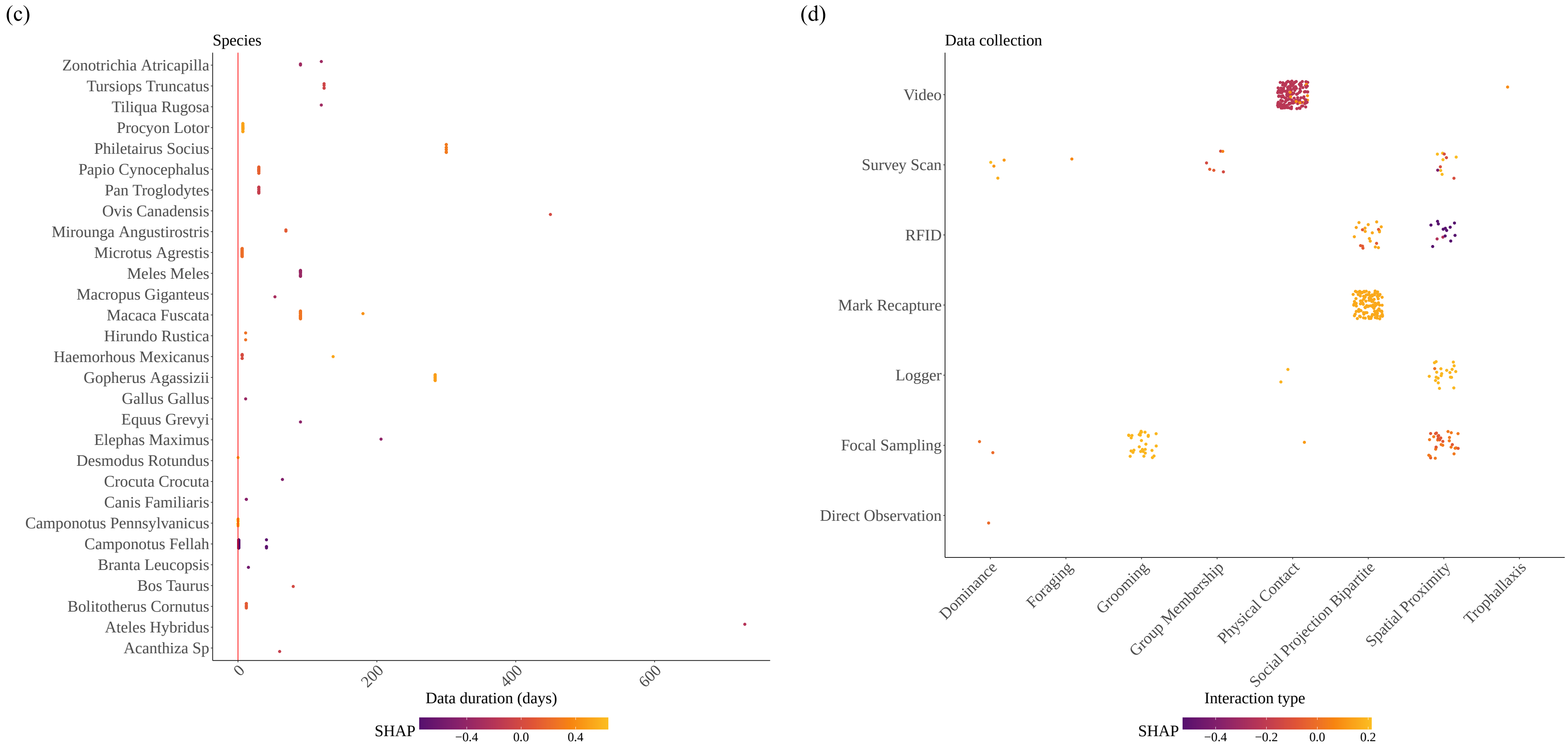
(c)
Species
Zonotrichia Atricapilla
Tursiops Truncatus
Tiliqua Rugosa
Procyon Lotor
Philetairus Socius
Papio Cynocephalus
Pan Troglodytes
Ovis Canadensis
Mirounga Angustirostris
Microtus Agrestis
Meles Meles
Macropus Giganteus
Macaca Fuscata
Hirundo Rustica
Haemorhous Mexicanus
Gopherus Agassizii
Gallus Gallus
Equus Grevyi
Elephas Maximus
Desmodus Rotundus
Crocuta Crocuta
Canis Familiaris
Camponotus Pennsylvanicus
Camponotus Fellah
Branta Leucopsis
Bos Taurus
Bolitotherus Cornutus
Ateles Hybridus
Acanthiza Sp
0
200
400
600
Data duration (days)
SHAP
−0.4
0.0
0.4
(d)
Data collection
Video
Survey Scan
RFID
Mark Recapture
Logger
Focal Sampling
Direct Observation
Dominance
Foraging
Grooming
Group Membership
Physical Contact
Social Projection Bipartite
Spatial Proximity
Trophallaxis
Interaction type
SHAP
−0.4
−0.2
0.0
0.2

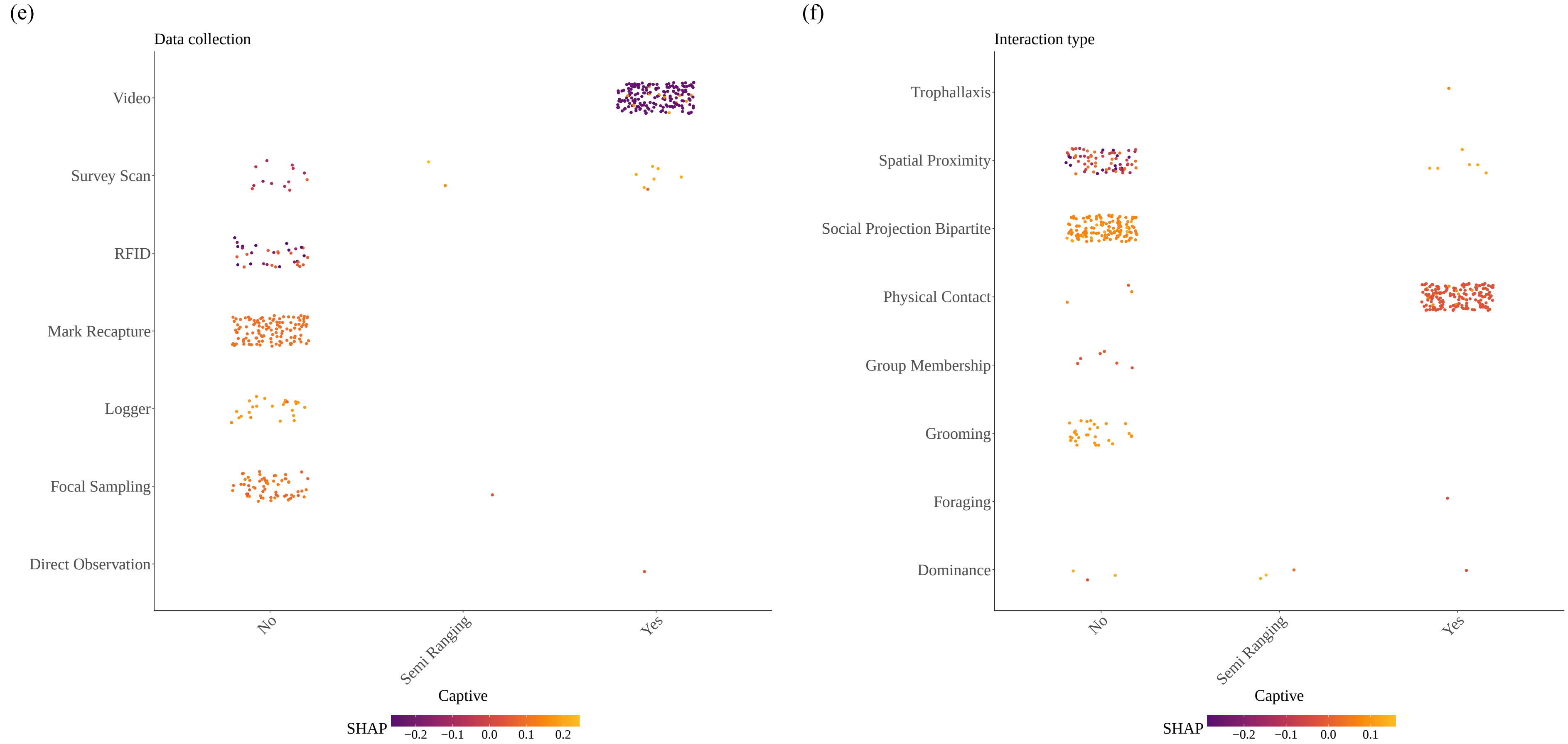


Fig. S10: This plot shows the pairwise interaction effect of (a) species and data collection, (b) species and interaction type, (c) species and data duration, (d) data collection and interaction type, (e) data collection and captive, and (f) interaction type and captive on the prediction of the scale-free class with species characteristics. This plot provides insights into how the model's prediction changes as different metadata interacts. Overall, this plot helps to understand the model's behavior with respect to metadata interaction.

Table S9: Summary of SBM network simulations with lower probability of between-community connection.

| k | Name | Pred | Order | Edges | M.Ecc | M.Path | Energy | Mod | Diam | Betw | Trans | Spec.R | Eig.C | Deg.C | M.Deg | MinCut | Fied | N.Fied | Close | Assort |
|---|---|---|---|---|---|---|---|---|---|---|---|---|---|---|---|---|---|---|---|---|
| 2 | sbm | SW | 100 | 1434 | 2.93 | 1.783 | 362.907 | 0.437 | 3 | 0.0146 | 0.4833 | 29.086 | 0.223 | 0.0739 | 28.68 | 18 | 3.367 | 0.118 | 0.0057 | -0.027 |
| 3 | sbm | Spatial | 100 | 1123 | 3 | 1.989 | 313.98 | 0.58 | 3 | 0.025 | 0.548 | 24.082 | 0.615 | 0.127 | 22.46 | 15 | 1.958 | 0.093 | 0.0051 | 0.221 |
| 4 | sbm | Spatial | 100 | 893 | 3 | 2.129 | 283.88 | 0.643 | 3 | 0.0297 | 0.553 | 19.561 | 0.664 | 0.072 | 17.86 | 9 | 1.921 | 0.103 | 0.0048 | 0.327 |
| 5 | sbm | SW | 100 | 731 | 3.16 | 2.251 | 267.794 | 0.648 | 4 | 0.044 | 0.496 | 15.658 | 0.468 | 0.064 | 14.62 | 7 | 1.553 | 0.126 | 0.0045 | 0.235 |
| 6 | sbm | SW | 100 | 623 | 3.42 | 2.352 | 252.062 | 0.650 | 4 | 0.052 | 0.467 | 14.755 | 0.719 | 0.116 | 12.46 | 5 | 1.554 | 0.139 | 0.0043 | 0.305 |
| 2 | sbm | Spatial | 150 | 3797 | 2.893 | 1.730 | 634.82 | 0.467 | 3 | 0.014 | 0.611 | 51.59 | 0.474 | 0.084 | 49.43 | 36 | 3.007 | 0.062 | 0.003 | 0.276 |
| 3 | sbm | SW | 150 | 2524 | 2.893 | 1.913 | 557.32 | 0.593 | 3 | 0.011 | 0.507 | 34.49 | 0.380 | 0.056 | 33.65 | 24 | 3.114 | 0.094 | 0.003 | 0.138 |
| 4 | sbm | Spatial | 150 | 1941 | 3.000 | 2.108 | 499.83 | 0.657 | 3 | 0.029 | 0.533 | 27.193 | 0.529 | 0.081 | 25.88 | 18 | 2.026 | 0.086 | 0.002 | 0.247 |
| 5 | sbm | SW | 150 | 1595 | 3.000 | 2.193 | 472.49 | 0.678 | 3 | 0.016 | 0.477 | 21.877 | 0.340 | 0.032 | 21.267 | 16 | 2.215 | 0.103 | 0.0031 | -0.021 |
| 6 | sbm | SW | 150 | 1344 | 3.013 | 2.277 | 444.41 | 0.677 | 4 | 0.014 | 0.464 | 18.740 | 0.469 | 0.040 | 17.92 | 10 | 2.373 | 0.129 | 0.003 | 0.154 |
| 2 | sbm | SW | 200 | 6820 | 2.780 | 1.689 | 958.26 | 0.471 | 3 | 0.009 | 0.630 | 68.517 | 0.293 | 0.079 | 68.2 | 52 | 3.826 | 0.057 | 0.003 | 0.001 |
| 3 | sbm | Spatial | 200 | 4158 | 3.000 | 1.954 | 829.87 | 0.608 | 3 | 0.010 | 0.521 | 43.299 | 0.505 | 0.097 | 41.58 | 28 | 2.665 | 0.065 | 0.003 | 0.304 |
| 4 | sbm | SW | 200 | 3226 | 3.000 | 2.043 | 762.39 | 0.645 | 3 | 0.010 | 0.485 | 33.716 | 0.363 | 0.050 | 32.26 | 23 | 3.115 | 0.101 | 0.002 | 0.177 |
| 5 | sbm | SW | 200 | 2743 | 3.000 | 2.067 | 736.94 | 0.626 | 3 | 0.009 | 0.413 | 28.440 | 0.426 | 0.023 | 27.43 | 19 | 3.601 | 0.134 | 0.002 | 0.087 |
| 6 | sbm | SW | 200 | 2345 | 3.000 | 2.171 | 683.40 | 0.648 | 3 | 0.007 | 0.402 | 24.344 | 0.352 | 0.048 | 23.45 | 13 | 3.793 | 0.162 | 0.002 | 0.096 |
| 2 | sbm | SW | 250 | 9258 | 2.652 | 1.715 | 1359.07 | 0.456 | 3 | 0.004 | 0.629 | 74.532 | 0.256 | 0.064 | 74.064 | 56 | 6.331 | 0.086 | 0.002 | -0.010 |
| 3 | sbm | Spatial | 250 | 7790 | 2.844 | 1.812 | 1118.87 | 0.608 | 3 | 0.006 | 0.661 | 63.821 | 0.330 | 0.023 | 62.32 | 44 | 4.199 | 0.069 | 0.002 | 0.447 |
| 4 | sbm | Spatial | 250 | 5933 | 2.996 | 1.929 | 1052.75 | 0.668 | 3 | 0.008 | 0.587 | 48.821 | 0.552 | 0.063 | 47.464 | 36 | 4.111 | 0.087 | 0.002 | 0.264 |
| 5 | sbm | SW | 250 | 4024 | 3.000 | 1.990 | 894.90 | 0.668 | 3 | 0.007 | 0.531 | 39.405 | 0.508 | 0.083 | 32.192 | 26 | 4.334 | 0.109 | 0.002 | 0.142 |
| 6 | sbm | SW | 250 | 3404 | 3.000 | 2.102 | 815.20 | 0.676 | 3 | 0.007 | 0.478 | 34.984 | 0.289 | 0.038 | 31.96 | 18 | 4.239 | 0.133 | 0.002 | 0.334 |
| 2 | sbm | Spatial | 300 | 17315 | 2.263 | 1.712 | 1762.09 | 0.475 | 3 | 0.004 | 0.717 | 117.826 | 0.405 | 0.073 | 115.433 | 86 | 6.520 | 0.056 | 0.002 | 0.345 |
| 3 | sbm | Spatial | 300 | 10115 | 2.806 | 1.839 | 1452.45 | 0.608 | 3 | 0.005 | 0.638 | 71.711 | 0.446 | 0.042 | 67.433 | 49 | 6.703 | 0.098 | 0.002 | 0.438 |
| 4 | sbm | SW | 300 | 7734 | 3.000 | 1.929 | 1371.22 | 0.642 | 3 | 0.007 | 0.603 | 54.142 | 0.530 | 0.083 | 51.56 | 34 | 4.316 | 0.085 | 0.002 | 0.373 |
| 5 | sbm | SW | 300 | 6320 | 3.000 | 1.977 | 1278.87 | 0.689 | 3 | 0.004 | 0.581 | 43.730 | 0.476 | 0.043 | 42.133 | 27 | 4.981 | 0.114 | 0.002 | 0.356 |
| 6 | sbm | SW | 300 | 5684 | 3.000 | 2.065 | 1170.35 | 0.687 | 3 | 0.005 | 0.476 | 40.863 | 0.684 | 0.048 | 37.893 | 20 | 4.884 | 0.124 | 0.002 | 0.345 |
| 2 | sbm | Spatial | 350 | 19378 | 2.291 | 1.684 | 2124.54 | 0.509 | 3 | 0.004 | 0.731 | 122.753 | 0.507 | 0.067 | 110.731 | 80 | 6.935 | 0.078 | 0.002 | 0.777 |
| 3 | sbm | Spatial | 350 | 13923 | 2.903 | 1.795 | 1898.71 | 0.579 | 3 | 0.003 | 0.635 | 87.823 | 0.549 | 0.087 | 79.56 | 45 | 6.994 | 0.098 | 0.002 | 0.712 |
| 4 | sbm | SW | 350 | 10803 | 2.873 | 1.879 | 1730.53 | 0.630 | 3 | 0.003 | 0.616 | 68.991 | 0.546 | 0.087 | 61.731 | 42 | 5.973 | 0.094 | 0.002 | 0.484 |
| 5 | sbm | SW | 350 | 9159 | 3.000 | 1.947 | 1597.08 | 0.663 | 3 | 0.005 | 0.527 | 57.293 | 0.621 | 0.052 | 52.337 | 31 | 6.398 | 0.134 | 0.002 | 0.602 |
| 6 | sbm | SW | 350 | 7739 | 3.000 | 2.014 | 1510.94 | 0.637 | 3 | 0.004 | 0.495 | 47.474 | 0.638 | 0.042 | 44.223 | 26 | 6.146 | 0.150 | 0.002 | 0.423 |
| 2 | sbm | sbm | 400 | 24439 | 2.058 | 1.694 | 2711.32 | 0.440 | 3 | 0.002 | 0.538 | 130.470 | 0.450 | 0.082 | 122.195 | 88 | 13.106 | 0.108 | 0.002 | 0.059 |
| 3 | sbm | Spatial | 400 | 19870 | 2.173 | 1.751 | 2392.03 | 0.548 | 3 | 0.003 | 0.682 | 106.565 | 0.513 | 0.093 | 99.35 | 67 | 13.321 | 0.132 | 0.002 | 0.680 |
| 4 | sbm | Spatial | 400 | 15614 | 2.850 | 1.843 | 2064.10 | 0.636 | 3 | 0.003 | 0.636 | 83.180 | 0.502 | 0.097 | 78.07 | 50 | 7.664 | 0.107 | 0.002 | 0.644 |
| 5 | sbm | SW | 400 | 13182 | 2.993 | 1.887 | 1937.81 | 0.637 | 3 | 0.003 | 0.569 | 68.891 | 0.523 | 0.003 | 65.91 | 40 | 7.981 | 0.137 | 0.002 | 0.523 |
| 6 | sbm | SW | 400 | 11293 | 2.998 | 1.921 | 1804.53 | 0.682 | 3 | 0.003 | 0.528 | 58.295 | 0.377 | 0.039 | 56.465 | 33 | 8.627 | 0.150 | 0.002 | 0.373 |
| 2 | sbm | sbm | 450 | 33441 | 2.017 | 1.686 | 3221.16 | 0.435 | 3 | 0.002 | 0.764 | 158.385 | 0.443 | 0.087 | 148.627 | 112 | 17.149 | 0.118 | 0.002 | 0.715 |
| 3 | sbm | Spatial | 450 | 23999 | 2.836 | 1.783 | 2813.97 | 0.611 | 3 | 0.003 | 0.628 | 117.748 | 0.643 | 0.058 | 106.662 | 74 | 6.081 | 0.053 | 0.002 | 0.724 |
| 4 | sbm | Spatial | 450 | 17968 | 2.902 | 1.865 | 2404.44 | 0.643 | 3 | 0.003 | 0.603 | 90.787 | 0.576 | 0.045 | 79.857 | 50 | 6.220 | 0.094 | 0.002 | 0.584 |
| 5 | sbm | Spatial | 450 | 14423 | 2.987 | 1.930 | 2286.22 | 0.667 | 3 | 0.003 | 0.495 | 72.030 | 0.690 | 0.072 | 63.658 | 44 | 7.585 | 0.126 | 0.002 | 0.526 |
| 6 | sbm | SW | 450 | 12530 | 3.000 | 1.978 | 2114.17 | 0.678 | 3 | 0.003 | 0.476 | 63.852 | 0.687 | 0.032 | 55.689 | 34 | 7.918 | 0.137 | 0.002 | 0.364 |
| 2 | sbm | Spatial | 500 | 45587 | 2.066 | 1.654 | 3482.72 | 0.467 | 3 | 0.001 | 0.687 | 197.375 | 0.478 | 0.075 | 182.348 | 145 | 10.347 | 0.057 | 0.002 | 0.531 |
| 3 | sbm | sbm | 500 | 30985 | 2.168 | 1.722 | 3119.11 | 0.537 | 3 | 0.001 | 0.652 | 134.893 | 0.272 | 0.042 | 123.94 | 103 | 13.226 | 0.108 | 0.002 | 0.123 |
| 4 | sbm | SW | 500 | 24224 | 2.996 | 1.910 | 2822.98 | 0.633 | 3 | 0.001 | 0.547 | 97.917 | 0.289 | 0.026 | 96.896 | 75 | 12.865 | 0.133 | 0.002 | 0.140 |
| 5 | sbm | SW | 500 | 19769 | 3.000 | 1.966 | 2744.90 | 0.653 | 3 | 0.002 | 0.508 | 79.720 | 0.313 | 0.024 | 79.076 | 62 | 10.482 | 0.133 | 0.002 | 0.078 |
| 6 | sbm | Spatial | 500 | 17253 | 2.964 | 1.997 | 2347.44 | 0.663 | 3 | 0.003 | 0.456 | 70.747 | 0.484 | 0.047 | 69.012 | 51 | 10.446 | 0.154 | 0.002 | 0.220 |
| 2 | sbm | sbm | 550 | 54031 | 2.000 | 1.642 | 4182.95 | 0.457 | 3 | 0.001 | 0.632 | 198.576 | 0.299 | 0.067 | 196.476 | 167 | 16.326 | 0.082 | 0.002 | 0.319 |
| 3 | sbm | Spatial | 550 | 34644 | 2.745 | 1.758 | 3381.13 | 0.578 | 3 | 0.002 | 0.637 | 146.213 | 0.399 | 0.043 | 126.524 | 89 | 13.731 | 0.108 | 0.002 | 0.571 |
| 4 | sbm | Spatial | 550 | 28197 | 2.794 | 1.839 | 3222.19 | 0.648 | 3 | 0.002 | 0.608 | 116.114 | 0.434 | 0.052 | 102.535 | 86 | 9.805 | 0.095 | 0.002 | 0.716 |
| 5 | sbm | Spatial | 550 | 23560 | 2.929 | 1.903 | 3039.03 | 0.659 | 3 | 0.002 | 0.550 | 90.977 | 0.390 | 0.043 | 85.673 | 51 | 9.108 | 0.110 | 0.002 | 0.560 |
| 6 | sbm | Spatial | 550 | 20074 | 2.995 | 1.923 | 2815.06 | 0.672 | 3 | 0.002 | 0.507 | 77.828 | 0.488 | 0.083 | 72.997 | 41 | 10.473 | 0.145 | 0.002 | 0.582 |
| 2 | sbm | Spatial | 600 | 65325 | 2.535 | 1.642 | 4462.55 | 0.494 | 3 | 0.002 | 0.703 | 226.676 | 0.409 | 0.029 | 217.75 | 177 | 6.072 | 0.027 | 0.002 | 0.760 |
| 3 | sbm | Spatial | 600 | 45668 | 2.007 | 1.745 | 4147.2 | 0.565 | 3 | 0.001 | 0.609 | 161.712 | 0.425 | 0.047 | 152.227 | 103 | 20.339 | 0.123 | 0.002 | 0.720 |
| 4 | sbm | Spatial | 600 | 34988 | 2.833 | 1.813 | 3754.95 | 0.634 | 3 | 0.001 | 0.573 | 123.336 | 0.415 | 0.037 | 116.627 | 72 | 12.163 | 0.101 | 0.002 | 0.673 |
| 5 | sbm | Spatial | 600 | 29495 | 2.920 | 1.854 | 3454.84 | 0.662 | 3 | 0.002 | 0.529 | 103.742 | 0.370 | 0.058 | 98.317 | 59 | 11.770 | 0.136 | 0.002 | 0.667 |
| 6 | sbm | SW | 600 | 25223 | 2.999 | 1.899 | 3047.10 | 0.684 | 3 | 0.002 | 0.526 | 88.293 | 0.370 | 0.043 | 84.077 | 48 | 11.596 | 0.142 | 0.002 | 0.583 |
| 2 | sbm | Spatial | 650 | 81754 | 2.153 | 1.613 | 4875.12 | 0.481 | 3 | 0.002 | 0.735 | 254.640 | 0.378 | 0.043 | 251.551 | 218 | 9.347 | 0.037 | 0.002 | 0.307 |
| 3 | sbm | Spatial | 650 | 51234 | 2.409 | 1.751 | 4475.49 | 0.608 | 3 | 0.002 | 0.621 | 165.316 | 0.546 | 0.063 | 157.643 | 120 | 10.550 | 0.069 | 0.002 | 0.684 |
| 4 | sbm | Spatial | 650 | 39156 | 2.633 | 1.823 | 4176.28 | 0.644 | 3 | 0.002 | 0.593 | 126.949 | 0.534 | 0.059 | 120.482 | 89 | 11.419 | 0.097 | 0.002 | 0.524 |
| 5 | sbm | Spatial | 650 | 32038 | 2.848 | 1.878 | 3897.19 | 0.678 | 3 | 0.002 | 0.520 | 103.108 | 0.565 | 0.040 | 98.732 | 73 | 11.802 | 0.126 | 0.002 | 0.397 |
| 6 | sbm | Spatial | 650 | 27230 | 2.978 | 1.897 | 3377.73 | 0.660 | 3 | 0.002 | 0.482 | 87.207 | 0.512 | 0.038 | 83.785 | 51 | 13.024 | 0.146 | 0.002 | 0.466 |
| 2 | sbm | sbm | 700 | 91435 | 2.008 | 1.627 | 5275.38 | 0.474 | 3 | 0.002 | 0.692 | 265.101 | 0.379 | 0.047 | 261.243 | 220 | 34.588 | 0.158 | 0.002 | 0.478 |
| 3 | sbm | Spatial | 700 | 65706 | 2.329 | 1.733 | 4855.73 | 0.607 | 3 | 0.001 | 0.576 | 192.189 | 0.533 | 0.047 | 187.731 | 153 | 12.535 | 0.068 | 0.002 | 0.408 |
| 4 | sbm | Spatial | 700 | 50164 | 2.457 | 1.796 | 4544.63 | 0.635 | 3 | 0.001 | 0.617 | 148.975 | 0.310 | 0.043 | 143.326 | 117 | 13.244 | 0.093 | 0.002 | 0.394 |
| 5 | sbm | SW | 700 | 40685 | 2.814 | 1.844 | 4375.18 | 0.679 | 3 | 0.001 | 0.579 | 117.182 | 0.312 | 0.037 | 116.243 | 91 | 14.319 | 0.120 | 0.002 | 0.245 |
| 6 | sbm | Spatial | 700 | 33164 | 2.957 | 1.897 | 3742.14 | 0.682 | 3 | 0.002 | 0.533 | 99.747 | 0.534 | 0.043 | 94.754 | 57 | 12.437 | 0.132 | 0.002 | 0.617 |

| k | Name | Pred | Order | Edges | M.Ecc | M.Path | Energy | Mod | Diam | Betw | Trans | Spec.R | Eig.C | Deg.C | M.Deg | MinCut | Fied | N.Fied | Close | Assort |
|---|---|---|---|---|---|---|---|---|---|---|---|---|---|---|---|---|---|---|---|---|
| 2 | sbm | sbm | 750 | 113277 | 2.000 | 1.597 | 6156.34 | 0.469 | 3 | 0.001 | 0.716 | 302.450 | 0.135 | 0.047 | 302.072 | 277 | 24.172 | 0.081 | 0.002 | 0.033 |
| 3 | sbm | Spatial | 750 | 68366 | 2.438 | 1.758 | 5541.24 | 0.632 | 3 | 0.001 | 0.642 | 193.262 | 0.607 | 0.043 | 182.309 | 152 | 10.685 | 0.057 | 0.002 | 0.696 |
| 4 | sbm | Spatial | 750 | 50907 | 2.638 | 1.823 | 5132.21 | 0.649 | 3 | 0.001 | 0.587 | 146.263 | 0.620 | 0.045 | 135.752 | 94 | 13.967 | 0.103 | 0.002 | 0.653 |
| 5 | sbm | Spatial | 750 | 41611 | 2.944 | 1.866 | 4615.42 | 0.677 | 3 | 0.002 | 0.517 | 117.903 | 0.633 | 0.043 | 110.963 | 77 | 12.442 | 0.102 | 0.002 | 0.577 |
| 6 | sbm | Spatial | 750 | 35250 | 2.961 | 1.896 | 4610.62 | 0.663 | 3 | 0.001 | 0.486 | 100.214 | 0.661 | 0.043 | 94.267 | 67 | 13.942 | 0.126 | 0.002 | 0.494 |
| 2 | sbm | sbm | 800 | 117532 | 2.000 | 1.623 | 7002.70 | 0.466 | 2 | 0.001 | 0.671 | 300.77 | 0.445 | 0.047 | 293.83 | 242 | 31.316 | 0.107 | 0.002 | 0.096 |
| 3 | sbm | Spatial | 800 | 78667 | 2.036 | 1.754 | 6055.452 | 0.586 | 3 | 0.0010 | 0.636 | 213.78 | 0.480 | 0.054 | 196.67 | 121 | 17.514 | 0.080 | 0.0007 | 0.796 |
| 4 | sbm | Spatial | 800 | 60956 | 2.668 | 1.815 | 5466.84 | 0.656 | 3 | 0.001 | 0.615 | 152.39 | 0.493 | 0.042 | 152.39 | 79 | 15.963 | 0.103 | 0.002 | 0.790 |
| 5 | sbm | Spatial | 800 | 49001 | 2.938 | 1.870 | 5221.97 | 0.669 | 3 | 0.001 | 0.483 | 129.701 | 0.642 | 0.043 | 122.503 | 75 | 14.945 | 0.114 | 0.002 | 0.573 |
| 6 | sbm | Spatial | 800 | 42763 | 2.939 | 1.881 | 4610.51 | 0.672 | 3 | 0.001 | 0.512 | 114.294 | 0.543 | 0.043 | 106.908 | 66 | 15.705 | 0.150 | 0.002 | 0.645 |
| 2 | sbm | sbm | 850 | 121251 | 2.000 | 1.623 | 7396.35 | 0.432 | 2 | 0.001 | 0.629 | 294.219 | 0.647 | 0.051 | 285.297 | 201 | 35.289 | 0.118 | 0.002 | 0.047 |
| 3 | sbm | Spatial | 850 | 86245 | 2.592 | 1.765 | 5956.49 | 0.595 | 3 | 0.001 | 0.631 | 227.649 | 0.520 | 0.047 | 202.929 | 131 | 11.293 | 0.057 | 0.002 | 0.872 |
| 4 | sbm | Spatial | 850 | 66489 | 2.741 | 1.819 | 5816.65 | 0.672 | 3 | 0.001 | 0.631 | 173.915 | 0.467 | 0.043 | 156.444 | 101 | 12.602 | 0.074 | 0.002 | 0.783 |
| 5 | sbm | Spatial | 850 | 53740 | 2.989 | 1.880 | 5454.31 | 0.670 | 3 | 0.001 | 0.592 | 141.117 | 0.001 | 0.047 | 126.447 | 79 | 10.671 | 0.083 | 0.002 | 0.763 |
| 6 | sbm | Spatial | 850 | 45101 | 2.937 | 1.909 | 5511.18 | 0.698 | 3 | 0.001 | 0.532 | 118.622 | 0.544 | 0.043 | 106.12 | 67 | 13.276 | 0.124 | 0.002 | 0.744 |
| 2 | sbm | Spatial | 900 | 154775 | 2.054 | 1.618 | 7841.67 | 0.483 | 3 | 0.001 | 0.733 | 353.086 | 0.449 | 0.043 | 343.944 | 306 | 10.596 | 0.031 | 0.002 | 0.615 |
| 3 | sbm | Spatial | 900 | 96462 | 2.631 | 1.763 | 7032.59 | 0.623 | 3 | 0.001 | 0.645 | 227.083 | 0.479 | 0.043 | 214.36 | 159 | 13.773 | 0.069 | 0.002 | 0.818 |
| 4 | sbm | Spatial | 900 | 70012 | 2.976 | 1.845 | 6584.31 | 0.689 | 3 | 0.002 | 0.605 | 170.406 | 0.567 | 0.047 | 155.582 | 109 | 10.537 | 0.070 | 0.002 | 0.733 |
| 5 | sbm | Spatial | 900 | 57005 | 2.997 | 1.886 | 6071.18 | 0.698 | 3 | 0.001 | 0.542 | 137.931 | 0.601 | 0.047 | 126.678 | 87 | 11.184 | 0.087 | 0.002 | 0.709 |
| 6 | sbm | Spatial | 900 | 48461 | 2.984 | 1.901 | 5398.36 | 0.683 | 3 | 0.001 | 0.483 | 115.773 | 0.592 | 0.038 | 107.691 | 76 | 12.549 | 0.103 | 0.002 | 0.621 |
| 2 | sbm | sbm | 950 | 170835 | 2.003 | 1.602 | 8760.57 | 0.475 | 3 | 0.001 | 0.583 | 361.761 | 0.358 | 0.040 | 359.653 | 325 | 17.452 | 0.049 | 0.002 | 0.135 |
| 3 | sbm | sbm | 950 | 96781 | 2.057 | 1.683 | 7635.71 | 0.579 | 3 | 0.001 | 0.520 | 220.791 | 0.565 | 0.047 | 203.749 | 156 | 20.587 | 0.096 | 0.002 | 0.729 |
| 4 | sbm | Spatial | 950 | 78657 | 2.846 | 1.837 | 7128.46 | 0.616 | 3 | 0.001 | 0.541 | 183.137 | 0.655 | 0.047 | 165.594 | 117 | 13.449 | 0.078 | 0.002 | 0.724 |
| 5 | sbm | Spatial | 950 | 65344 | 2.865 | 1.865 | 6697.52 | 0.679 | 3 | 0.001 | 0.523 | 148.137 | 0.634 | 0.047 | 137.566 | 93 | 15.634 | 0.106 | 0.002 | 0.696 |
| 6 | sbm | sbm | 950 | 56714 | 2.832 | 1.862 | 6324.09 | 0.682 | 3 | 0.001 | 0.487 | 127.91 | 0.602 | 0.047 | 119.398 | 81 | 17.773 | 0.153 | 0.002 | 0.584 |
| 2 | sbm | sbm | 1000 | 142265 | 2.000 | 1.715 | 10044.26 | 0.449 | 3 | 0.001 | 0.623 | 298.311 | 0.262 | 0.047 | 284.53 | 238 | 28.660 | 0.101 | 0.002 | 0.197 |
| 3 | sbm | Spatial | 1000 | 116556 | 2.466 | 1.796 | 8267.34 | 0.649 | 3 | 0.001 | 0.584 | 242.791 | 0.497 | 0.047 | 233.11 | 188 | 12.977 | 0.056 | 0.002 | 0.695 |
| 4 | sbm | Spatial | 1000 | 95054 | 2.531 | 1.812 | 7728.61 | 0.653 | 3 | 0.001 | 0.584 | 202.286 | 0.579 | 0.047 | 159.106 | 151 | 15.926 | 0.089 | 0.002 | 0.712 |
| 5 | sbm | Spatial | 1000 | 76224 | 2.621 | 1.832 | 7270.23 | 0.661 | 3 | 0.001 | 0.521 | 163.912 | 0.613 | 0.047 | 152.448 | 107 | 20.303 | 0.128 | 0.002 | 0.660 |
| 6 | sbm | Spatial | 1000 | 63486 | 2.633 | 1.885 | 7033.92 | 0.589 | 3 | 0.001 | 0.489 | 136.912 | 0.838 | 0.001 | 128.972 | 82 | 19.445 | 0.147 | 0.001 | 0.848 |

k = communities; Pred = predicted class; M.Ecc = mean eccentricity; M.Path = mean path length; Mod = modularity; Diam = diameter; Betw = betweenness centrality; Trans = transitivity; Spec.R = spectral radius; Eig.C = eigenvector centrality; Deg.C = degree centrality; M.Deg = mean degree; Fied = Fiedler value; N.Fied = normalized Fiedler value; Close = closeness centrality; Assort = degree assortativity coefficient; SW = small-world.

Table S10: Summary of SBM network simulations with higher probability of between-community connection.

| k | Name | Pred | Order | Edges | M.Ecc | M.Path | Energy | Mod | Diam | Betw | Trans | Spec.R | Eig.C | Deg.C | M.Deg | MinCut | Fied | N.Fied | Close | Assort |
|---|---|---|---|---|---|---|---|---|---|---|---|---|---|---|---|---|---|---|---|---|
| 2 | sbm | sbm | 100 | 2397 | 2.1 | 1.51602 | 428.5896 | 0.25523 | 2 | 0.00433 | 0.55808 | 48.78933 | 0.24722 | 0.11177 | 47.94 | 34 | 21.99272 | 0.47366 | 0.00673 | 0.16501 |
| 3 | sbm | sbm | 100 | 1950 | 2 | 1.60601 | 436.3823 | 0.16078 | 2 | 0.00744 | 0.4338 | 40.3062 | 0.35855 | 0.18181 | 39 | 27 | 20.03213 | 0.5821 | 0.00630 | 0.10232 |
| 4 | sbm | sbm | 100 | 1677 | 2.02 | 1.66143 | 409.8359 | 0.22235 | 3 | 0.00917 | 0.39586 | 34.79976 | 0.30976 | 0.14604 | 33.54 | 21 | 15.21726 | 0.50402 | 0.00607 | 0.15647 |
| 5 | sbm | SW | 100 | 1619 | 2 | 1.67329 | 412.7814 | 0.20121 | 2 | 0.00709 | 0.35867 | 33.1547 | 0.30862 | 0.13776 | 32.38 | 21 | 17.32963 | 0.58614 | 0.00643 | 0.01786 |
| 6 | sbm | SW | 100 | 1552 | 2.02 | 1.68667 | 413.3622 | 0.15906 | 3 | 0.00775 | 0.34802 | 32.06998 | 0.34002 | 0.13099 | 31.04 | 18 | 15.4669 | 0.58121 | 0.00595 | 0.05632 |
| 2 | sbm | sbm | 150 | 4619 | 2 | 1.58687 | 762.8837 | 0.27527 | 2 | 0.00375 | 0.48982 | 62.3975 | 0.25268 | 0.13157 | 61.58667 | 47 | 26.41986 | 0.45624 | 0.00433 | 0.13164 |
| 3 | sbm | SW | 150 | 3916 | 2 | 1.69557 | 762.104 | 0.26302 | 2 | 0.00314 | 0.40819 | 53.03859 | 0.37878 | 0.10951 | 52.21333 | 37 | 25.43206 | 0.52008 | 0.00471 | 0.05792 |
| 4 | sbm | SW | 150 | 3338 | 2.013 | 1.70138 | 720.6619 | 0.26853 | 3 | 0.00367 | 0.46205 | 45.0829 | 0.25165 | 0.08058 | 44.50667 | 28 | 21.83847 | 0.55952 | 0.00394 | 0.12226 |
| 5 | sbm | SW | 150 | 2485 | 2 | 1.68514 | 734.3975 | 0.17727 | 2 | 0.00547 | 0.34931 | 47.7289 | 0.33159 | 0.12768 | 44.56667 | 31 | 23.44745 | 0.57267 | 0.00379 | 0.07552 |
| 6 | sbm | SW | 150 | 3332 | 2 | 1.70183 | 732.4482 | 0.17484 | 2 | 0.00414 | 0.35043 | 45.6086 | 0.32449 | 0.12863 | 44.42667 | 31 | 23.6964 | 0.62969 | 0.00347 | 0.06834 |
| 2 | sbm | sbm | 200 | 8620 | 2 | 1.56684 | 1082.736 | 0.25673 | 2 | 0.00113 | 0.63192 | 86.61113 | 0.35528 | 0.08447 | 86.2 | 69 | 38.42023 | 0.45056 | 0.00326 | 0.07733 |
| 3 | sbm | sbm | 200 | 6564 | 2 | 1.67015 | 1127.435 | 0.26262 | 2 | 0.00244 | 0.37872 | 66.6794 | 0.36617 | 0.12787 | 65.84 | 46 | 31.57158 | 0.50967 | 0.00301 | 0.11819 |
| 4 | sbm | sbm | 200 | 6111 | 2 | 1.693 | 1121.80 | 0.212 | 2 | 0.003 | 0.344 | 62.738 | 0.283 | 0.105 | 61.11 | 35 | 29.0398 | 0.563 | 0.003 | 0.185 |
| 5 | sbm | sbm | 200 | 6632 | 2 | 1.667 | 1160.912 | 0.146 | 2 | 0.004 | 0.359 | 67.879 | 0.352 | 0.174 | 66.32 | 44 | 37.352 | 0.657 | 0.003 | 0.073 |
| 6 | sbm | SW | 200 | 6453 | 2 | 1.67579 | 1156.135 | 0.14524 | 2 | 0.00292 | 0.40755 | 65.0237 | 0.21259 | 0.02764 | 64.53 | 38 | 39.42136 | 0.57535 | 0.003 | 0.01639 |
| 2 | sbm | sbm | 250 | 12936 | 2 | 1.61708 | 1687.096 | 0.24182 | 2 | 0.00185 | 0.43778 | 103.2444 | 0.21751 | 0.08524 | 103.488 | 80 | 58.89462 | 0.50993 | 0.00251 | 0.01039 |
| 3 | sbm | sbm | 250 | 11984 | 2 | 1.61763 | 1572.607 | 0.28152 | 2 | 0.00164 | 0.64138 | 96.1183 | 0.31189 | 0.10307 | 95.152 | 66 | 42.60721 | 0.47568 | 0.00284 | 0.22909 |
| 4 | sbm | sbm | 250 | 10792 | 2 | 1.65326 | 1564.13 | 0.28152 | 2 | 0.00121 | 0.46484 | 87.3197 | 0.20915 | 0.06704 | 86.336 | 53 | 38.53818 | 0.56811 | 0.00294 | 0.10807 |
| 5 | sbm | sbm | 250 | 10267 | 2 | 1.67013 | 1579.064 | 0.21014 | 2 | 0.00204 | 0.98451 | 83.0789 | 0.25558 | 0.09855 | 82.136 | 61 | 42.8628 | 0.56901 | 0.00249 | 0.12807 |
| 6 | sbm | sbm | 250 | 9277 | 2 | 1.70194 | 1537.118 | 0.21014 | 2 | 0.00174 | 0.3582 | 75.7210 | 0.34518 | 0.10756 | 74.216 | 53 | 39.8532 | 0.56811 | 0.00261 | 0.10807 |
| 2 | sbm | sbm | 300 | 22180 | 2 | 1.50545 | 2107.292 | 0.25523 | 2 | 0.0072 | 0.56296 | 148.4721 | 0.17523 | 0.00747 | 147.8667 | 122 | 70.5991 | 0.48112 | 0.00222 | 0.10652 |
| 3 | sbm | sbm | 300 | 17645 | 2 | 1.60867 | 2165.102 | 0.18434 | 2 | 0.00183 | 0.42339 | 118.3689 | 0.34515 | 0.09214 | 117.6333 | 90 | 64.26022 | 0.55634 | 0.00202 | 0.21067 |
| 4 | sbm | sbm | 300 | 15090 | 2 | 1.65376 | 2069.158 | 0.20609 | 2 | 0.00197 | 0.39748 | 103.3969 | 0.29731 | 0.12192 | 100.5333 | 66 | 49.2815 | 0.50613 | 0.00212 | 0.21067 |
| 5 | sbm | sbm | 300 | 14106 | 2 | 1.68548 | 2042.118 | 0.21755 | 2 | 0.00121 | 0.39385 | 95.3634 | 0.34828 | 0.08437 | 94.04 | 66 | 49.6925 | 0.55634 | 0.00185 | 0.10552 |
| 6 | sbm | sbm | 300 | 13578 | 2 | 1.69758 | 2017.642 | 0.21455 | 2 | 0.00194 | 0.34294 | 92.3067 | 0.28789 | 0.10829 | 90.52 | 64 | 47.7607 | 0.56667 | 0.00197 | 0.13785 |
| 2 | sbm | sbm | 350 | 26295 | 2 | 1.56964 | 2709.222 | 0.26853 | 2 | 0.00765 | 0.47066 | 153.1949 | 0.26336 | 0.11011 | 150.2571 | 114 | 84.7566 | 0.57248 | 0.00167 | 0.33851 |
| 3 | sbm | sbm | 350 | 23725 | 2 | 1.61142 | 2676.683 | 0.18577 | 2 | 0.00163 | 0.44243 | 140.7781 | 0.28651 | 0.12702 | 135.5714 | 85 | 67.7067 | 0.58113 | 0.00178 | 0.33851 |
| 4 | sbm | sbm | 350 | 22153 | 2 | 1.63725 | 2366.006 | 0.18677 | 2 | 0.00127 | 0.40183 | 135.7775 | 0.23159 | 0.10146 | 126.5886 | 90 | 58.4729 | 0.53602 | 0.00175 | 0.21258 |
| 5 | sbm | sbm | 350 | 19716 | 2 | 1.67718 | 2565.946 | 0.20356 | 2 | 0.00123 | 0.36731 | 115.5748 | 0.25063 | 0.19411 | 112.6629 | 71 | 61.5795 | 0.63051 | 0.0017 | 0.15086 |
| 6 | sbm | sbm | 350 | 15243 | 2 | 1.68492 | 2366.653 | 0.17755 | 2 | 0.0017 | 0.34507 | 131.907 | 0.24682 | 0.11793 | 87.1029 | 79 | 64.74134 | 0.63051 | 0.00170 | 0.06717 |
| 2 | sbm | sbm | 400 | 36432 | 2 | 1.54393 | 3469.006 | 0.1314 | 2 | 0.00485 | 0.47267 | 183.9299 | 0.19223 | 0.04437 | 182.16 | 132 | 128.1974 | 0.72479 | 0.00162 | 0.16591 |
| 3 | sbm | sbm | 400 | 26614 | 2 | 1.53087 | 2961.112 | 0.15307 | 2 | 0.00407 | 0.40898 | 165.754 | 0.31422 | 0.06503 | 133.07 | 141 | 129.015 | 0.75807 | 0.00163 | 0.13962 |
| 4 | sbm | sbm | 400 | 31011 | 2 | 1.61135 | 3244.894 | 0.20778 | 2 | 0.00101 | 0.34005 | 157.989 | 0.24142 | 0.1101 | 155.07 | 113 | 78.5879 | 0.54893 | 0.00155 | 0.12717 |
| 5 | sbm | sbm | 400 | 20007 | 2 | 1.62397 | 2564.838 | 0.16585 | 2 | 0.0038 | 0.41544 | 154.982 | 0.24542 | 0.12013 | 100.035 | 107 | 70.0608 | 0.59156 | 0.00154 | 0.10552 |
| 6 | sbm | sbm | 400 | 27924 | 2 | 1.66007 | 3226.277 | 0.16738 | 2 | 0.00518 | 0.37663 | 140.6768 | 0.20902 | 0.08569 | 139.62 | 104 | 80.04569 | 0.59805 | 0.00151 | 0.07626 |
| 2 | sbm | sbm | 450 | 61149 | 2 | 1.4957 | 4174.373 | 0.26853 | 2 | 0.00404 | 0.61845 | 271.7737 | 0.17899 | 0.09295 | 271.7733 | 194 | 192.2356 | 0.75807 | 0.00142 | 0.16225 |
| 3 | sbm | sbm | 450 | 32272 | 2 | 1.69406 | 3599.436 | 0.26853 | 2 | 0.00601 | 0.53334 | 157.437 | 0.17899 | 0.09295 | 143.4311 | 132 | 58.8829 | 0.56401 | 0.00146 | 0.04293 |
| 4 | sbm | sbm | 450 | 34684 | 2 | 1.66466 | 3362.436 | 0.22509 | 2 | 0.00051 | 0.38456 | 161.037 | 0.18195 | 0.08686 | 155.0578 | 119 | 88.08883 | 0.50828 | 0.00134 | 0.18727 |
| 5 | sbm | sbm | 450 | 30566 | 2 | 1.69445 | 3454.927 | 0.17176 | 2 | 0.00158 | 0.35337 | 150.997 | 0.25502 | 0.16199 | 136.2933 | 97 | 71.5599 | 0.56401 | 0.00131 | 0.15738 |
| 6 | sbm | sbm | 450 | 30661 | 2 | 1.69650 | 3690.476 | 0.17517 | 2 | 0.00111 | 0.37388 | 138.3115 | 0.26223 | 0.09792 | 136.2711 | 103 | 80.27211 | 0.61678 | 0.00131 | 0.15836 |
| 2 | sbm | sbm | 500 | 67387 | 2 | 1.53989 | 4379.472 | 0.29682 | 2 | 0.00484 | 0.64562 | 233.3249 | 0.25208 | 0.08082 | 269.548 | 179 | 103.1858 | 0.45624 | 0.00130 | 0.53107 |
| 3 | sbm | sbm | 500 | 64458 | 2 | 1.50346 | 4676 | 0.18956 | 2 | 0.00404 | 0.64229 | 219.4751 | 0.15090 | 0.08473 | 217.832 | 171 | 131.1796 | 0.63514 | 0.00128 | 0.20162 |
| 4 | sbm | sbm | 500 | 40310 | 2 | 1.69127 | 4317.614 | 0.21814 | 2 | 0.00057 | 0.42905 | 198.5435 | 0.31478 | 0.09771 | 161.24 | 160 | 121.9872 | 0.72479 | 0.00129 | 0.10273 |
| 5 | sbm | sbm | 500 | 45695 | 2 | 1.63370 | 4545.513 | 0.19187 | 2 | 0.00651 | 0.39253 | 184.4573 | 0.18907 | 0.08901 | 182.78 | 134 | 102.8226 | 0.61952 | 0.00122 | 0.17685 |
| 6 | sbm | sbm | 500 | 42515 | 2 | 1.65918 | 4464.11 | 0.17176 | 2 | 0.00506 | 0.39681 | 172.6992 | 0.27478 | 0.10096 | 170.06 | 128 | 96.27144 | 0.61952 | 0.00120 | 0.15117 |
| 2 | sbm | sbm | 550 | 74389 | 2 | 1.50727 | 5396.507 | 0.19518 | 2 | 0.00202 | 0.53123 | 271.1488 | 0.10280 | 0.05008 | 270.5055 | 238 | 162.5021 | 0.60468 | 0.00120 | 0.09262 |
| 3 | sbm | sbm | 550 | 63317 | 2 | 1.58081 | 5346.286 | 0.17393 | 2 | 0.00578 | 0.45644 | 235.6225 | 0.33417 | 0.11795 | 230.2436 | 162 | 139.5227 | 0.57248 | 0.00115 | 0.17137 |
| 4 | sbm | sbm | 550 | 53324 | 2 | 1.64680 | 5013.17 | 0.23277 | 2 | 0.00714 | 0.41564 | 197.1511 | 0.21585 | 0.08704 | 193.9055 | 135 | 101.7535 | 0.53647 | 0.00110 | 0.16411 |
| 5 | sbm | sbm | 550 | 50590 | 2 | 1.66481 | 5045.142 | 0.21973 | 2 | 0.00922 | 0.38088 | 186.2647 | 0.23573 | 0.08385 | 184.8727 | 143 | 100.2512 | 0.57248 | 0.00109 | 0.15117 |
| 6 | sbm | sbm | 550 | 48394 | 2 | 1.67945 | 5008.973 | 0.18624 | 2 | 0.00729 | 0.35969 | 179.3593 | 0.24548 | 0.09206 | 175.9782 | 112 | 99.58619 | 0.63045 | 0.00108 | 0.13913 |
| 2 | sbm | sbm | 600 | 73620 | 2 | 1.59031 | 5188.958 | 0.27273 | 2 | 0.00494 | 0.58897 | 248.3533 | 0.25502 | 0.06849 | 245.4 | 205 | 61.20324 | 0.25609 | 0.00105 | 0.45302 |
| 3 | sbm | sbm | 600 | 85559 | 2 | 1.52937 | 6252.233 | 0.14752 | 2 | 0.00254 | 0.84063 | 286.4461 | 0.12777 | 0.06478 | 285.1967 | 237 | 205.7878 | 0.56667 | 0.00109 | 0.08309 |
| 4 | sbm | sbm | 600 | 70175 | 2 | 1.60948 | 5857.016 | 0.20252 | 2 | 0.00064 | 0.42927 | 239.3818 | 0.29542 | 0.11533 | 233.9167 | 166 | 119.693 | 0.54893 | 0.00103 | 0.17137 |
| 5 | sbm | sbm | 600 | 65272 | 2 | 1.63677 | 5851.797 | 0.19079 | 2 | 0.00486 | 0.40246 | 221.5357 | 0.18811 | 0.09176 | 217.5733 | 147 | 121.5281 | 0.64337 | 0.00102 | 0.21702 |
| 6 | sbm | sbm | 600 | 58562 | 2 | 1.67411 | 5570.41 | 0.20477 | 2 | 0.00512 | 0.36485 | 198.2843 | 0.25024 | 0.07111 | 195.2067 | 127 | 113.6776 | 0.64337 | 0.00099 | 0.14408 |
| 2 | sbm | sbm | 650 | 95488 | 2 | 1.54728 | 5983.414 | 0.33853 | 2 | 0.00288 | 0.58475 | 294.5371 | 0.13119 | 0.04801 | 293.8092 | 257 | 83.8106 | 0.32045 | 0.00096 | 0.15242 |
| 3 | sbm | sbm | 650 | 77197 | 2 | 1.63400 | 5566.927 | 0.29277 | 2 | 0.00262 | 0.44112 | 251.6294 | 0.22997 | 0.05154 | 237.5292 | 176 | 104.8365 | 0.50828 | 0.00094 | 0.22917 |
| 4 | sbm | sbm | 650 | 75364 | 2 | 1.64289 | 6516.968 | 0.21834 | 2 | 0.00055 | 0.40272 | 235.4475 | 0.25339 | 0.10323 | 231.8892 | 167 | 112.0324 | 0.53394 | 0.00093 | 0.20927 |
| 5 | sbm | sbm | 650 | 69079 | 2 | 1.67296 | 6502.487 | 0.21834 | 2 | 0.00583 | 0.36927 | 213.6209 | 0.22589 | 0.08236 | 212.5508 | 167 | 113.5506 | 0.53394 | 0.00092 | 0.12460 |
| 6 | sbm | sbm | 650 | 68838 | 2 | 1.67506 | 6525.866 | 0.16294 | 2 | 0.00534 | 0.36027 | 213.1001 | 0.20978 | 0.09717 | 210.8862 | 163 | 127.5968 | 0.67628 | 0.00092 | 0.12460 |
| 2 | sbm | sbm | 700 | 101121 | 2 | 1.53177 | 7182.284 | 0.27592 | 2 | 0.00523 | 0.64496 | 328.0482 | 0.25024 | 0.05899 | 289.1171 | 227 | 145.0471 | 0.45624 | 0.00093 | 0.47802 |
| 3 | sbm | sbm | 700 | 91213 | 2 | 1.58255 | 7156.392 | 0.27592 | 2 | 0.00087 | 0.64496 | 272.9452 | 0.25024 | 0.05899 | 260.6086 | 216 | 157.1344 | 0.54893 | 0.00086 | 0.22967 |
| 4 | sbm | sbm | 700 | 94272 | 2 | 1.61466 | 7291.741 | 0.22878 | 2 | 0.00033 | 0.43431 | 272.3753 | 0.20101 | 0.08146 | 269.3485 | 219 | 147.4321 | 0.54893 | 0.00086 | 0.22967 |
| 5 | sbm | SW | 700 | 86251 | 2 | 1.64249 | 7241.537 | 0.21755 | 2 | 0.00081 | 0.39752 | 247.9493 | 0.21584 | 0.06089 | 246.4314 | 200 | 147.4321 | 0.61952 | 0.00086 | 0.24557 |
| 6 | sbm | Spatial | 700 | 77162 | 2 | 1.68460 | 7077.598 | 0.20518 | 2 | 0.00735 | 0.35751 | 224.3445 | 0.26764 | 0.10634 | 220.4629 | 165 | 128.1321 | 0.61994 | 0.00085 | 0.12046 |

| k | Name | Pred | Order | Edges | M.Ecc | M.Path | Energy | Mod | Diam | Betw | Trans | Spec.R | Eig.C | Deg.C | M.Deg | MinCut | Fied | N.Fied | Close | Assort |
|---|---|---|---|---|---|---|---|---|---|---|---|---|---|---|---|---|---|---|---|---|
| 2 | sbm | sbm | 750 | 154528 | 2 | 1.449 | 8290.51 | 0.204 | 2 | 0.00014 | 0.588 | 412.419 | 0.090 | 0.053 | 412.075 | 376 | 242.003 | 0.590 | 0.0009 | 0.0036 |
| 3 | sbm | sbm | 750 | 97925 | 2 | 1.651 | 7562.38 | 0.328 | 2 | 0.0004 | 0.441 | 261.823 | 0.153 | 0.064 | 261.13 | 217 | 109.139 | 0.416 | 0.0008 | 0.077 |
| 4 | sbm | sbm | 750 | 95252 | 2 | 1.66 | 7932.162 | 0.229 | 2 | 0.0004 | 0.389 | 258.298 | 0.2789 | 0.097 | 254.005 | 200 | 135.206 | 0.578 | 0.0008 | 0.297 |
| 5 | sbm | Spatial | 750 | 89243 | 2 | 1.63226 | 7853.635 | 0.23993 | 2 | 0.00297 | 0.36026 | 239.4028 | 0.16265 | 0.03429 | 237.9813 | 195 | 124.1644 | 0.531 | 0.0008 | 0.10838 |
| 6 | sbm | Spatial | 750 | 89274 | 2 | 1.68215 | 7892.371 | 0.16729 | 2 | 0.00458 | 0.34477 | 239.5789 | 0.18572 | 0.07501 | 237.3973 | 184 | 138.4181 | 0.51782 | 0.00079 | 0.36717 |
| 2 | sbm | sbm | 800 | 159947 | 2 | 1.51205 | 9093.686 | 0.22495 | 2 | 0.00185 | 0.53998 | 393.0785 | 0.17125 | 0.07703 | 389.8675 | 323 | 210.9505 | 0.54542 | 0.00083 | 0.35161 |
| 3 | sbm | sbm | 800 | 130464 | 2 | 1.59173 | 8719.964 | 0.25627 | 2 | 0.00255 | 0.45259 | 321.7265 | 0.15147 | 0.06303 | 326.16 | 267 | 178.9541 | 0.52974 | 0.00079 | 0.12291 |
| 4 | sbm | sbm | 800 | 110966 | 2 | 1.65373 | 8570.581 | 0.25567 | 2 | 0.00291 | 0.41182 | 281.6157 | 0.27774 | 0.08048 | 276.67 | 219 | 132.7757 | 0.50896 | 0.00076 | 0.29684 |
| 5 | sbm | sbm | 800 | 103107 | 2 | 1.67113 | 8697.797 | 0.26055 | 2 | 0.00408 | 0.37488 | 266.7554 | 0.32842 | 0.08790 | 257.7675 | 183 | 144.0778 | 0.56583 | 0.00075 | 0.18331 |
| 6 | sbm | sbm | 800 | 104038 | 2 | 1.67427 | 8766.46 | 0.19755 | 2 | 0.00603 | 0.31467 | 264.6096 | 0.28945 | 0.11773 | 260.095 | 209 | 195.0528 | 0.63201 | 0.00075 | 0.15913 |
| 2 | sbm | sbm | 850 | 185170 | 2 | 1.487 | 10526.497 | 0.111 | 2 | 0.0002 | 0.539 | 444.059 | 0.185 | 0.100 | 435.694 | 343 | 327.154 | 0.769 | 0.00079 | 0.209 |
| 3 | sbm | sbm | 850 | 134838 | 2 | 1.62036 | 9317.279 | 0.25152 | 2 | 0.00459 | 0.45169 | 341.6268 | 0.29191 | 0.13939 | 317.2659 | 195 | 114.3469 | 0.44869 | 0.00073 | 0.40717 |
| 4 | sbm | sbm | 850 | 113466 | 2 | 1.68553 | 8692.288 | 0.31152 | 2 | 0.00571 | 0.40446 | 304.6456 | 0.19245 | 0.06985 | 267.0965 | 201 | 121.1854 | 0.47469 | 0.00070 | 0.17679 |
| 5 | sbm | sbm | 850 | 98856 | 2 | 1.72325 | 8740.713 | 0.26350 | 2 | 0.00327 | 0.35972 | 242.8881 | 0.31689 | 0.08392 | 234.8553 | 162 | 119.3339 | 0.47921 | 0.00070 | 0.35887 |
| 6 | sbm | sbm | 850 | 100774 | 2 | 1.72017 | 8644.295 | 0.22214 | 2 | 0.00282 | 0.32765 | 244.0773 | 0.25696 | 0.07452 | 237.1153 | 168 | 130.1162 | 0.52459 | 0.00070 | 0.30115 |
| 2 | sbm | sbm | 900 | 175863 | 2 | 1.56528 | 9335.539 | 0.36527 | 2 | 0.00202 | 0.46114 | 392.8939 | 0.18190 | 0.05472 | 390.8067 | 342 | 103.849 | 0.26369 | 0.00071 | 0.44132 |
| 3 | sbm | sbm | 900 | 132382 | 2 | 1.67279 | 9562.679 | 0.36527 | 2 | 0.00387 | 0.42559 | 298.8454 | 0.14787 | 0.06976 | 294.1822 | 226 | 117.357 | 0.35169 | 0.00067 | 0.43442 |
| 4 | sbm | sbm | 900 | 104599 | 2 | 1.73831 | 9096.162 | 0.37235 | 2 | 0.00387 | 0.37195 | 242.8869 | 0.26691 | 0.06348 | 234.3533 | 177 | 103.4738 | 0.46969 | 0.00064 | 0.42748 |
| 5 | sbm | sbm | 900 | 106337 | 2 | 1.73397 | 9171.951 | 0.30264 | 2 | 0.00346 | 0.34425 | 243.1827 | 0.35169 | 0.07419 | 236.3044 | 194 | 109.8125 | 0.46025 | 0.00064 | 0.40125 |
| 6 | sbm | sbm | 900 | 113326 | 2 | 1.71977 | 9521.951 | 0.30264 | 2 | 0.00184 | 0.32332 | 243.1827 | 0.51712 | 0.03503 | 251.8356 | 158 | 135.4873 | 0.50245 | 0.00065 | 0.30835 |
| 2 | sbm | sbm | 950 | 207862 | 2 | 1.52607 | 10933.67 | 0.26285 | 2 | 0.00184 | 0.48332 | 437.4926 | 0.11712 | 0.03503 | 437.2253 | 304 | 207.0143 | 0.43516 | 0.00069 | 0.30835 |
| 3 | sbm | sbm | 950 | 170924 | 2 | 1.62074 | 11031.84 | 0.23441 | 2 | 0.00307 | 0.40813 | 360.7437 | 0.21845 | 0.05946 | 359.8398 | 306 | 207.0143 | 0.43516 | 0.00065 | 0.30835 |
| 4 | sbm | sbm | 950 | 146338 | 2 | 1.67536 | 11042.7 | 0.23441 | 2 | 0.00307 | 0.38357 | 314.5349 | 0.21845 | 0.05191 | 308.08 | 235 | 136.4081 | 0.41106 | 0.00063 | 0.39115 |
| 5 | sbm | sbm | 950 | 134044 | 2 | 1.72052 | 10886.08 | 0.19012 | 2 | 0.00385 | 0.34526 | 285.8251 | 0.21845 | 0.09677 | 282.1989 | 210 | 150.0146 | 0.41106 | 0.00062 | 0.39115 |
| 6 | sbm | sbm | 950 | 141919 | 2 | 1.68538 | 11256.56 | 0.19012 | 2 | 0.00365 | 0.34785 | 302.8642 | 0.21174 | 0.07219 | 298.5663 | 220 | 177.557 | 0.64594 | 0.00063 | 0.26254 |
| 2 | sbm | sbm | 1000 | 208401 | 2 | 1.52578 | 13313.2 | 0.24215 | 2 | 0.00248 | 0.45881 | 417.5071 | 0.27882 | 0.02325 | 416.802 | 367 | 289.5842 | 0.50993 | 0.00063 | 0.45292 |
| 3 | sbm | sbm | 1000 | 180770 | 2 | 1.63809 | 11836.63 | 0.24214 | 2 | 0.00261 | 0.45585 | 377.101 | 0.27882 | 0.03352 | 361.54 | 249 | 123.9482 | 0.39631 | 0.00061 | 0.56297 |
| 4 | sbm | sbm | 1000 | 174617 | 2 | 1.64848 | 11972.84 | 0.23715 | 2 | 0.00047 | 0.41008 | 360.3718 | 0.21547 | 0.11524 | 349.234 | 254 | 156.9137 | 0.50993 | 0.00061 | 0.06072 |
| 5 | sbm | sbm | 1000 | 172966 | 2 | 1.64562 | 12368.09 | 0.19625 | 2 | 0.00424 | 0.37932 | 346.4108 | 0.14590 | 0.05166 | 345.392 | 292 | 209.3184 | 0.60602 | 0.00060 | 0.06072 |
| 6 | sbm | sbm | 1000 | 161440 | 2 | 1.67679 | 12244.83 | 0.23715 | 2 | 0.00047 | 0.34768 | 326.9368 | 0.21479 | 0.06209 | 322.88 | 227 | 194.4283 | 0.62459 | 0.00060 | 0.06772 |

k = communities; Pred = predicted class; M.Ecc = mean eccentricity; M.Path = mean path length; Mod = modularity; Diam = diameter; Betw = betweenness centrality; Trans = transitivity; Spec.R = spectral radius; Eig.C = eigenvector centrality; Deg.C = degree centrality; M.Deg = mean degree; Fied = Fiedler value; N.Fied = normalized Fiedler value; Close = closeness centrality; Assort = degree assortativity coefficient; SW = small-world

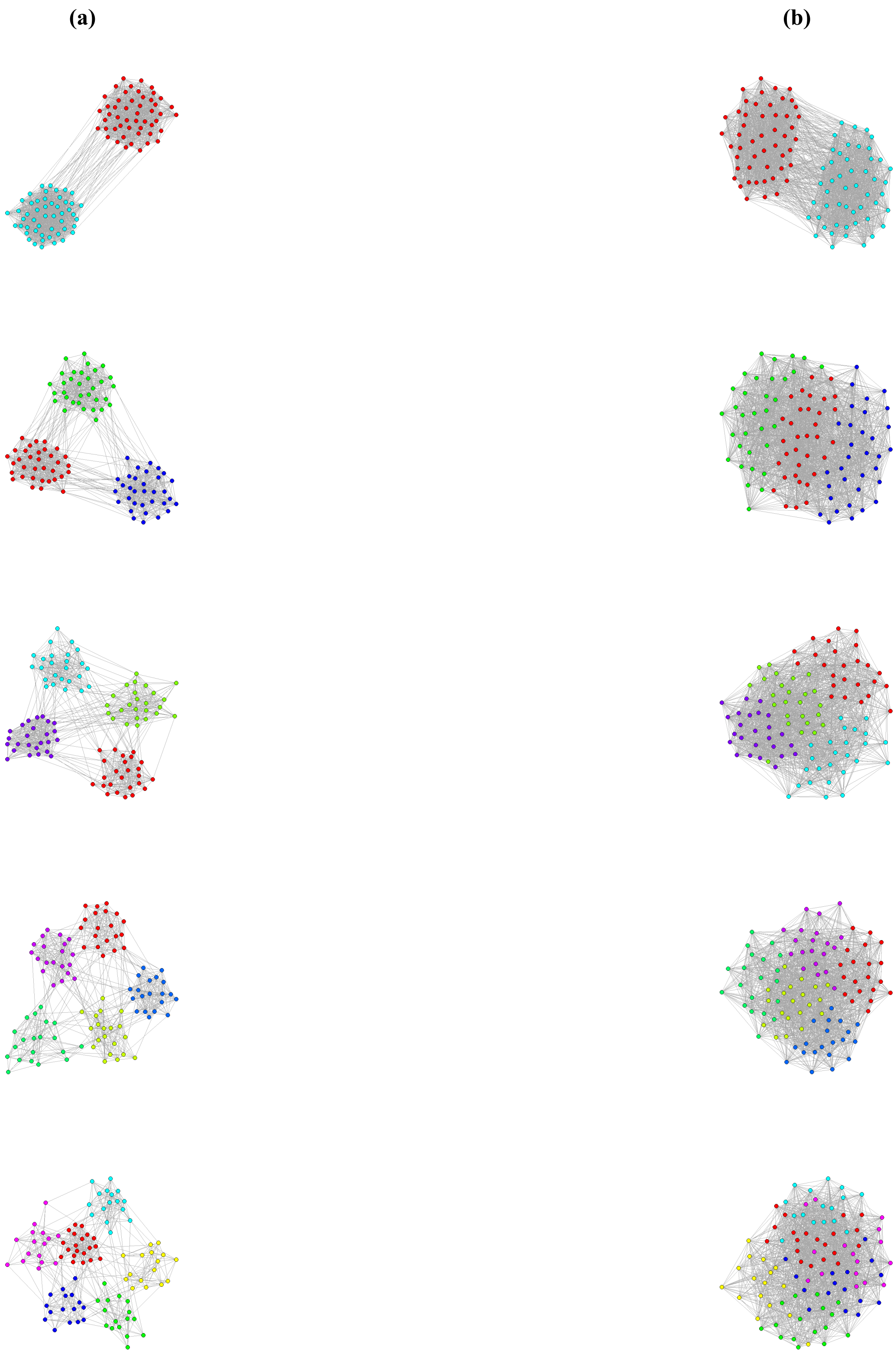


Fig. S11: Network visualizations for different numbers of communities (k=2 to 6) with (a) lower probability of between-community connections ($0.01 \leq p_{between} \leq 0.04$) and (b) higher probability of between-community connections ($0.1 \leq p_{between} \leq 0.4$). All networks shown have order $n = 100$ and within-community connection probability $0.5 \leq p_{within} \leq 0.8$.

### References


D. L. Barack. “What is foraging?” *Biology & Philosophy* 39.1, p. 3. https://doi.org/10.1007/s10539-024-09939-z.

J. A. Bondy. *Graph Theory With Applications*. GBR: Elsevier Science Ltd., p. 264. ISBN: 0444194517.

S. Bonner, J. Brennan, I. Kureshi & A. S. McGough. “Efficient comparison of massive graphs through the use of ’graph fingerprints’”. In: *12th International Workshop on Mining and Learning with Graphs*. Newcastle University.

N. Boyland, R. James, D. Mlynski, J. Madden & D. Croft. “Spatial proximity loggers for recording animal social networks: consequences of inter-logger variation in performance”. *Behavioral Ecology and Sociobiology* 67, pp. 1877–1890. https://doi.org/10.1007/s00265-013-1622-6.

N. Carnegie. “Effects of contact network structure on epidemic transmission trees: implications for data required to estimate network structure”. *Stat Med* 37.2, pp. 236–48. https://doi.org/10.1002/SIM.7259.

M. C. Crofoot, D. I. Rubenstein, A. S. Maiya & T. Y. Berger-Wolf. “Aggression, grooming and group-level cooperation in white-faced capuchins (Cebus capucinus): Insights from social networks”. *American Journal of Primatology* 73.8, pp. 821–833. https://doi.org/10.1002/ajp.20959.

D. P. Croft, J. R. Madden, D. W. Franks & R. James. “Hypothesis testing in animal social networks”. *Trends in ecology & evolution* 26.10, pp. 502–507. https://doi.org/10.1016/j.tree.2011.05.012.

R. E. van Dijk, J. C. Kaden, A. Argüelles-Ticó, D. A. Dawson, T. Burke & B. J. Hatchwell. “Cooperative investment in public goods is kin directed in communal nests of social birds”. *Ecology letters* 17.9, pp. 1141–1148.

L. C. Freeman et al., “Centrality in social networks: Conceptual clarification”. *Social network: critical concepts in sociology. Londres: Routledge* 1, pp. 238–263. https://doi.org/10.1016/0378-8733(78)90021-7.

J. H. Friedman & B. E. Popescu. “Predictive learning via rule ensembles”. *The Annals of Applied Statistics* 2.3, pp. 916–954. https://doi.org/10.1214/07-AOAS148.

F. Harary. *Graph Theory (on Demand Printing Of 02787)*. CRC Press.

T. Kajdanowicz & M. Morzy. “Using graph and vertex entropy to compare empirical graphs with theoretical graph models”. *Entropy* 18.9, p. 320. https://doi.org/10.3390/e18090320.

M. J. Keeling, L. Danon, A. P. Ford, T. House, C. P. Jewell, G. O. Roberts, J. V. Ross & M. C. Vernon. “Networks and the epidemiology of infectious disease”. *Interdisciplinary Perspectives on Infectious Diseases* 2011, p. 284909. https://doi.org/10.1155/2011/284909.

G. Li, M. Semerci, B. Yener & M. Zaki. “Effective graph classification based on topological and label attributes”. *Stat Anal Data Min* 5.4, pp. 265–83. https://doi.org/10.1002/sam.11153.

S. Lundberg, P. Allen & S. Lee. “A Unified Approach to Interpreting Model Predictions”. *cited*. https://doi.org/10.48550/arXiv.1705.0787[illegible]

W. E. Marcílio & D. M. Eler. “From explanations to feature selection: assessing SHAP values as feature selection mechanism”. In: *2020 33rd SIBGRAPI conference on Graphics, Patterns and Images (SIBGRAPI)*. IEEE, pp. 340–347. https://doi.org/10.1109/SIBGRAPI51738.2020.00053.

M. E. J. Newman. “Mixing patterns in networks”. *Phys. Rev. E* 67 (2), p. 026126. https://doi.org/10.1103/PhysRevE.67.026126.

M. E. Newman. “Spectral methods for community detection and graph partitioning”. *Physical Review E* 88.4, p. 042822. https://doi.org/10.1103/PhysRevE.88.042822.

M. Newman. *Networks: An Introduction*. https://doi.org/10.1093/acprof:oso/9780199206650.001.0001.

D. M. Powers. "Evaluation: from precision, recall and F-measure to ROC, informedness, markedness and correlation". *arXiv preprint arXiv:2010.16061*. https://doi.org/10.48550/arXiv.2010.16061.

L. E. Rocha, J. Ryckebusch, K. Schoors & M. Smith. "The scaling of social interactions across animal species". *Scientific Reports* 11.1, p. 12584. https://doi.org/10.1038/s41598-021-92025-1.

P. Sah, S. Leu, P. Cross, P. Hudson & S. Bansal. "Unraveling the disease consequences and mechanisms of modular structure in animal social networks". *Proc Natl Acad Sci* 114.16, pp. 4165–70. https://doi.org/10.1073/pnas.1613616114.

A. J. Seary & W. D. Richards. *Spectral methods for analyzing and visualizing networks: an introduction*. National Academies Press, Washington, DC, pp. 209–228.

M. Shirley & S. Rushton. "The impacts of network topology on disease spread". *Ecol Complex* 2.3, pp. 287–99. https://doi.org/10.1016/j.ecocom.2005.04.005.

D. Shizuka & D. B. McDonald. "A social network perspective on measurements of dominance hierarchies". *Animal Behaviour* 83.4, pp. 925–934. https://doi.org/10.1016/j.anbehav.2012.01.011.

M. Sokolova & G. Lapalme. "A systematic analysis of performance measures for classification tasks". *Information processing & management* 45.4, pp. 427–437. https://doi.org/10.1016/j.ipm.2009.03.002.

M. Stoer & F. Wagner. "A simple min-cut algorithm". *Journal of the ACM (JACM)* 44.4, pp. 585–591. https://doi.org/10.1145/263867.263

E. Štrumbelj & I. Kononenko. "Explaining prediction models and individual predictions with feature contributions". *Knowledge and information systems* 41, pp. 647–665. https://doi.org/10.1007/s10115-013-0679-x.

S. Wasserman & K. Faust. "Social network analysis: Methods and applications". 2, pp. 1–22. https://doi.org/10.1017/CBO9780511815478

P. Wills & F. G. Meyer. "Metrics for graph comparison: a practitioner's guide". *PLoS One* 15.2, e0228728. https://doi.org/10.1371/journal.pone.0228728.

R. Yacouby & D. Axman. "Probabilistic extension of precision, recall, and f1 score for more thorough evaluation of classification models". In: *Proceedings of the first workshop on evaluation and comparison of NLP systems*, pp. 79–91. https://doi.org/10.18653/v1/2020.eval4nlp-1.9.
